# YAP controls cell migration and invasion through a Rho-GTPase switch

**DOI:** 10.1101/602052

**Authors:** Sagar R. Shah, Chunxiao Ren, Nathaniel D. Tippens, JinSeok Park, Ahmed Mohyeldin, Shuyan Wang, Guillermo Vela, Juan C. Martinez-Gutierrez, Seth S. Margolis, Susanne Schmidt, Shuli Xia, Alfredo Quiñones-Hinojosa, Andre Levchenko

**Affiliations:** Department of Biomedical Engineering, The Johns Hopkins University School of Medicine, Baltimore, MD 21231, USA; Department of Neurosurgery, The Johns Hopkins University School of Medicine, Baltimore, MD 21231, USA; Department of Biological Chemistry, The Johns Hopkins University School of Medicine, Baltimore, MD 21231, USA; Department of Oncology, The Johns Hopkins University School of Medicine, Baltimore, MD 21231, USA; Department of Neurology, Kennedy Krieger Institute, Baltimore, MD 21231, USA; CRBM-CNRS, University of Montpellier, 1919 route de Mende, 34293 Montpellier, France; Department of Biomedical Engineering and Yale Systems Biology Institute, Yale University, New Haven, CT 06511, USA

## Abstract

Delineating the mechanisms controlling the invasive spread of non-diseased and transformed cells is central to understanding diverse processes including cancer progression. Here, we found that Yes-associated protein (YAP), a central transcriptional regulator implicated in controlling organ and body size, modulated a Rho-GTPase switch that drives cellular migration by transactivating the Rac1-GEF protein TRIO through direct modulation of its intronic enhancer. Additionally, YAP and TRIO may promote invasive behavior through putative crosstalk with STAT3 signaling, a potential downstream target. Although we found this YAP-dependent infiltrative program in many cell types, it was particularly enhanced in a patient-specific manner in the most common malignant brain tumor, glioblastoma (GBM), where hyperactivation of the YAP, TRIO, and STAT3 signatures also conferred poor clinical outcome. Our analysis suggests that the YAP-TRIO-Rho-GTPase signaling network identified in this study is a ubiquitous regulator of invasive cell spread in both physiological and pathological contexts.

## INTRODUCTION

Cell migration is a tightly regulated process integrating multiple extracellular cues through intracellular signaling networks, which in turn defines the extent of cell polarization and the persistence and speed of locomotion. Extensive studies over the past few decades identified the Rho family of small GTPases, including Rac1 and RhoA, as key regulators of these processes. Ongoing analysis of the regulation of these GTPases and their interaction(*1*) suggests that they form a molecular circuit integrating the inputs from multiple other signaling networks. Insight into these mechanistic links can help identify important molecular components, whose activity can determine the outcome of cell migration-dependent events in development and disease. Ultimately, unraveling these converging pathways will clarify the determinants of cell migration, and may offer new therapeutic paradigms against the aberrant invasion seen in many cancers. Targeting migration pathways may be particularly effective against cancer where most deaths are caused by invasive and metastatic cells rather than the primary tumor mass.

Many extracellular cues affecting cell migration also guide decisions such as cellular identity; therefore, it is likely that migratory behaviors are tightly integrated with transcriptional decision networks. Notably, Yes-associated protein (YAP) is known to integrate many inputs (Hippo signaling, E-cadherin, mechanical cues, etc.) to orchestrate diverse cellular functions including growth, apoptosis, and migration, as well as cancer cell invasion and metastasis(*2–12*). From these studies, *YAP* has emerged as a proto-oncogene frequently overexpressed in a variety of cancers, including a particularly aggressive and infiltrative form of brain cancer, glioblastoma (GBM)(*7–9*, *13–19*). However, it is not fully understood how *YAP* expression and activity play a role in the regulation of the molecular networks controlling cell migration and invasion, and whether this role may enhance the progression of aggressive cancers. Here, we demonstrated that YAP bound a distal enhancer region to promote transcription of *TRIO*, a guanine nucleotide exchange factor (GEF) protein that regulates Rho GTPases function(*20*). YAP-mediated TRIO up-regulation activated Rac1, which in turn inhibited RhoA to increase migration speed. In addition, an increase in YAP-mediated TRIO signaling conferred invasive potential, likely involving STAT3 as a downstream target. These findings reveal YAP-TRIO signaling as a central regulator of cell dispersion that promotes motility and invasion through a Rho-GTPase switch. In addition, our work provides new insights into the clinical significance of YAP to the Mesenchymal subtype of GBM and the potential prognostic relevance of the connected YAP, TRIO, and STAT3 signaling signatures in GBM.

## RESULTS

### YAP regulates migratory capacity

To analyze putative mechanisms of YAP-mediated control of migration and invasion, we used primary patient-derived cultures of human glioblastoma (GBM) cells due to their highly aggressive and invasive spread in vivo that may be associated with YAP hyperactivity(*13*, *21–24*). We also tested another commonly used GBM cell line, GBMA172. In addition, we explored whether these YAP-driven migratory and invasive mechanisms were conserved in non-pathological systems using cells from two different tissues of origin: adult human brain astrocytes (NHA) and adult human breast cells (MCF10A).

Using intraoperatively obtained primary GBM tissues from patients, we found that YAP protein expression was indeed elevated and transcriptionally active in a majority of GBMs examined (**Fig. 1A, table S1, fig. S1A-B**). From these primary GBM tissues, we derived primary cultures (JHGBM612, JHGBM640, JHGBM651, JHGBM1A) and confirmed that these cell lines had similarly high YAP protein levels as the primary GBM tissues from which they were derived (**fig. S1C**). To test whether this elevated YAP protein level contributes to changes in cell migration in these GBM cells, we used multiple shRNA constructs to specifically inhibit *YAP* expression (**fig. S1D-F**). We then used this perturbation to investigate the migration of cells plated at low density on laminin- (NHA and GBM cell lines) or collagen-coated (MCF10A) 2D surfaces. We followed the migratory response at 10-minute intervals for 8 hours. Following the shRNA-mediated stable knockdown of *YAP* (shYAP) using two different targeting sequences (shYAP/shYAP-1, shYAP-2), we observed a significant decrease in cell displacement and migration speed when compared to the control non-targeting vector (shCtrl) in all GBM cell lines and non-neoplastic NHA and MCF10A cells (**Fig. 1B-C, fig. S1G-H, Movie S1**). Conversely, stable overexpression of a constitutively active *YAP* (YAP OE) versus control vector (CONT) in GBM (JHGBM612 and JHGBM1A) and NHA cells showed that YAP up-regulation was sufficient to increase migration speed (**Fig. 1D, fig. S1I-M**). To determine whether YAP affects infiltrative cell migration and spread in vivo, we used a murine intracranial xenograft tumor model of human brain cancer(*25*, *26*), which recapitulated the cellular infiltration along the corpus callosum seen in the donor patient tumor (**fig. S1N**). To do this, we injected 150,000 shCtrl or shYAP JHGBM612 cells into the striatum of the right hemisphere (at pre-determined coordinates from the bregma; see Materials and Methods) of immunocompromised mice and quantified the number of human GBM cells infiltrating along the corpus callosum into the left hemisphere 5 weeks post-injection. Firstly, we confirmed the presence of tumor after the xenograft transplantation of GBM cells by performing a standard H&E staining. The images indeed displayed features commonly seen in murine xenograft models of human GBM (**fig. S1O**). More importantly, we observed significantly fewer shYAP cells infiltrating the contralateral hemisphere than shCtrl JHGBM612 cells (**Fig. 1E-F, fig. S1P**). These findings, coupled with our in vitro observations, suggested that YAP could enhance infiltrative cell migration in different micro-environmental contexts.

**Figure 1:**
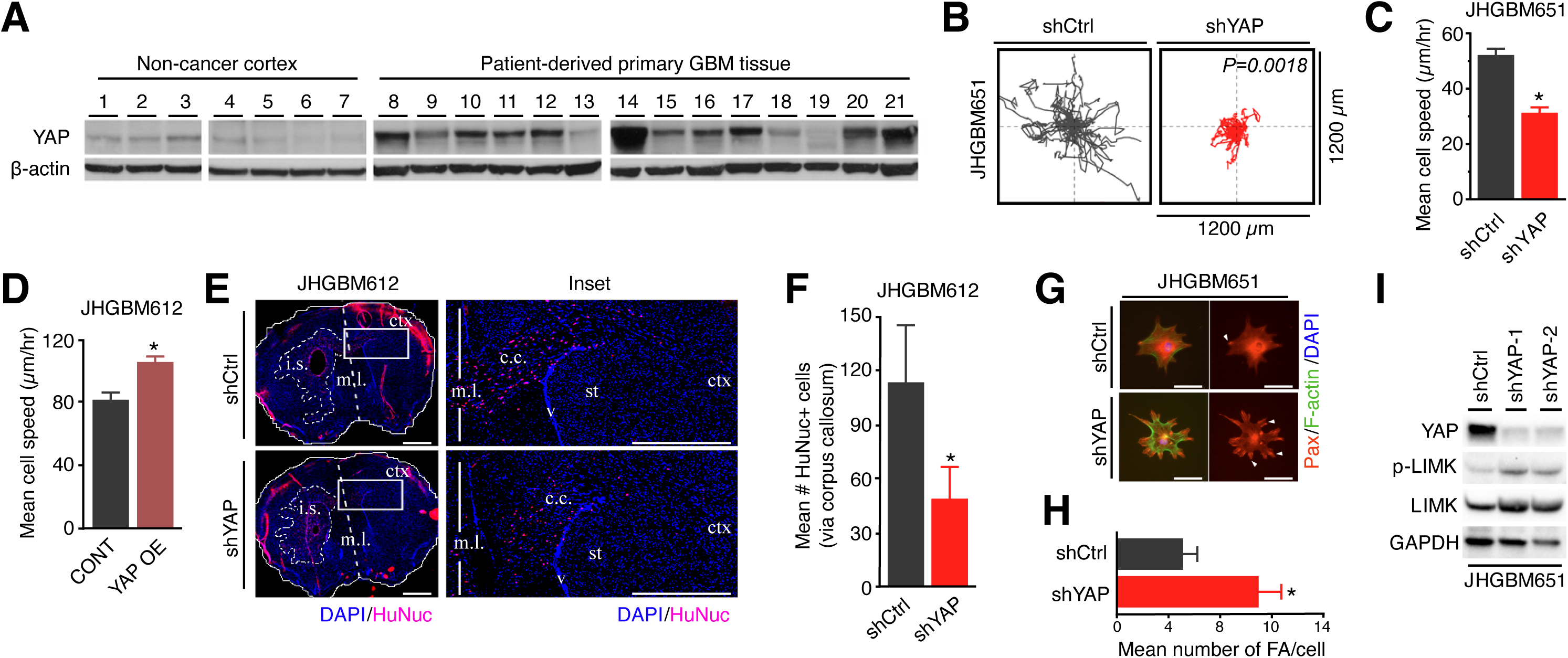
YAP regulates migratory capacity. **(A)** Immunoblots of YAP protein level in lysates from human non-cancer cortex (N=7) and primary glioblastoma patient tissues (N=14). Blots are representative of 3 independent experiments using the same set of patient samples. **(B)** Migration pattern and average distance traveled by shCtrl or shYAP JHGBM651 cells in vitro on 2D surface. Trajectories are calculated from 20 cells each group and are representative of 3 independent experiments. **(C and D)** Mean migration speed of shCtrl versus shYAP JHGBM651 cells (C) or the empty backbone vector transduced cells (CONT) versus *YAP*-overexpressing (YAP OE) JHGBM612 cells (D). Mean migration speed is calculated from 30 cells in each group and are representative of 3 independent experiments; * *P*<0.05, Wilcoxon Rank-Sum test. **(E)** Left: Immunofluorescent staining for human-cell nuclei (HuNuc, red) with an all-nuclei counterstain (DAPI, blue) in mouse brain coronal sections after injection of shCtrl or shYAP JHGBM612 cells (i.s., injection site). Dashed lines outline the bulk tumor margins on the ipsilateral hemisphere. White boxes outline the area of the corpus callosum (c.c.) shown magnified in the right image. Scale bar = 1mm. m.l., midline; st, striatum; ctx, cortex; v, ventricle. Scale bar = 1mm. Images are representative of 5 mice per group. See also fig. S1P. **(F)** Mean number of shCtrl or shYAP cells invading past the midline into the contralateral hemisphere. Quantifications are per mouse and are pooled from 5 mice per group. Student’s t-test; * *P*<0.05. **(G)** Immunofluorescent images of paxillin (PAX, red), F-actin (Phalloidin, green), and cell nuclei (DAPI, blue) in shCtrl and shYAP JHGBM651 cells, representative of 3 independent experiments. Scale bar = 5μm. **(H)** Mean number of focal adhesions (FA) per cell in shCtrl (N=25 cells) and shYAP (N=16 cells) JHGBM651 cells. Results are representative of 3 independent experiments. Student’s t-test; * *P*<0.05. **(I)** Immunoblots of YAP and total and phosphorylated LIMK expression in shCtrl and shYAP JHGBM651 cells, representative of 3 independent experiments. Two independent shYAP constructs were used. In (C-D, F, and H), data shown are mean ± SEM.

We then sought to identify the mechanism by which YAP modulates migratory behavior in non-neoplastic and cancer cells. To do so, we examined the regulation of small Rho GTPases, which are implicated as central regulators of cell motility(*1*). We found that *YAP* knockdown increased the number and area of focal adhesions, strongly affecting the shape of the cells (**Fig. 1G-H, fig. S1Q-R**) in a pattern reminiscent of the focal adhesion assembly phenotype observed upon RhoA hyperactivation(*27*). Consistent with pronounced focal adhesion assembly, an increase in absolute levels and ratio of phosphorylated and total LIM-kinase, a downstream target of Rho-associated protein kinase (ROCK), a major effector of RhoA(*1*), was observed in shYAP JHGBM651, JHGBM612, NHA, and MCF10A cells than their corresponding shCtrl cells (**Fig. 1I, fig. S1S-U**). From these results, we concluded that YAP regulated cytoskeletal organization and dynamics to potentiate cellular motility in vitro and in vivo.

### YAP regulates migration through a Rho-GTPase switch

To better understand the regulation of RhoA by YAP, we measured its GTP load through G-LISA assays. We observed a significant increase in RhoA-GTP levels after *YAP* knockdown in JHGBM651 and NHA cells (**Fig. 2A, fig. S2A**). Based on this data, we sought to test the relevance of RhoA-ROCK activity on cell speed. We found that pharmacologic inhibition of ROCK I/II using either Y27632 or H1152 restored the migration speed of shYAP-1 and -2 cells in a dose-dependent manner while having no effect on shCtrl cells (**Fig. 2B, fig. S2B-L, Movie S2**). Furthermore, treatment with Y12732 or H1152 decreased the phosphorylation of Nonmuscle Myosin II (NMII), confirming the inhibition of ROCK activity (**fig. S2M-N**). These results demonstrated that YAP could modulate RhoA-ROCK activity and that ROCK inhibition was sufficient to rescue the migratory speed of *YAP* knockdown cells. Thus, YAP promoted migration at least in part by suppressing RhoA-ROCK activity.

**Figure 2:**
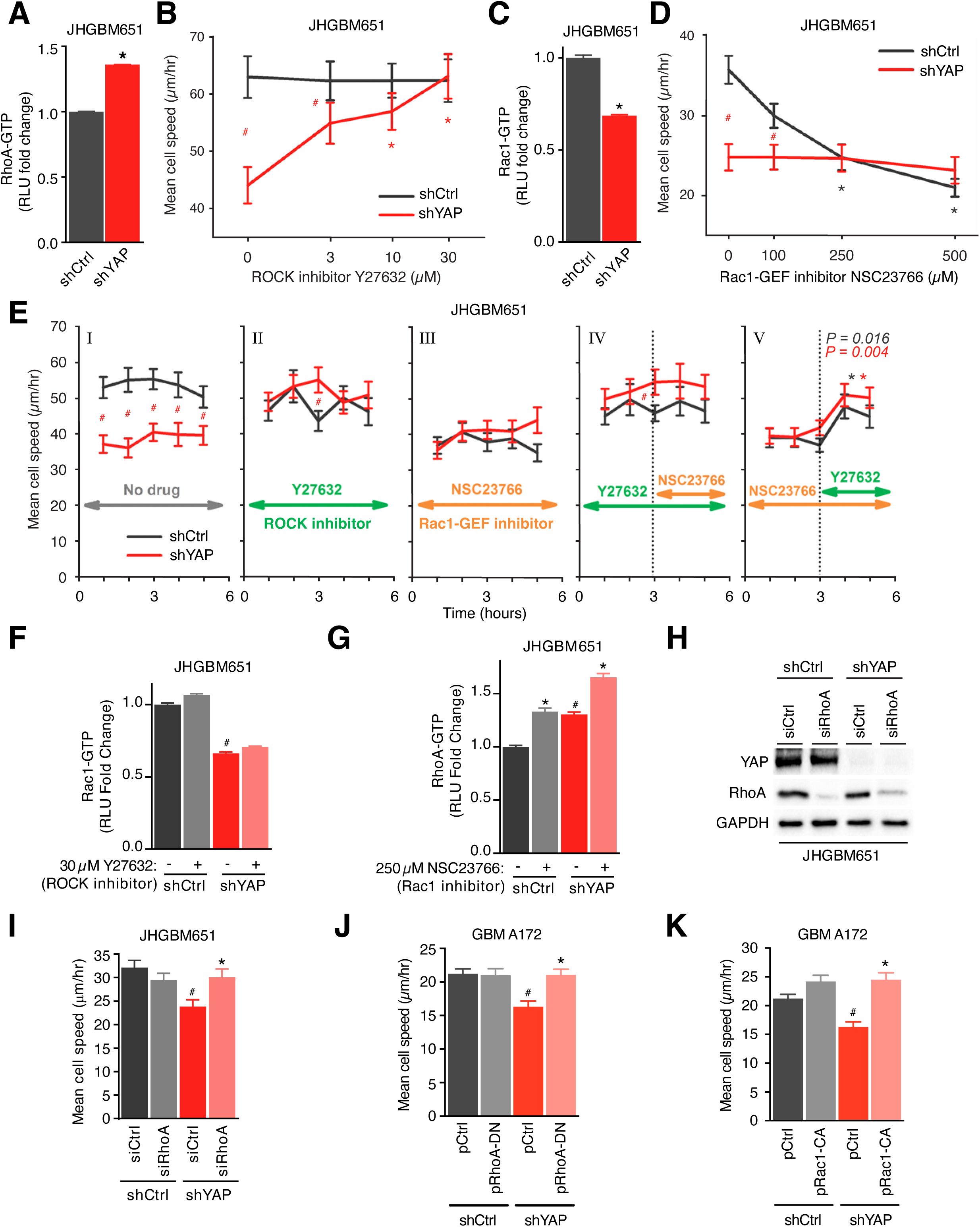
YAP regulates migration through a Rho-GTPase switch. **(A)** G-LISA analysis of RhoA-GTP levels in shCtrl or shYAP JHGBM651 cells pooled from 3 independent experiments. Student’s t-test; * P<0.05, between shCtrl and shYAP cells. **(B)** Mean migration speed of shCtrl or shYAP JHGBM651 cells treated with increasing doses (3, 10, 30µM) of the ROCK inhibitor Y27632. Wilcoxon Rank-Sum test; red * P<0.05, shYAP + Y27632 (10 or 30µM) vs no drug treatment; red # P<0.05, between shCtrl and shYAP cells at the corresponding dose. **(C)** G-LISA analysis of Rac1-GTP levels in shCtrl or shYAP JHGBM651 cells pooled from 3 independent experiments. Analyzed as in (A). **(D)** Mean migration speed of shCtrl or shYAP JHGBM651 cells treated with increasing doses of the Rac1-GEF inhibitor NSC23766. Wilcoxon Rank-Sum test; black * P<0.05, shCtrl + NSC23766 (250 or 500µM) vs shCtrl without drug; red # P<0.05, shCtrl vs shYAP cells of the indicated condition. **(E)** Mean migration speed of shCtrl and shYAP JHGBM651 cells that were unperturbed (“No drug”, first graph) or treated with 10µM Y27632 or 250µM NSC23766 for 3 or 6 hours as indicated. Wilcoxon Rank-Sum test; black *P<0.05, hours 1 to 2 vs hours 4 to 5 in shCtrl cells and in shYAP cells (black and red asterisks with P values, respectively); red # P<0.05, shYAP cells vs shCtrl cells at the marked time points. **(F)** G-LISA analysis of Rac1-GTP levels in shCtrl and shYAP JHGBM651 cells treated with vehicle or 30µM Y27632. Results are pooled from 3 independent experiments. Student’s t-test; * P<0.05, between drug and no drug treatment in shCtrl or shYAP cells; # P<0.05, between shYAP and shCtrl cells with no drug treatment. **(G)** G-LISA analysis of RhoA-GTP levels in shCtrl or shYAP JHGBM651 cells treated with vehicle or 250µM NSC23766. Results are pooled from 3 independent experiments. Analyzed as in (F). **(H)** Immunoblot of YAP and RhoA expression in shCtrl or shYAP JHGBM651 cells transfected with siCtrl or siRhoA, representative of 3 independent experiments. **(I)** Mean migration speed of shCtrl or shYAP JHGBM651 cells transfected with siCtrl or siRhoA. Wilcoxon Rank-Sum test; * P<0.05, between conditions in shYAP cells; # P<0.05, between the siCtrl conditions. **(J)** Mean migration speed of shCtrl or shYAP GBMA172 cells transfected with the empty backbone vector (pCtrl) or a dominant-negative RhoA vector (pRhoA-DN). Wilcoxon Rank-Sum test; * P<0.05, between pRhoA-DN and pCtrl in shYAP cells; # P<0.05, between pCtrl in shYAP and that in shCtrl cells. **(K)** Mean migration speed of shCtrl and shYAP GBMA172 cells transfected with the empty backbone vector (pCtrl) or a constitutively active Rac1 vector (pRac1-CA). Wilcoxon Rank-Sum test; * P<0.05, between the conditions in shYAP cells; # P<0.05, between the pCtrl conditions. In (B, D, E, and I-K), mean migration speed is calculated from 30 cells [from a representative one of / pooled from] X independent experiments. In (A-G, and I-K), the data shown are mean ± SEM.

YAP can modulate the activity of RhoA either directly or indirectly. For instance, prior studies suggest that another small Rho-GTPase, Rac1, can antagonistically regulate RhoA(*28–30*), raising the possibility that modulation of RhoA activity by YAP might involve Rac1. We indeed found that Rac1-GTP levels were reduced after *YAP* knockdown (**Fig. 2C, fig. S3A**). We also examined a downstream target of Rac1, p21-activated kinase (PAK), and found a pronounced decrease in phosphorylated PAK (p-PAK) level after normalizing to the total PAK protein level in shYAP cells compared to shCtrl cells (**fig. S3B**). Thus, we inquired whether changes in Rac1 activity accounted for the decreased migratory speed of shYAP cells. We found that a pharmacological inhibitor of Rac1-GEFs, NSC23766, significantly decreased the migration speed of shCtrl cells in a dose-dependent manner, but not that of shYAP-1 or -2 cells (**Fig 2D, fig. S3C-I, Movie S3**). Collectively, these findings demonstrated that YAP regulated the small Rho-GTPase network governing cellular polarity and migration, leading to the activation of Rac1 and inhibition of RhoA in normal and cancer cells **(fig. S3J**).

We next sought to clarify the precise regulatory mechanisms of YAP’s interaction with Rac1 and RhoA given multiple scenarios of mutual regulation between the components. Our results suggested three possible hypotheses: 1) YAP activates Rac1 to inhibit RhoA, 2) YAP inhibits RhoA to activate Rac1, or 3) YAP independently modulates both Rac1 and RhoA. To test these hypotheses, we applied the logic of epistatic analysis by using pharmacological inhibitors of these small Rho-GTPases in lieu of dominant negative mutants. More specifically, we used sequential perturbation of the network components to deduce the regulatory hierarchy (further details and logic of the experiment are presented in **fig. S4A**). Unlike the experiments presented above (**Fig. 2, B and D**) which explored the dose dependence of the inhibitors on shCtrl and shYAP cells, our experiment here explored the time dependence of cell responses to either single or sequential inhibitor application at fixed concentrations, high enough to achieve the maximal effect (10μM Y27632 or 250μM NSC23766). In addition, the initiation time of the experiment was after the first inhibitor being used was already present long enough for the inhibitory effect to be achieved (1 hour). Thereafter, the migration speed of the cells could only change if the application of the second inhibitor has an added effect. Otherwise, the migratory speed should remain unchanged over time. Using this logic, we first verified that the migration speeds of shCtrl and shYAP-1or -2 cells were stable over a 6-hour period with or without treatment using Y27632 or NSC23766 (**Fig. 2E; fig. S4B-C**). These results were consistent with our prior observations in **Fig. 2B and 2D**. Next, we used one or both pharmacological inhibitors of Rac1 and ROCK to determine how these signaling intermediates regulated each other in these cells. Specifically, we treated shCtrl and shYAP-1 or shYAP-2 cells with either the Rac1-GEF inhibitor or the ROCK inhibitor for 3 hours, followed by the addition of the other inhibitor for the next 3 hours (**Fig. 2E; fig. S4B-C**). The order of inhibitor treatments was indispensable in this analysis to ascertain which one of the aforementioned hypotheses is valid (**fig. S4A**). Notably, we found that in cells treated to inhibit ROCK, subsequent inhibition of Rac1-GEF did not further alter the migration speed in either shCtrl or shYAP cells (**Fig. 2E, fig. S4B-C**). However, ROCK inhibition after the initial Rac1-GEF inhibition rescued the cell migration speed of both shCtrl and shYAP cells (**Fig. 2E, fig. S4B-C**). Altogether, these results strongly favored the model (Hypothesis 1 in **fig. S4A**) in which YAP activated Rac1 to inhibit ROCK signaling, resulting in the enhancement of cell migration.

We then investigated the effect of Rac1-GEF or ROCK inhibition on the activity of RhoA and Rac1 in shCtrl and shYAP cells. Our model predicts that Rac1 activity is not dependent on RhoA-ROCK signaling, and indeed we observed no change in Rac1-GTP or RhoA-GTP levels upon ROCK inhibition (**Fig. 2F, fig. S4D-G**). Again, consistent with our model, we observed a significant increase in RhoA-GTP levels in both shCtrl and shYAP cells following Rac1 inhibition with NSC23766 (**Fig. 2G, fig. S4H**). As expected, Rac1 inhibition also significantly reduced Rac1-GTP levels in both shCtrl and shYAP cells (**fig. S4I**). Finally, we tested whether RhoA perturbation was sufficient to restore the migratory speed of shYAP cells. Treatment with two RhoA siRNAs or a dominant-negative RhoA (DN-RhoA) rescued the migratory speed of shYAP cells, with no significant effect on the migration speed of shCtrl cells (**Fig. 2H-J**). Likewise, introducing a constitutively active Rac1 construct yielded similar results (**Fig. 2K**). These findings demonstrated that YAP activated Rac1, resulting in the inhibition of RhoA-ROCK signaling, which altogether enhanced cellular migration (**fig. S4J**). Thus, YAP simultaneously and sequentially tuned Rac1 and RhoA activities to potentiate cell motility.

### YAP directly transactivates the Rac1-GEF TRIO to modulate Rac1 activity

Next, we sought to determine how the transcriptional co-activator YAP regulates the activity of Rac1. Small GTPases can be regulated by GTPase activating proteins (GAPs) and guanine nucleotide exchange factors (GEFs)(*31*). Because NSC23766 disrupts the interaction of Rac1 with two Rac1-GEFs, TRIO, and TIAM1(*32*), we sought to determine if YAP regulates one or both of these Rac1-GEFs. We observed a decrease in TRIO protein and gene expression after *YAP* knockdown in all cell lines tested using two independent *YAP*-targeting shRNA sequences (**Fig. 3A-B, fig. S5A-G**). Moreover, TRIO protein level increased following *YAP* overexpression both in GBM and MCF10A cells (**Fig. 3C, fig. S5H**). However, we did not observe any changes in *TIAM1* gene expression in shYAP cells compared to shCtrl cells (**fig. S5I**). Given that YAP, a transcriptional co-activator, cooperates with TEAD family transcription factors to directly modulate gene expression(*6*), we examined whether *TRIO* is transcriptionally regulated by YAP and whether YAP is found at known enhancer and/or promoter regions of *TRIO*. Using published chromatin immunoprecipitation (ChIP) sequencing data for YAP and TEAD4 in MDA-MB-231 cells(*33*), we found that both YAP and TEAD4 bound an intronic element of *TRIO* (Site 1: 14265617 to 14266192 base pairs on chromosome 5) that bears histone modification patterns indicative of active enhancers(*34*) (**Fig. 3D**). Specifically, this locus bear enhancer-associated H3K4 monomethylation (H3K4me1) and H3K27 acetylation (H3K27ac), with less promoter-associated H3K4 trimethylation (H3K4me3)(*35*). Notably, this enhancer region is independently reported to physically interact with *TRIO*’s promoter(*36*). In addition, we found a weaker enrichment of YAP-TEAD4 binding at a second, intronic binding site (Site 2: 14268225 to 14268865 bp on chromosome 5) (**Fig. 3D**). We also scanned for the presence of TEAD binding motifs at these two *TRIO* enhancer loci in silico. Using previously reported TEAD motifs (*37*), we observed its presence at both *TRIO* enhancer sites. We also identified an additional TEAD4 consensus motif in the *TRIO* enhancer site 2 according to the JASPAR database(*38*) **(Suppl. File 1)**. By contrast, we found no evidence of YAP or TEAD4 binding at *TIAM1* enhancer and promoter elements. Given these findings, we tested the MDA-MB-231 YAP binding sites by ChIP-PCR on both intronic *TRIO* enhancer sites in GBM cells. Indeed, significant YAP binding was observed at both enhancer sites of *TRIO* (**Fig. 3E, fig. S5J**). To evaluate the functional consequence of this binding, we performed an enhancer luciferase assay using a 150 bp sequence (centered on the peak of YAP binding) paired with the promoter region of *TRIO* in HEK293 cells, with or without overexpression of *YAP*. We found that this sequence increased reporter activity compared to negative control and showed increased activity upon *YAP* overexpression (**Fig. 3F**). To further confirm that YAP only interacts with the enhancer and not the promoter sequence of *TRIO*, we performed an additional control experiment using a 1470 bp promoter sequence of *TRIO* paired with a luciferase reporter, with or without overexpression of *YAP*. Because Zanconato *et al*. previously show that *YAP*’s transcriptional regulation is largely restricted to direct enhancer binding with a few exceptions like *CTGF* where YAP occupies its promoter region, we also included a 612 bp sequence of *CTGF*’s minimal promoter linked to a luciferase reporter as a positive control. Unlike *CTGF*’s promoter, *TRIO*’s full-length promoter sequence did not increase reporter activity and showed no enhancement in activity upon *YAP* overexpression (**fig. S5K**). All these results suggested that YAP directly activated *TRIO* transcription by binding to its intragenic enhancer region.

**Figure 3:**
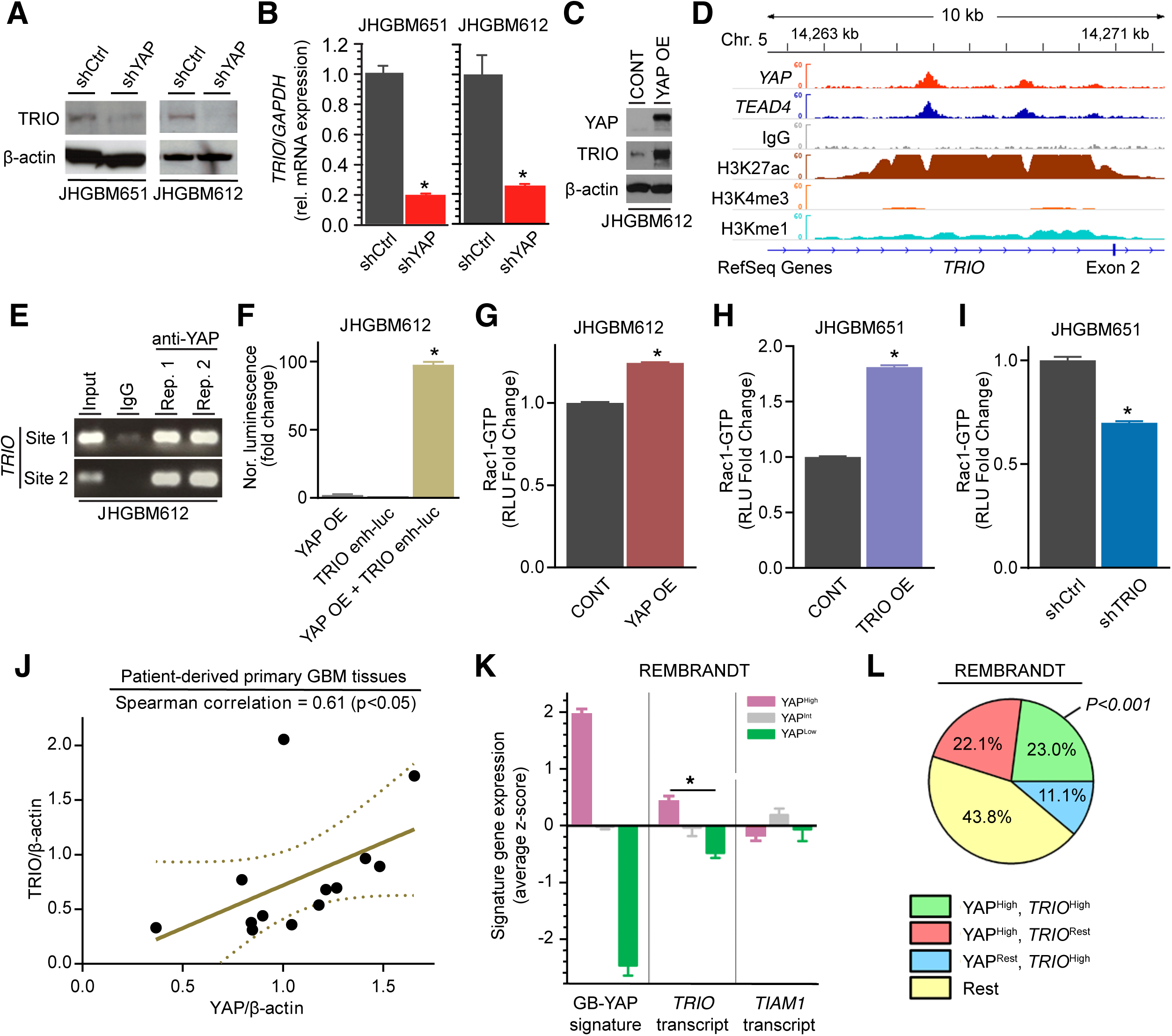
YAP directly transactivates the Rac1-GEF TRIO. **(A)** Immunoblot of TRIO protein level in shCtrl or shYAP JHGBM651/JHGBM612 cells, representative of 3 independent experiments. **(B)** *TRIO* mRNA expression in shCtrl or shYAP JHGBM651/JHGBM612 cells. Results are pooled from 3 independent experiments. Student’s t-test; * P<0.05. **(C)** Immunoblot of TRIO protein level in the empty backbone vector transduced cells (CONT) or YAP OE JHGBM612 cells, representative of 3 independent experiments. **(D)** YAP/TEAD4 binding to an intronic region of *TRIO* as observed in a publicly available ChIP-sequencing dataset using MDA-MD-231 cells (*33*). **(E)** ChIP-PCR querying across two enhancer elements of *TRIO* in JHGBM612 cells. Results are representative of 3 independent experiments. **(F)** *TRIO* gene enhancer luciferase assay using *TRIO* enhancer segment in cells with or without YAP OE in JHGBM612 cells. Student’s t-test; * P<0.05. **(G-I)** G-LISA analysis of Rac1-GTP levels in the empty backbone vector transduced cells (CONT), YAP OE JHGBM612 cells (G), the empty backbone vector transduced cells (CONT), TRIO OE JHGBM612 cells (H) and in shCtrl and shTRIO JHGBM651 cells (I), each from 3 independent experiments. Student’s t-test; * P<0.05. **(J)** Correlation plot of relative YAP and TRIO protein levels (normalized to β-actin) in patient-derived primary GBM tissues resulting from densitometric analysis of immunoblots described in Fig. 6C and fig. S5T. **(K)** *TRIO* and *TIAM1* expression in YAP high, -intermediate, or -low patient groups in the REMBRANDT GBM dataset. Student’s t-test; *P<0.05. **(L)** Percentage of patients with enriched GB-YAP signature, high expression of *TRIO* transcript, or both in the REMBRANDT GBM dataset. P calculated by Fisher’s exact test vs other patient groups. In (B, F-I, and K), data shown are mean ± SEM.

Next, we examined whether YAP-TRIO signaling was sufficient to regulate Rac1 activity. We found an increase in Rac1-GTP levels in YAP OE cells compared to CONT cells (**Fig. 3G, fig. S5L)**. Furthermore, stable overexpression of wildtype *TRIO* (TRIO OE) was sufficient to increase Rac1-GTP levels (**Fig. 3H, fig. S5M)**. Consistent with these findings, shRNA-mediated stable knockdown of *TRIO* (shTRIO) decreased Rac1-GTP levels (**Fig. 3I, fig. S5N-O)**. Given the antagonistic relationship between Rac1 and RhoA signaling, we assessed the activity of RhoA-GTP in these cells. We observed decreased RhoA-GTP levels in YAP OE and TRIO OE cells vs. control cells (**fig. S5P-Q**), and increased RhoA-GTP levels in shTRIO cells vs. shCtrl cells **(fig. S5R)**, paralleling our observations in shYAP cells. Collectively, these results demonstrated that YAP directly transactivated the Rac1-GEF TRIO and that TRIO was necessary and sufficient to activate Rac1 and inhibit RhoA.

Given that there has been evidence suggesting that TRIO-Rac1 signaling can inhibit YAP phosphorylation in HEK293 cells(*39*), we explored this potential feedback regulation in GBM cells. Notably, we did not observe any significant change in either phospho-YAP or YAP protein levels when we knocked down *TRIO* (**fig. S5S**), indicating that no potential direct feedback regulation between YAP and TRIO-Rac1 signaling exists in the context of GBM cells.

To test whether our in vitro observations were consistent with data from clinical samples, we assessed TRIO protein levels in intraoperatively obtained primary GBM tissues from patients (**fig. S5T**). We found that the full-length TRIO protein was indeed expressed in a majority of the GBM tissues analyzed. More importantly, its protein level positively correlated with YAP protein level in these clinical samples (**Fig. 3J**). Next, we surveyed gene expression profiles from the Repository for Molecular Brain Neoplasia Data (REMBRANDT) and The Cancer Genome Atlas (TCGA). Specifically, we explored whether there was a correlation between *YAP* and either *TRIO* or *TIAM1* expression in these two GBM datasets. To enable this analysis, we first derived a list of genes that were significantly downregulated upon stable suppression of *YAP* (by shYAP) compared to controls (shCtrl) in mRNA samples from JHGBM651 and JHGBM612 cells (**table S2**). We referred to these YAP-regulated genes as the GB-YAP signature. Gene signatures allow a more robust estimation of YAP activity from gene expression datasets than mRNA alone, given that many transcriptional regulators, including YAP, undergo substantial post-transcriptional regulation. We found that the GB-YAP signature of GBM cells correlated with YAP signatures derived from other cell types(*40*) (**fig. S5U**). Furthermore, the REMBRANT data suggested that the patients with elevated GB-YAP signature genes also had significantly higher expression of *TRIO*, but not *TIAM1* (**Fig. 3K**). Indeed, patients with *TRIO* overexpression were significantly overrepresented among patients with high GB-YAP signature expression in this dataset (comparing all observed patient group sizes to random expectation using Fisher’s exact test (**Fig. 3L**). Together, these results derived from patient-derived tumor tissues supported our in vitro observations that YAP modulated TRIO expression.

### TRIO is necessary and sufficient for YAP-driven migration

Given our previous results, we sought to test whether TRIO was necessary and/or sufficient for YAP’s control of cell migration. Consistent with its putative functional role, we found that knocking down the expression of *TRIO* decreased the migration speed of GBM, NHA, and MCF10A cells (**Fig. 4A-B, fig. S6A-F, Movie S5**). As a preliminary test, we used a specific pharmacological inhibitor of TRIO, ITX3 to modulate its activity(*41*). We confirmed ITX3 specificity by treating shTRIO cells with this compound and observed no further change in Rac1-GTP levels or migration speed (**fig. S6G-J**). We then found that ITX3 reduced the migration speed of shCtrl cells in a dose-dependent manner without affecting shYAP-1 or -2 cells (**fig. S6K-M, fig. S7A-G, Movie S4**). Furthermore, higher doses of ITX3 also decreased the migration speed of YAP OE GBM cells to match the speed of CONT cells (**fig. S7H)**. Furthermore, ITX3 treatment significantly reduced Rac1-GTP levels in shCtrl and shYAP cells (**fig. S7I**). Conversely, upon treatment with ITX3, a significant increase in RhoA-GTP level was observed in shCtrl, but not in shYAP cells (**fig. S7J)**, demonstrating negative regulation of RhoA activity by TRIO-Rac1 signaling downstream of YAP. These results suggested that the effects of YAP on RhoA-GTPases and cell migration could be entirely explained by TRIO expression.

**Figure 4:**
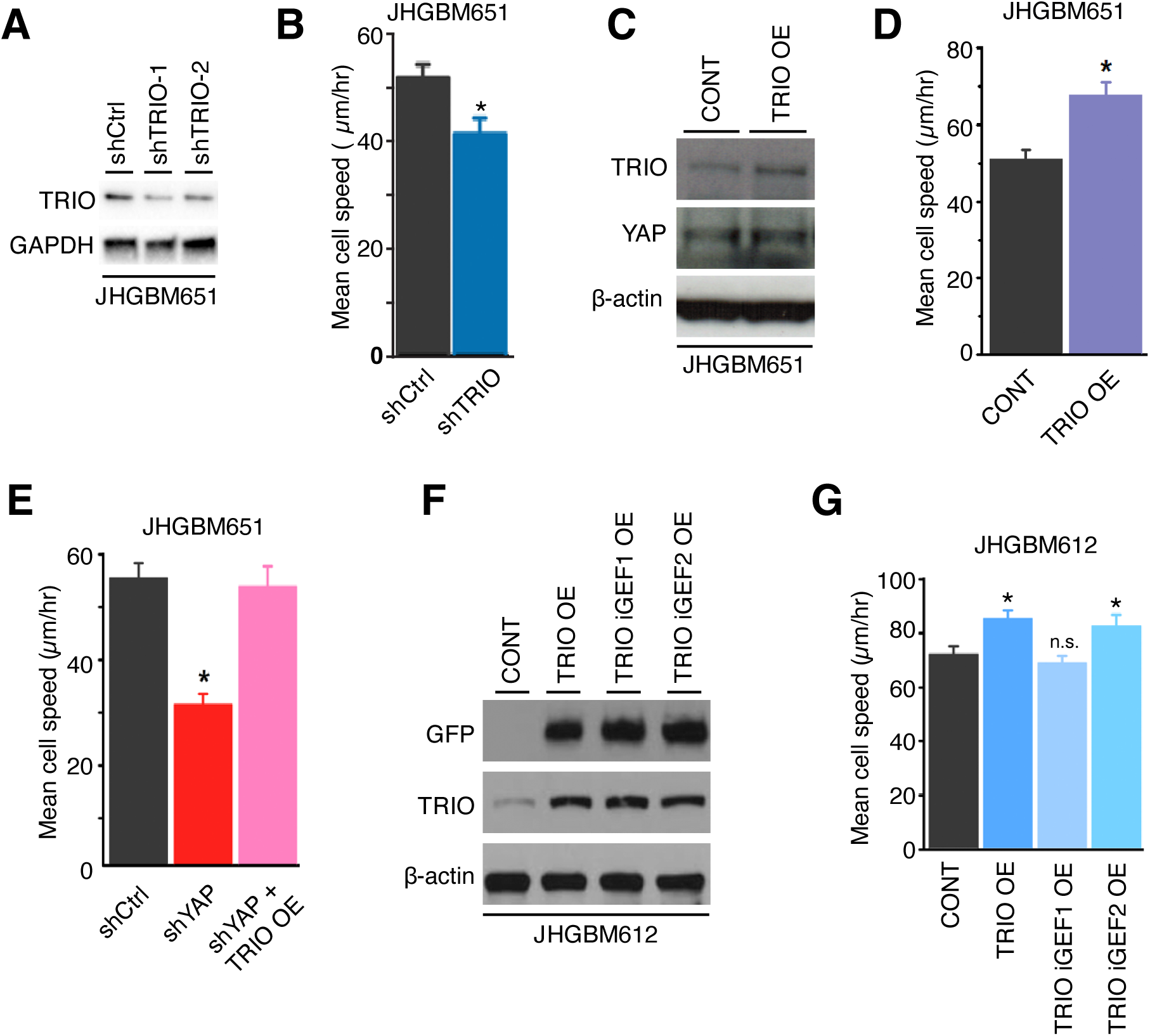
YAP-TRIO signaling regulates migration. **(A)** Representative immunoblot of TRIO protein level in shCtrl and shTRIO JHGBM651 cells with two independent shTRIO constructs from 3 independent experiments. **(B)** Mean migration speed of shCtrl or shTRIO JHGBM651 cells. Wilcoxon Rank-Sum test; *P<0.05. **(C)** Representative immunoblot of TRIO and YAP protein level in the empty backbone vector transduced cells (CONT) and TRIO OE JHGBM651 cells from 3 independent experiments. **(D)** Mean migration speed of the empty backbone vector transduced cells (CONT) and TRIO OE JHGBM651 cells. Wilcoxon Rank-Sum test; * P<0.05. **(E)** Mean migration speed of shCtrl, shYAP, and shYAP+TRIO OE JHGBM651 cells. Wilcoxon Rank-Sum test; * P<0.05 vs shCtrl. **(F)** Immunoblot of GFP and TRIO protein levels in the empty backbone transduced cells (CONT), TRIO–wild-type OE, TRIO-iGEF1 OE, and TRIO-iGEF2 OE JHGBM612 cells, representative of 3 independent experiments. **(G)** Mean migration speed of the empty backbone vector transduced cells (CONT), TRIO OE (wild-type TRIO), TRIO iGEF1 OE, and TRIO iGEF2 OE JHGBM612 cells. Wilcoxon Rank-Sum test; * P<0.05 vs CONT. In (B, D, E, and G), mean migration speed is calculated from 30 cells and results are representative from 3 independent experiments. Data shown is mean ± SEM.

We further explored whether YAP-mediated Rac1 activation was largely driven by TRIO. Given that we found that, in contrast to *TRIO*, the gene expression of *TIAM1* was not reduced in shYAP cells, combined TIAM1/TRIO inhibition using NSC23766 did not significantly alter the migration speed or Rac1-GTP level of shTRIO-1 or -2 cells (**fig. S7K-P**). On the other hand, the introduction of constitutively active Rac1 (Rac1-CA) rescued the migration speed in shTRIO cells but not shCtrl cells (**fig. S7Q**). Furthermore, we found no significant change in RhoA-GTP level in shTRIO cells treated with NSC23766, which contrasted with the increase seen in shCtrl cells treated with the inhibitor compared to vehicle (**fig. S7R**). Additionally, the introduction of constitutively active Rac1 (Rac1-CA) significantly decreased RhoA-GTP levels in shTRIO cells but not in shCtrl cells (**fig. S7S**). These results confirmed the specific role of TRIO but not TIAM1 signaling in mediating YAP’s control of cell migration through activation of Rac1.

We then explored the effect of perturbations of TRIO on RhoA-dependent signaling and its phenotypic consequences. We found that ITX3 treatment significantly increased the RhoA-GTP level of shCtrl, but not shTRIO cells (**fig. S8A**), providing additional evidence of the inhibitory effect of TRIO-dependent signaling on RhoA. Furthermore, the introduction of a dominant-negative RhoA construct (RhoA-DN) rescued the migratory phenotype of shTRIO cells but not of shCtrl cells (**fig. S8B**). Suppression of the downstream effector of RhoA with the ROCK inhibitors, Y27632 or H1152, rescued the migration speed of shTRIO cells, like our observations with shYAP cells (**fig. S8C-H, Movie S6**). Consistent with our prior results, ROCK inhibitors did not significantly affect Rac1-GTP or RhoA-GTP levels in shTRIO cells (**fig. S8I-L)**. However, the introduction of a dominant-negative RhoA construct (RhoA-DN) successfully rescued Rac1-GTP levels in shTRIO cells (**fig. S8M**). These findings suggested that there existed a feedback regulation from RhoA, and not from downstream of RhoA (ROCK), to Rac1, which has been supported by other studies(*30*, *42*, *43*).

Next, we inquired whether TRIO was sufficient to rescue the migration speed of shYAP cells. First, we confirmed that overexpression of *TRIO* increased migratory speed (**Fig. 4C-D, fig. S8N-O**). Remarkably, overexpression of a knockdown-resistant *TRIO* in shYAP-1 or -2 cells (shYAP-1 or -2 + TRIO OE) completely rescued migration speed to levels of shCtrl cells (**Fig. 4E, fig. S8P)**. To further explore the molecular mechanisms of TRIO activity, we focused on its GEF function. TRIO has two distinct GEF domains, GEF1 and GEF2 that control Rac1 and RhoA activation, respectively. However, our findings indicated that TRIO activated Rac1 signaling while inhibiting RhoA to potentiate migration. Thus, we sought to determine whether one or both GEF domains are relevant for this TRIO-mediated migratory behavior seen in GBM cells. We monitored migration speed upon stable overexpression of wildtype TRIO (TRIO OE), TRIO with inactivating GEF1 (TRIO iGEF1 OE) or GEF2 (TRIO iGEF2 OE) mutations (**Fig. 4F-G**). Notably, we observed an increase in migration speed of TRIO iGEF2 cells, phenocopying the migratory behavior of TRIO OE cells (**Fig. 4F-G**). However, we observed no change in migratory speed of TRIO iGEF1 cells (**Fig. 4F-G**). These results demonstrated that the GEF1 domain of TRIO, responsible for activation of Rac1, was indispensable for TRIO’s pro-migratory effect. Moreover, TRIO’s RhoA-activating GEF2 domain appeared dormant and was dispensable for YAP-TRIO-driven migration in GBM cells. Our results demonstrated that TRIO was necessary and sufficient to account for Rac1 activation by YAP, resulting in RhoA inhibition and increased cell migration.

### YAP-TRIO signaling promotes cell invasion

In addition to modulation of migration speed, infiltration of the surrounding stroma requires enhanced invasive capacity, enabling cell navigation in structurally complex tissue environments. In both the TCGA and REMBRANDT tumor gene expression datasets, the GB-YAP signature is associated with higher expression of an invasive gene signature (IGS)(*44*) (**Fig. 5A, fig. S9A**). In a reciprocal fashion, patients with elevated expression of the IGS were significantly overrepresented among patients with high GB-YAP signature expression (**fig. S9B**). Consistent with these observations, we found that fewer shYAP and more YAP OE cells invaded through Matrigel-coated Boyden chambers than their respective control cells in vitro (**Fig. 5B-C, fig. S9C-D**). This led us to explore if the YAP-TRIO signaling is also responsible for the regulation of cell invasion. Indeed, there was a significant decrease in the number of invaded shTRIO cells compared to shCtrl cells, similar to what we saw in shYAP cells (**Fig. 5D, fig. S9E-F**). And there was a significant increase in the number of invaded TRIO OE cells than control cells (**Fig. 5E**). When applying the TRIO inhibitor ITX3, we observed reduced invasion of shCtrl cells but not in shYAP and shTRIO cells after treatment (**fig. S9G-H**). As YAP-TRIO activated Rac1 and inhibited RhoA as illustrated earlier in this study, we also tested the involvement of RhoA in the regulation of cell invasion by applying the ROCK inhibitors, Y26732 and H1152, in shYAP and shTRIO cells. We found that ROCK inhibition rescued the invasive capacity of shYAP and shTRIO cells without significantly altering the invasion of shCtrl cells (**fig. S9I-L**). Given the critical role of YAP-TRIO signaling in driving the migratory and invasive capacity of GBM cells in vitro, we sought to evaluate the relevance of this pathway in vivo using our murine intracranial xenograft model of GBM infiltration. To that end, we established control (CONT) and *YAP* overexpressing (YAP OE) JHGBM612 cells with or without *TRIO* knockdown. Next, we injected 50,000 CONT + shCtrl, CONT + shTRIO, YAP OE + shCtrl, or YAP OE + shTRIO JHGBM612 cells into the striatum (at pre-determined coordinates from the bregma; see Materials and Methods) of immunocompromised mice and quantified the number of human GBM cells infiltrating along the corpus callosum 5-weeks post-injection. We observed that overexpression of *YAP* was sufficient to increase the invasion of JHGBM612 cells through the corpus callosum into the contralateral hemisphere (CONT + shCtrl vs. YAP OE + shCtrl; **Fig. 5F-G**). Moreover, knocking down *TRIO* significantly attenuated the invasive potential of these GBM cells (CONT + shCtrl vs. CONT + shTRIO; **Fig. 5F-G**). In addition, we found a significant reduction in the number of YAP OE + shTRIO cells vs. YAP OE + shCtrl cells migrating through the corpus callosum (**Fig. 5F-G**). Together, these results indicated that whereas YAP was sufficient to increase the invasive capacity of GBM cells, TRIO was necessary to promote YAP-driven cell dispersion in vivo.

**Figure 5:**
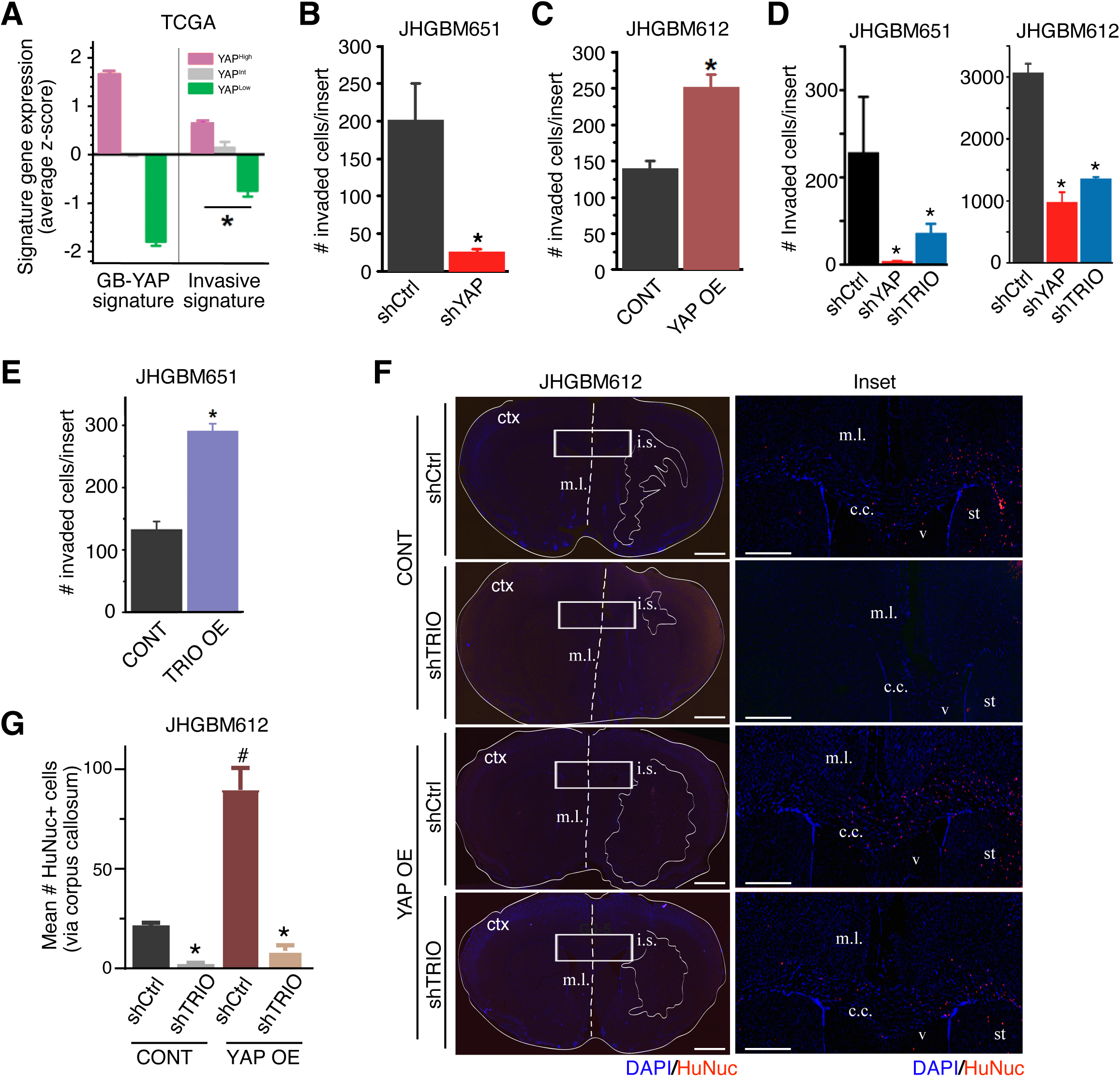
YAP-TRIO signaling promotes cell invasion. (**A)** GB-YAP and Invasive gene signature expression in YAP high, intermediate, or low patient groups in the TCGA GBM dataset. Student’s t-test; * = P<0.05. **(B and C)** Matrigel Boyden invasion assay of shCtrl or shYAP JHGBM651 and the empty backbone vector transduced cells (CONT) or YAP OE JHGBM612 cells, respectively, from 3 independent experiments. Student’s t-test; * P<0.05. **(D)** Matrigel Boyden invasion assay of shCtrl, shYAP, or shTRIO JHGBM651 and JHGBM612 cells from 3 independent experiments. Student’s t-test; * P<0.05. **(E)** Matrigel Boyden invasion assay of the empty backbone vector transduced cells (CONT) and TRIO OE JHGBM651 cells from 3 independent experiments. Student’s t-test; * P<0.05, between CONT and TRIO OE. **(F)** Left: Staining for human nuclei and DAPI in mouse brain coronal sections after injections of the empty backbone vector transduced cells (CONT) + shCtrl, CONT + shTRIO, YAP OE shCtrl, or YAP OE + shTRIO JHGBM612 cells (i.s., injection site). Bulk tumor margins are indicated on the ipsilateral hemisphere. White boxes outline the area of the corpus callosum (c.c.) that is magnified in the right image. Scale bar = 1mm. m.l., midline; st, striatum; ctx, cortex; v, ventricle). Scale bar = 500μm. Images are representative of 5 mice per group. **(G)** Mean number of the empty backbone vector transduced cells (CONT) + shCtrl, CONT + shTRIO, YAP OE + shCtrl, or YAP OE + shTRIO cells invading through the corpus callosum into the contralateral hemisphere. Quantifications are per mouse from 5 mice per group. Student’s t-test; * and # P<0.05 vs shCtrl-CONT cells. In (A-E, and G), data shown are mean ± SEM.

### STAT3 is potentially involved in YAP-mediated cell invasion

Next, we wanted to investigate a potential downstream target of YAP-TRIO signaling that might promote cell invasion. There is evidence suggesting that STAT3, a transcription factor known to promote invasion and metastasis(*45*), can be activated by Rac1(*46*). It is reported that Rac1 can directly bind to and regulate STAT3 activity(*47*, *48*). Rac1 can also indirectly induce STAT3 activity through the autocrine production of cytokine IL-6(*49*, *50*). Various biological processes involve activation of the Rac1-STAT3 signaling such as gland involution(*51*), chondrogenesis(*52*), and primordial follicle pool production(*53*). Additionally, studies have shown that the Rac1-STAT3 axis is important in promoting invasion and epithelial-mesenchymal transition in colorectal cancer(*48*) and gastric cancer(*54*). However, few efforts have been put into exploring this signaling axis in GBM and its potential crosstalk with YAP-TRIO. Notably, we found evidence of an association between elevated expression of the GB-YAP signature and a published GB-STAT3 signature(*55*), with significant overrepresentation of patients with elevated expression of the two signatures in GBM samples from both clinical datasets (**Fig. 6A-B, fig. S10A**). We then surveyed primary GBM tissues for expression and phosphorylation of STAT3 to assess a possible clinical significance of this factor (**Fig. 6C**). We observed that the majority of specimens with elevated YAP protein levels also exhibited high STAT3 phosphorylation levels **(Fig. 6C-D)**. Conversely, only a few tumor samples with low YAP protein levels had high STAT3 phosphorylation **(Fig. 6C-D)**. Given STAT3’s presentation in clinical specimens and datasets, we sought to investigate its connection to the YAP-TRIO-Rac1 network. Firstly, we observed decreased phosphorylation of STAT3 on Tyr^705^ in shYAP-1 or -2 cells (**Fig. 6E, fig. S10B**). Moreover, YAP OE cells expressed higher levels of phosphorylated STAT3 than CONT cells **(Fig. 6F, fig. S10C)**. Functionally, pharmacological inhibition of STAT3 using LLL12 impaired both the invasive and migratory capacity of shCtrl cells more than that of shYAP cells (**Fig. 6G, fig. S10D-F, Movie S7**). Similar to shYAP cells, we observed a decrease in STAT3 phosphorylation following the knocking down of *TRIO* expression vs. shCtrl cells (**Fig. 6H, fig. S10G-H**). Also, applying TRIO inhibitor ITX3 decreased STAT3 phosphorylation levels in a dose-dependent manner (**fig. S10I**). Furthermore, TRIO OE GBM cells exhibited an increase in STAT3 phosphorylation compared to CONT cells (**Fig. 6I**). Remarkably, both stable overexpression of *TRIO* in shYAP cells (shYAP + TRIO OE) and overexpression of *YAP* in shTRIO cells (shTRIO + YAP OE) recovered the phosphorylation levels of STAT3 to shCtrl cell levels (**Fig. 6J-K**). To explore the link between Rac1 and STAT3, we treated the cells with the Rac1-GEF inhibitor, and we observed decreased phospho-STAT3 protein levels in a dose-dependent manner, mirroring our findings with TRIO inhibition (**fig. S10J**). Collectively, these results suggested that STAT3 was involved and could be a potential candidate downstream of the YAP-TRIO-Rac1 axis to drive cell invasion.

**Figure 6:**
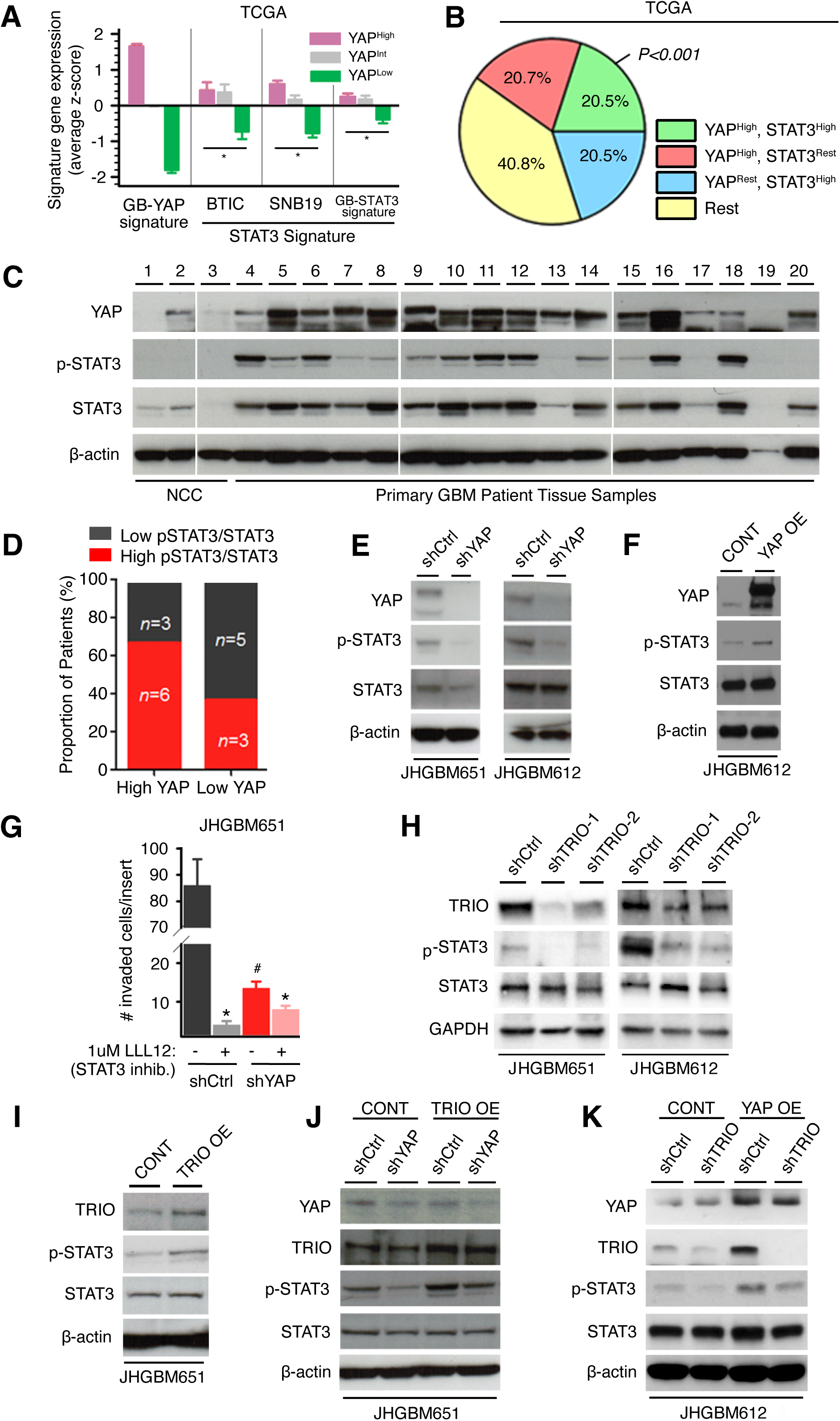
STAT3 is potentially involved in YAP-mediated cell invasion. **(A)** Expression of STAT3 signatures in YAP high, intermediate, or low patient samples in the TCGA GBM dataset. GB-STAT3 signature was derived from conserved STAT3 signatures observed in both BTIC or SNB19 GBM cell lines; Student’s t-test; * P<0.05. **(B)** Percentage of patients with an enriched GB-YAP signature, GB-STAT3 signature, or both in the REMBRANDT GBM dataset. P calculated by Fisher’s exact test vs other patient groups. **(C)** immunoblots of YAP and total and phosphorylated (at Tyr^705^) STAT3 protein levels in tissue homogenates of non-cancer cortices (NCC) and primary glioblastoma specimen, representative of 3 independent experiments using the same set of patient samples. **(D)** Percentage of GBM patients with high (N=9) or low (N=8) YAP protein level that had a high or low Tyr^705^-phsophorylated-to-total STAT3 protein ratio, quantified from immunoblotting analysis described in (C). **(E and F)** Immunoblots of YAP and total and phospho-Tyr^705^ STAT3 abundance in shCtrl or shYAP JHGBM651 and JHGBM612 and CONT or YAP OE JHGBM612 cells, respectively. Results are representative of 3 independent experiments. **(G)** Matrigel Boyden invasion assay of shCtrl and shYAP JHGBM651 cells treated with vehicle or the STAT3 inhibitor LLL12 (1µM). Results are pooled from 3 independent experiments. Student’s t-test; * P<0.05 vs the corresponding no drug (-) treatment; # P<0.05 vs shCtrl (-) cells. **(H)** Immunoblot of total and phospho-Tyr^705^ STAT3 protein levels in shCtrl and shTRIO-1 JHGBM651 and JHGBM612 cells, representative of 3 independent experiments each using two independent shTRIO constructs. **(I)** Immunoblot of total and phospho-Tyr^705^ STAT3 expression in CONT and TRIO OE JHGBM651 cells, representative of 3 independent experiments. **(J)** Immunoblot of total and phospho-Tyr^705^ STAT3 protein levels in shCtrl and shYAP JHGBM651 cells with or without *TRIO* overexpression, representative of 3 independent experiments. **(K)** Immunoblot of total and phospho-Tyr^705^ STAT3 protein levels in shCtrl and shTRIO JHGB612 cells with or without *YAP* overexpression, representative of 3 independent experiments. In (A and G), data shown are mean ± SEM.

### YAP, TRIO, and STAT3 signatures predict poor clinical outcomes in glioblastoma

Our results suggest that YAP-driven migration and invasion are critical for the biology of aggressive cancers. We, therefore, explored the clinical implications of these findings for gliomas, of which GBM is the highest grade (Grade 4) tumor with a median survival of 14 months. We found that GB-YAP signature expression increased with glioma grade in the REMBRANDT dataset (**fig. S11A**). Additionally, the percentage of glioma patients overexpressing the GB-YAP signature compared to non-cancer cortex increased with glioma grade (**Fig. 7A**). Having found that YAP is hyperactive in GBMs, we focused on its prognostic value for these patients as determined by Kaplan-Meier analyses. Strikingly, higher expression of the GB-YAP signature predicted poor prognosis of GBM patients from both TCGA (**Fig. 7B**) and REMBRANDT cohorts (**fig. S11B**). In addition, the *YAP* gene signature was more predictive than 91.5% of 1 million simulated, size-matched gene sets (empirical *p* = 0.085), suggesting it is one of the most predictive gene sets of its size.

**Figure 7:**
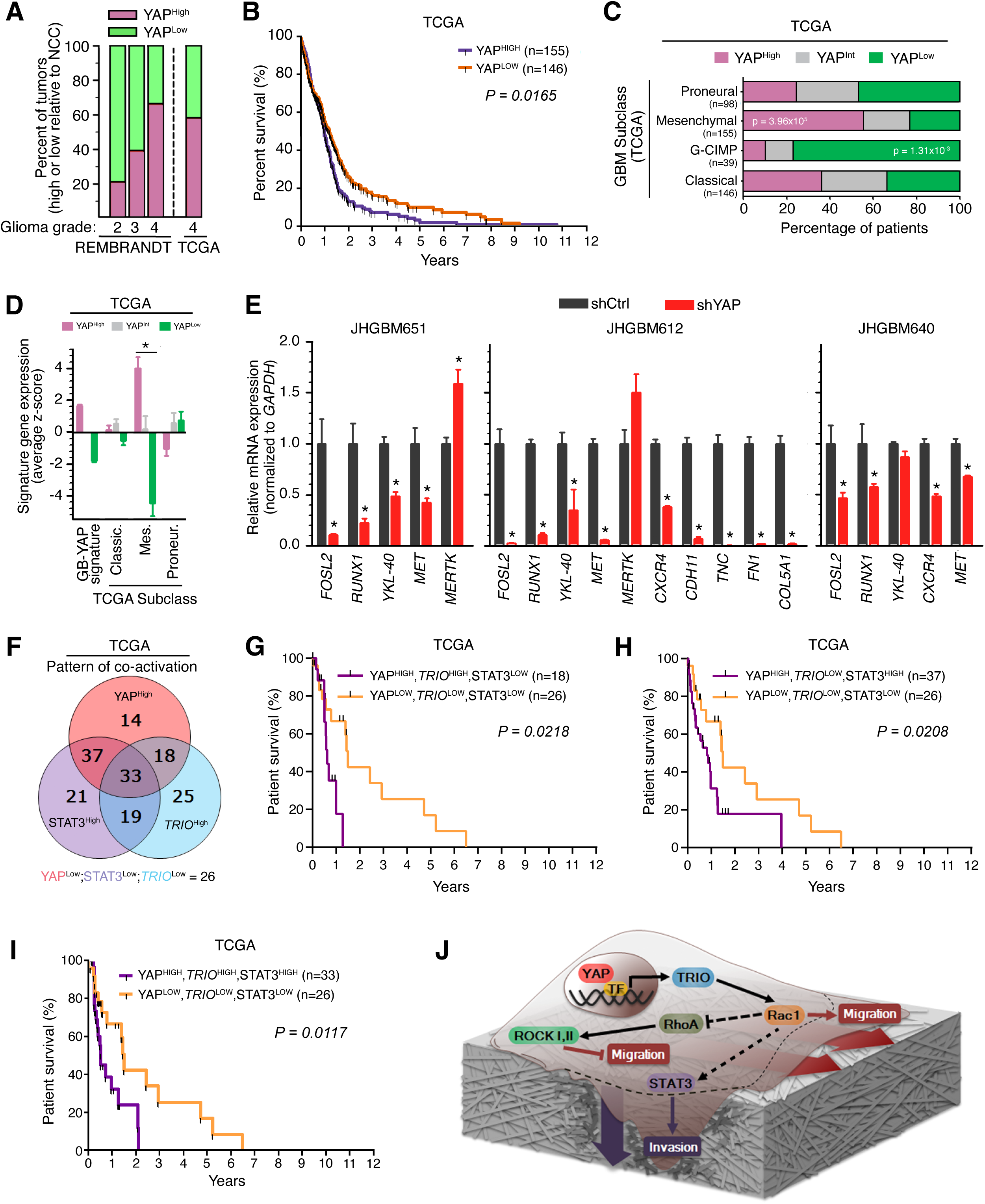
YAP, TRIO, and STAT3 signaling predict poor clinical outcomes in glioblastoma. **(A)** Percentage of patients with an enriched GB-YAP signature expression, grouped by glioma grade, from the REMBRANDT and TCGA datasets. N = 99 grade 2 REM., 84 grade 3 REM., 226 grade 4 REM., and 301 grade 4 TCGA. **(B)** Kaplan-Meier graph of cumulative progression-free survival in patient groups defined by GB-YAP signature expression in TCGA dataset. P calculated by Log-Rank test. **(C)** Percentage of GBM patients with an enriched GB-YAP signature expression, grouped by subtype, from the TCGA. P calculated by Fisher’s exact test; n as noted. **(D)** Expression of GBM YAP high, intermediate, and low patient samples in subclass signatures from the TCGA dataset (as derived from TCGA “Core” samples). Classic., classical, Mes., mesenchymal; Proneur., proneural. Data shown are mean ± SEM. Student’s t-test; * P<0.05, between YAP^High^ and YAP^Low^ within the mesenchymal subset. **(E)** mRNA expression of “Mesenchymal” GBM subclass signature genes in shCtrl or shYAP JHGB651, JHGB640, and JHGB612 cells. Data shown are mean ± SEM. Student’s t-test; * P<0.05 vs corresponding shCtrl. **(F)** Distribution of patient groups expressing either one, two, or all three of the GB-YAP signature, *TRIO* transcripts, and GB-STAT3 signature expression patterns. (**G to I)** Kaplan-Meier graphs of cumulative progression-free survival in patient groups defined by the GB-YAP signature, *TRIO* transcript expression, and GB-STAT3 signature expression in the TCGA dataset. The triple-low group (yellow) is the same in each panel for comparison. P calculated by Log-Rank test. **(J)** Schematic of the YAP-driven pro-migratory and invasive signaling pathway in a single cell on a layer of extracellular matrix. Thick red and purple arrows denote movement of the cell body during migration and invasion, respectively. Darker shading of the extracellular matrix around the invading edge of the cell represents the degradation of the surrounding tissue.

Due to the molecular heterogeneity of GBMs, we explored if YAP activity was associated with one or more of the GBM subtypes including proneural, classical, mesenchymal, and CpG island methylator phenotype (G-CIMP) subtypes described previously(*56*). Notably, Mesenchymal GBMs are distinguished by the invasion of brain parenchyma coupled with pronounced angiogenesis(*57*). We found that there was a significant over-representation of GB-YAP-high patients among the Mesenchymal GBMs but not among other subclasses (**Fig. 7C**). Consistently, higher expression of the GB-YAP signature also corresponded to higher expression of Mesenchymal subclass genes (**Fig. 7D**). In vitro, we observed decreased expression levels of several Mesenchymal subclass-associated genes after *YAP* knockdown in three different GBM cell lines (**Fig. 7E**). On the other hand, we found that GB-YAP-low patients were significantly overrepresented among GBMs in the glioma G-CIMP subclass (**Fig. 7C**), which exhibits hypermethylation at CpG islands and is frequently associated with lower grade gliomas(*58*). To confirm its prognostic value, we evaluated progression-free survival along with GB-YAP signature expression using Kaplan-Meier analyses in each GBM subclass individually. Strikingly, despite its strong association with the Mesenchymal subclass in terms of signature gene expression, higher expression of the GB-YAP signature predicted significantly quicker recurrence in all the GBM subclasses (**fig. S11C-E**).

We then investigated whether incorporating TRIO and STAT3 with GB-YAP signatures would maintain their predictive power considering TRIO and STAT3 expression have also been reported to increase with glioma grade(*59*) similarly to YAP. First, using the TCGA dataset, we determined the distribution of patient groups with hyperactivation of either one, two, or all three of the signaling components (YAP, TRIO, and STAT3). Hyperactivation was determined using the GB-YAP signature, *TRIO* transcript, and GB-STAT3 signature (**Fig. 7F**). Indeed, we found a significant over-representation of GBM patients with hyperactivation of all the three members of *YAP*, *TRIO* and *STAT3*. Next, we stratified patients into 8 groups representing every possible combination of *YAP*, *TRIO*, and *STAT3* co-activation, and performed Kaplan-Meier survival analyses. Notably, the groups that exhibited *YAP* co-activation with either *TRIO* or *STAT3* or both had a significantly worse prognosis than the group without hyperactivation of any of these proteins (YAP^LOW^, TRIO^LOW^, STAT3^LOW^) (**Fig. 7G-I, fig. S11F-G, fig. S12)**. Hyperactivation of *TRIO* or *STAT3* or both without high *YAP* expression did not appear to be associated with significantly worse prognosis in this dataset (**fig. S12**) similarly to the case when high *YAP* expression was combined with low expression level of both *TRIO* and *STAT3*, emphasizing the importance of YAP and also the rather strong interplay among YAP, TRIO and STAT3 in contributing to poor clinical outcome of GBM patients. These results should be viewed as preliminary, as tumor purity, mRNA degradation, and cohort size might be confounding factors in these analyses. Larger patient cohorts with matched expression data would help to validate these results. Nevertheless, collectively, our study provides important initial evidence of the clinical value of the YAP-driven pro-migratory and invasive signaling network in predicting patient outcomes (**Fig. 7J**).

## DISCUSSION

Our findings suggest a central and widespread role for YAP in controlling the migratory speed and invasiveness of cancer and normal cell types through the modulation of a Rho-GTPase switch. Previous work has illustrated the function of cytoplasmic YAP in regulating endothelial cell migration (*60*). Our work uncovered a YAP-mediated signaling network focusing on the role of nuclear YAP as a transcriptional coactivator. Specifically, we demonstrated that YAP increased the expression of the Rac1-GEF *TRIO* by direct modulation of its intronic enhancer, leading to activation of Rac1 and inhibition of RhoA small GTPases. This intricate cascade promoted not only cell motility but also invasive capacity, using both transcriptional and post-transcriptional mechanisms (**Fig. 7J**). Furthermore, we found that, although TRIO harbored two GEF domains, its YAP-driven functions were mediated by the GEF1 domain’s activation of Rac1. This agrees with studies in different systems that demonstrate Rac1 is the major target of TRIO (*20*). One known example of RhoA activation by TRIO is Gα_q_ signaling, where Gα_q_ binds to the PH2 module within the GEF2 domain, releasing PH2-mediated inhibition of DH2, thereby stimulating GEFD2 activity on RhoA (*61*). Nevertheless, the activity of both Rac1 and RhoA was found to be important in mediating the YAP-driven TRIO-dependent control of cell migration and invasion, suggesting the importance of the crosstalk between Rac1 and RhoA, downstream of TRIO activation.

It has been reported that TRIO-mediated Rho-GTPase signaling can regulate YAP (*62–67*) which, coupled with our results, suggests that regulation between YAP and Rho-GTPases can be mutually enhancing, constituting a feedback loop that can stabilize migratory and invasive phenotypes. There have also been other reports suggesting alternative mechanisms of YAP regulating cell motility or Rac1 more specifically. For example, it has been shown that YAP regulates actin dynamics through the modulation of ARHGAP29 to promote metastasis of gastric cancer cells(*68*). Unlike in GBM, YAP modulates Rac1 through transactivation of the Rac1-GEF, TIAM1, through an enhancer-mediated mechanism, during invadopodia formation in breast cancer, supporting the argument for disease-specific regulation(*69*).

In this study, we found that STAT3 was potentially involved in mediating YAP-driven infiltrative spread through an unknown mechanism. Because YAP expression and activity are controlled by a diverse range of stimuli and genetic alterations (*3*, *5*, *7*, *14*, *15*, *63*, *66*, *70*), the evidence of crosstalk between YAP, Rho-GTPases, and STAT3 establishes an even wider spectrum of inputs and mechanisms that regulate cell locomotion. A previous study reports the role of YAP signaling in the activation of the JAK-STAT3 pathway to regulate endothelial cell proliferation during angiogenesis (*71*), further establishing the significance of YAP-mediated activation of STAT3 in different phenotypic and cellular contexts. Likewise, additional studies have shown that STAT3 activation by YAP through other intermediatory regulators can be facilitated by direct interaction. For example, others have shown YAP-TEAD can co-regulate with other partners (such as the AP-1 family transcription factors, including JUNB) on a variety of target sites including STAT3 (*72–74*). Thus, a detailed exploration of YAP-TRIO’s mechanistic connection to STAT3 in the context of Rho-GTPase signaling is warranted. Altogether, these findings support a model of crosstalk between multiple shared signaling components in a cell-type and context-specific manner.

Together with our previous study by Park *et al.* (*75*), our results suggested that YAP-controlled cell migration and invasion were fundamental functions present in many cell types. This pathway may be particularly important in developmental and physiological contexts involving episodes of large-scale migration such as wound healing and progenitor cell navigation during neural tissue development.

Given the widespread hyperactivation of YAP in many cancers (*11*), our study further clarified the mechanisms controlling invasive cancer cell phenotypes. One of the hallmarks of tumor malignancy and progression to higher pathological grades is the ability of cancer cells to invade their surrounding parenchyma, intravasate surrounding blood, and lymphatic vessels, and eventually, seed distant tissues as metastatic tumors (*76*). Tumor recurrence and lethality can be greatly aided by the migratory and invasive capacity of cells (*77*). Whereas the YAP-TEAD interaction has been shown to promote metastasis in breast cancer (*6*), the downstream signals responsible for increased metastatic potential have remained poorly understood. As with metastatic cancers, GBMs often evade eradication because individual cells spread from the primary bulk tumor, thus making complete resection and localized radiation treatment virtually impossible. Clinically, more than half of metastatic tumors display pronounced local infiltration (*24*) and can recur in as little as 3 months despite radical surgery, chemo- and radiotherapy (*21–23*). Thus, understanding the molecular basis for cell dispersal can inform patient prognosis and facilitate the development of improved treatment modalities (*77–80*). Notably, our work highlighted the predictive power of YAP, TRIO, and STAT3 signatures in the prognosis of GBM patients. Lastly, though tumor growth is another important aspect of aggressive cancers, our study was restricted to delineating the role of YAP in invasive spread only. Although YAP has been implicated as a master regulator in cancers whereby its depletion can attenuate tumor growth in multiple cancers—including breast cancer, gastric cancer, and others (*40*, *81*, *82*), very limited efforts have focused on studying the contributions of TRIO in the context of YAP signaling in this aspect (*83*). Intriguingly, in our in vivo invasive spread study, we observed a potential effect on tumor size upon YAP and TRIO modulation. However, our studies did not directly measure tumor growth in a controlled or detailed manner. Thus, further careful examination of the role of YAP-TRIO signaling in cell proliferation and tumor growth would be interesting and clinically informative for the GBM field. Altogether, our study suggests that a network, systems perspective on the etiology of aggressive cancers could benefit from explicit analysis of molecular cascades that integrate numerous signaling pathways, inform the design of targeted and combination therapies, as well as augment our understanding of the drivers of transcriptionally distinct tumor subtypes.

## Materials and Methods

### Cell culture

Patient primary glioblastoma tissue specimens were obtained at the Johns Hopkins Hospital under the approval of the Institutional Review Board (IRB). All primary cell lines were established from excess tumor tissue from patients undergoing surgical resection for glioblastoma as listed in **table S1.** GBM cell line derivation and culture protocols were as previously described(*26*), without the addition of N2 supplement and cultured on laminin-coated (1μg/cm^2^) plates. Specifically, DMEM/F12 (Invitrogen) was supplemented with B27 supplement (Invitrogen), antibiotic/antimycotic (Invitrogen), human EGF (Peprotech), and FGF (Peprotech). MCF10A cells were purchased from ATCC and maintained in complete MEGM media (Lonza) supplemented with 100 ng/mL cholera toxin (Sigma) as recommended. GBMA172 cells were purchased from ATCC and maintained in DMEM (Gibco) supplemented with 10% FBS (Gibco). NHA cells were purchased from Lonza and cultured as recommended. All cell lines were tested and confirmed to be mycoplasm free using a PCR-based MycoDtect^TM^ kit from Greiner Bio-One.

### Lentiviral transduction

Glycerol stocks of Human Mission TRC1 sequence-verified shRNA lentiviral plasmid vectors were obtained from the Johns Hopkins University High Throughput Biology Center (See **table S2**). Plasmids were isolated using a Spin Miniprep kit (Qiagen). Vesicular stomatitis virus glycoprotein-pseudotyped virus was produced by co-transfecting 293T cells using Lipofectamine 2000 (Invitrogen) with an shRNA transducing vector and 2 packaging vectors, psPAX2 and pMD2.G. On days 3 and 4 post-transfection, the virus was harvested and filtered through a 0.22-μm pore cellulose acetate filter before centrifugal concentration using a Centricon Plus-70 (Millipore). An empty TRC1 lentiviral construct was used as the Control virus (Sigma).

The TRIO (pCDH1-GFP TRIO), TRIO iGEF1 (TRIO QALE), TRIO iGEF2 (TRIO L2051E), and its corresponding control lentiviral plasmids (pCDH1) were generated by the Schmidt Lab. Vesicular stomatitis virus glycoprotein-pseudotyped virus was produced by co-transfecting 293T cells using PureFection transfection reagent (Systems Biosciences) with the pCDH1-based lentiviral vector and pPACKH1 packaging plasmid mix (Systems Biosciences).

The *YAP* constitutively active overexpression lentiviral vector was generated using pCMV-Flag YAP127/128/131/381A plasmid (Addgene) and Duet011 lentiviral plasmid.

Primary GBM, GBMA172, MCF10A, and NHA cells were transduced with equal titers of concentrated virus in complete growth media supplemented with 1 μg/ml polybrene (Sigma) for 24 hours. Following transduction, cells were given 24 hours to recover before selection in 0.25 μg/ml (JHGBM651, JHGBM640, JHGBM1A) or 0.5 μg/ml (JHGBM612, GBMA172, MCF10A, and NHA) puromycin (Sigma) for a minimum of 6 days.

### Time-lapse microscopy, drug treatments, and analysis of single-cell migration speed

GBM or MCF10A cells were seeded at a density of 2000-4000 cells/well in a 96-well tissue-culture polystyrene plate (BD Falcon) that was pretreated with 3 μg/cm^2^ laminin for GBM cells (Sigma) or 5 μg/cm^2^ collagen for MCF10A cells (Stem Cell Technologies). 48 hours after seeding, cells were fed with either control or treated complete growth medium (as indicated) and placed and imaged in a temperature- and gas-controlled environmental chamber (37 °C, 5% CO_2_). Y27632 (Tocris), NSC23766 (Tocris), and ITX3 (Tocris) were resuspended in sterile PBS concentrations for the indicated times. Time-lapse microscopy was performed using a motorized inverted microscope (Olympus IX81) equipped with a Cascade 512B II CCD camera. Phase-contrast images were captured with a 10x objective (NA = 0.30) under the control of Slidebook 4.1 (Intelligent Imaging Innovations, Denver, CO) at 10- to 20-min intervals for at least 6 hours. A single observer used a custom MATLAB script to view time-lapse images and manually mark cell body positions (**Movies S1 – S7**). Due to the inhomogeneous seeding of cells throughout individual wells, care was taken to image 5-8 fields of view per well that were most comparable to other experimental conditions. Additionally, a minimum of 30 cells was tracked from 4 replicate wells for each experimental condition. Cell migration speed was computed as displacement (between each captured time point) over time. Individual cell speeds were summarized as the average speed within one-hour intervals. Because the speeds of many cell populations failed to pass normality testing, a Wilcoxon rank-sum test was used to compare experimental conditions. Error bars represent the standard error of the mean of the cell population in a given experimental condition.

### Intracranial xenograft model and immunohistochemistry

All animal protocols were approved by the Johns Hopkins Animal Care and Use Committee (MO09M161). Briefly, JHGBM612 cells were resuspended at 75,000 cells/μl in complete growth media and placed on ice. 1-month-old male NOD-SCID gamma-null (NSG) mice were anesthetized before intracranial injections following a previously reported protocol (*84*). 2 μl of shCtrl or shYAP cells were injected into the striatum; specifically (X: 1.5, Y: 1.34, Z: -3.5) mms from the bregma (n = 5 per group).

5 weeks post-injection, mice were anesthetized before trans-cardiac perfusion with saline followed by 10% formalin. Brains were then harvested and fixed overnight at 4 °C in 10% formalin. Fixed brains were placed in OCT compound (Tissue-Tek) overnight at 4 °C and subsequently flash-frozen. Embedded brains were cut into 10-micron coronal sections using a cryostat microtome.

Brain sections spaced every 500μm were stained with hematoxylin and eosin to identify landmark neuroanatomical structures and therefore identify comparable slides from each mouse for further analysis. Slides immediately adjacent to the identified slides were then selected for immunohistochemistry and quantification. Antigen retrieval was performed using sodium citrate (10 mM) for 30 minutes in a water bath at 95°C. Slides were rinsed in PBS and blocked in 10% normal goat serum for 1 hour. Primary antibody was diluted in PBS with 0.1% Triton X-100 and 2% normal goat serum. Slides were incubated with primary antibody overnight at 4°C. After rinsing with PBS, the slides were incubated with the appropriate anti–IgG secondary antibody (Molecular Probes, Invitrogen) conjugated with a fluorochrome for 1 hour and rinsed. Slides were then incubated with DAPI for 30 minutes and rinsed in PBS before visualization on an Olympus IX81 inverted microscope. Antibodies are listed at the end of this section.

### Immunoblotting, antibodies, and immunocytochemistry

For immunoblotting, tissues from the operating room were briefly rinsed in HBSS before being flash-frozen in liquid nitrogen. Proteins were extracted using T-PER (Pierce) tissue protein extraction reagent and a tissue homogenizer. GBM, MCF10A, and NHA cells were lysed on ice with a cell scraper and radioimmunoprecipitation assay lysis buffer (Pierce) supplemented with protease inhibitor tablets (Roche) and a phosphatase inhibitor cocktail (Pierce). Extracts were incubated for 45 minutes for complete lysis and centrifuged at 10,000 RPM for 10 minutes at 4°C to pellet cell debris. Supernatant protein concentrations were quantified using a BCA Protein Assay kit (Pierce) and a spectrophotometric plate reader (BioTek). Most protein lysates were separated by running 30-100 μg of lysate on 10% Bis-Tris NuPage gels (Invitrogen) and subsequently, transferred to 0.2μm pore polyvinylidene fluoride (PVDF) membranes (BioRad). Because TRIO is an exceptionally large protein, some lysates were run on 4-12% Tris-Acetate NuPage gels and transferred to 0.45 μm pore PVDF membranes overnight. Primary antibody incubations were according to the manufacturer’s recommendations in 0.1% Tween TBS supplemented with 5% non-fat dry milk or BSA, as recommended. Immunoreactive bands were visualized using the appropriate horseradish peroxidase-conjugated anti-IgG antibodies (Pierce). Bands were detected using enhanced chemiluminescence or prime detection reagent (GE Healthcare) whenever appropriate. Densitometric analysis was done with ImageJ. All immunoblotting experiments were performed in triplicate and repeated three times with similar results.

For immunocytochemistry, 160,000 GBM cells were seeded on laminin-coated 35-mm cell culture plates. 48 hours later, cells were washed in cold PBS and fixed in 10% formalin for 30-60 minutes. Cells were blocked in 10% normal goat serum for 1 hour. Primary antibody was diluted in PBS with 0.1% Triton X-100 and 2% normal goat serum. Cells were incubated with primary antibody overnight at 4°C. After rinsing with cold PBS, cells were incubated with the appropriate anti–IgG secondary antibody (Molecular Probes, Invitrogen) conjugated with a fluorochrome for 1 hour. Cells were rinsed in PBS before incubation with DAPI for 30 minutes. Finally, cells were rinsed in PBS before visualization on an Olympus IX81 inverted microscope.

For focal adhesion analysis, high-resolution images were captured using a fluorescence microscope at 20- 40X. Quantification of area or size was conducted using ImageJ.

Antibodies used for these procedures are listed in **table S3**.

### Pull-down of GTP-bound small Rho GTPases

The Rac1-GTP Assay (Cell Bio Labs) and RhoA-GTP Assay (Millipore) kits were used in accordance with the manufacturer’s instructions. Briefly, 1.2 x 10^6^ MCF10A or GBM cells per dish were seeded into collagen- or laminin-coated 100 mm cell culture dishes (Corning). 48 hours later, cells were rinsed twice with ice-cold PBS and placed on ice. Extracts were harvested quickly by scraping in ice-cold 1X assay buffer supplemented with protease and phosphatase inhibitors. Lysis was completed by incubation for 30 minutes on ice and vortexing every 10 minutes. Protein supernatants were isolated and quantified as above. 4 mg of shCtrl or shYAP MCF10A lysate was used as input for RhoA-GTP pull-down and 2 mg as input for Rac1-GTP pull-down. 1 mg of protein lysate was used as input for the Rac1-GTP pull-down of JHGBM651 cells. 50 μg of input lysates were kept for immunoblotting. Agarose beads conjugated with Rhotekin binding domain (RBD) were incubated with lysates for 4 hours, and p21-binding domain (PBD) agarose beads were incubated with lysates for 1 hour. Beads were rinsed three times with cold 1X assay buffer, resuspended in 2X SDS-PAGE sample buffer, and boiled for 10 minutes. Proteins were separated on 10% Bis-Tris NuPage gels, transferred to 0.2 μm pore PVDF membranes, and immunoblotted as described above. Mouse anti-RhoA antibody (26C4) was purchased from Santa Cruz. Refer to the end of this section for a listing of all antibodies used. In addition, for more quantitative analyses, G-LISA RhoA or Rac1 activation kits from Cytoskeleton were used according to the manufacturer’s protocol.

### Motif Search

MotifMap(*85*) was used to search 10 kilobase regions upstream of select transcription start sites in the hg19 reference genome. Significant motifs were called at a false discovery rate below 0.05.

### Invasion assay

Matrigel-coated Boyden chambers (BD Biosciences) were used to assess the invasive capacity of cells. 100,000 cells were resuspended in their appropriate media supplemented with 0.5% FBS. Cell suspension aliquots were plated on the upper surface of the transwell plates with a porous PET membrane (8μm pores) pre-coated with a layer of growth factor reduced matrigel basement membrane matrix. 750 μl of 2% FBS containing DMEM media was placed in the lower chamber. Filters were incubated at 37C°, 5% CO2 for 48 hours. Invasion of cells was determined by fixing the membrane, staining the cells using the Diff-Quik® staining kit, directly counting the number of invaded cells in 9 high-power fields at 10x using an Olympus 1X81 microscope system, and calculating the mean per well. The assays were run in triplicates and at least three independent experiments were performed.

### RNA extraction and quantitative real-time PCR

Total RNA was extracted in TRIzol, phase-separated using chloroform, and precipitated overnight at 4°C in isopropanol. Further isolation of total RNA was performed using the RNeasy Mini Kit (Qiagen) according to the manufacturer’s instructions. 1 μg purified RNA was reverse transcribed using the Superscript III First-Strand cDNA Synthesis Kit (Invitrogen). Quantitative real-time PCR using SYBR Green (Applied Biosystems) was performed on a 7300 Cycler (Applied Biosystems) and the included data analysis software. GAPDH primers were used as a loading control. All primer sequences are listed in **table S4**.

### Biostatistical analysis

Statistical comparisons of experimental results from independent repeats across conditions were performed using unpaired two-tailed Student’s t-test. The comparisons for cell migration data were analyzed using Wilcoxon rank-sum test due to its robustness to outliers. The significance of the association between two categories of patients for example, TCGA mesenchymal subtype and high *YAP* expression were analyzed using Fisher’s exact test. The comparisons of survival data between different patient groups were performed using Kaplan Meier log-rank test. To ensure reproducibility, all in vitro and in vivo experiments were repeated at least three times. The sample sizes, number of replicates and statistical test being used are described in each figure legend.

For Affymetrix data (GB-YAP, TCGA, and REMBRANDT datasets), raw data files were processed in aroma.affymetrix (http://statistics.berkeley.edu/sites/default/files/tech-reports/745.pdf) using a custom CDF with updated probeset mappings(*86*) (BrainArray v15.1.0). Differential expression analysis was performed using the sam function of the siggenes package (version 1.28.0) in R. For Illumina beadchip data, signature probesets (GB-STAT3 and GB-C/EBPβ datasets (*55*)) were required to map to a single Entrez Gene ID. Redundant probesets mapping to the same gene were eliminated after differential expression analysis.

To derive gene signatures, each cell line was analyzed for differential expression independently (GB-YAP, GB-STAT3, and GB-C/EBPβ datasets). Genes with q < .05 and absolute fold-change > 1.5 were considered as differentially expressed genes, although q < 0.1 was used when no genes passed q < 0.05. This relaxed FDR was excused because the final signatures were taken as the genes significantly differentially expressed in both cell lines, at least one of which required q < .05.

To stratify patients into groups of low, intermediate, and high expression of a given gene signature, we used the gene set z-score(*87*). Briefly, RMA normalized data were quantile normalized, variance filtered, and log2 transformed. Gene sets were filtered to remove genes that were not present in the normalized, filtered data for each analysis. Each gene was converted into an expression z-score relative to other samples of the same class (such as other GBMs but not low-grade gliomas), and the gene set score (GSS) was computed as the sum of the member genes’ z-scores divided by the square root of the size of the gene set. Samples with GSS < -0.5 were categorized as low, samples with GSS > 0.5 were high, and those between were intermediate. Because microarray data to derive a gene signature of TRIO was unavailable, patients with a *TRIO* gene z-score < -0.5 were called low and those with a *TRIO* gene z-score > 0.5 were called high. To compare expression level of a specific gene or gene signature between pre-stratified patient groups, the z scores for all the genes in the signature were averaged. The statistical comparison of the averaged z scores between patient groups were performed using unpaired two-tailed Student’s t-test.

## Acknowledgements

We thank the Department of Neurological Surgery and Oncology at Johns Hopkins Hospital for access to intraoperatively-obtained glioblastoma tissues; Lakesha Johnson and Liron Noiman for establishing primary cell cultures of these primary glioblastoma specimens; Colette Ap Rhys for designing and preparing lentiviruses; Conover Talbot Jr. for technical support and helpful discussions of microarray analysis; Tom Schaffer for technical support with the RhoA-GTP pull-down assay; and Kaisorn L. Chaichana for glioblastoma patient MRI images.

## Author contributions

S.R.S. conceived and designed the project. S.R.S., N.D.T. performed the experiments. C.R. assisted with the experiments. N.D.T. performed bioinformatic analysis. J.P. tracked and analyzed migration time-lapse data. A.M. and G.V. assisted with early experiments. J.C.M-G assisted with YAP-TRIO in vitro rescue experiments. S.S. provided TRIO wildtype and mutant lentiviral and corresponding control constructs. S.W., and S. X. assisted with YAP-TRIO in vivo experiment. S.R.S., N.D.T., C.R., and J.P. interpreted the results. A.L. and A.Q.-H. supervised the project. S.R.S. and N.D.T. wrote the manuscript. S.R.S., C.R., S.M., A.L., and A.Q.-H. reviewed and revised the manuscript.

## Data and materials availability

Microarray data of shCtrl and shYAP JHGBM651 and JHGBM612 cells (**Data File S1**) have been deposited with NCBI Gene Expression Omnibus under accession ID GSE289667. All other data needed to evaluate the conclusions in the paper are present in the paper or the Supplementary Materials.

## Supplementary Materials for

**Figure S1:**
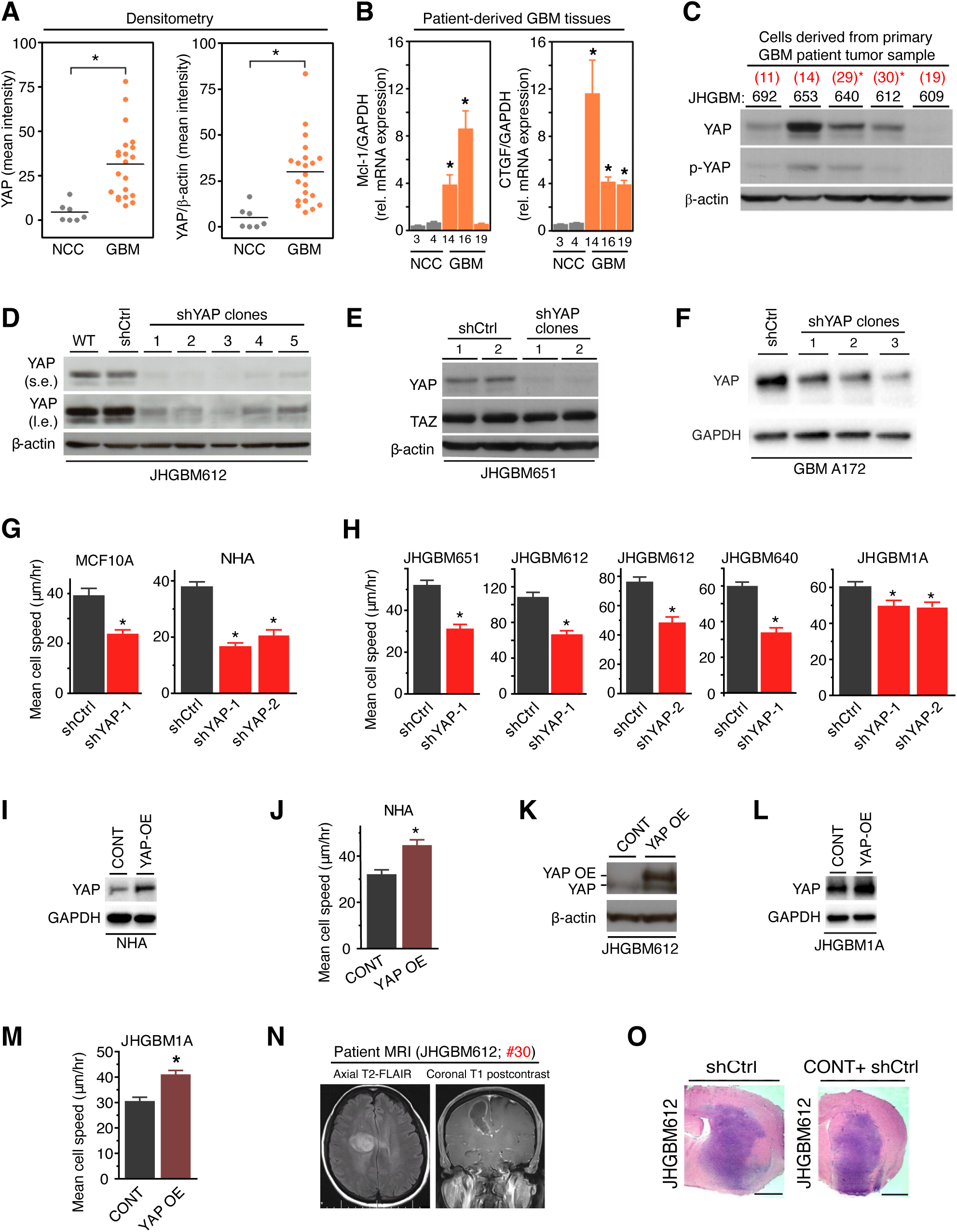
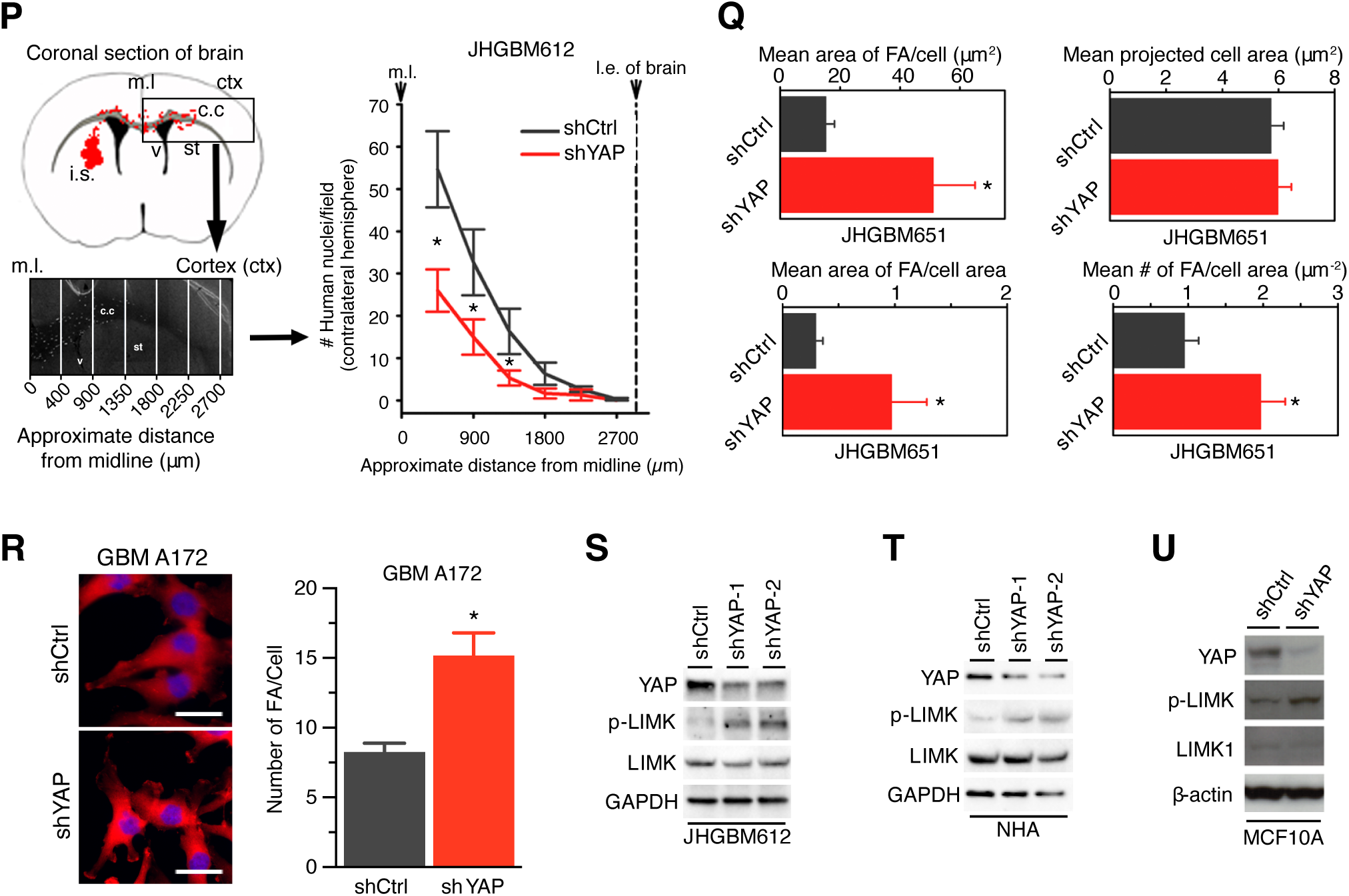
YAP regulates migration. **A,** Densitometric analysis of YAP and YAP/β-actin protein levels in Fig. 1A. Student’s t-test; * = P<0.05, between NCC (non-cancer cortex) and GBM. **B,** mRNA expression of YAP target genes, *Mcl-1* and *CTGF*, in glioblastoma tissues compared to the non-cancer cortex. Student’s t-test; * = P<0.05, comparing to NCC. Results are pooled from 3 independent experiments. **C,** Representative immunoblot of YAP and p-YAP protein levels in primary GBM cell cultures of 3 independent experiments. * = cells used in functional assays. Numbers correspond to Patient Sample ID listed in Suppl Table 1. **D,** Representative immunoblot of YAP protein level in wildtype (WT), shCtrl, or shYAP JHGBM612 cells with 5 independent shYAP constructs of 3 independent experiments. s.e. = short exposure, l.e. = long exposure. **E,** Representative immunoblot of YAP and TAZ protein levels in shCtrl or shYAP JHGBM651 cells with 2 independent shYAP constructs of 3 independent experiments. **F,** Representative immunoblot of YAP protein level in shCtrl or shYAP GBM A172 cells with 2 independent shYAP constructs of 3 independent experiments. **G-H,** Mean migration speed of shCtrl or shYAP MCF10A, NHA, JHGBM651/612/640/1A cells. Wilcoxon Rank-Sum test; * = P<0.05, comparing to shCtrl cells. **I, K-L,** Representative immunoblots of YAP protein level in the empty backbone vector transduced (CONT) or YAP OE NHA cells, JHGBM612, and JHGBM1A cells, respectively, of 3 independent experiments. **J, M,** Mean migration speed of the empty backbone vector transduced (CONT) or YAP OE NHA cells, and JHGBM1A cells, respectively. Wilcoxon Rank-Sum test; * = P<0.05, between CONT and YAP OE. **H, J, M,** Mean migration speed is calculated from 30 cells each group and are representative of 3 independent experiments. **N,** MRI images of Patient #30/JHGBM612. **O,** Representative H&E images showing the formation of solid tumors in the in vivo GBM xenograft model used in experiments presented in Fig.1E and Fig.5F after intracranial injection of JHGBM612 cells of 5 mice per group. Scale = 1mm. **P,** *Left:* Schematic of the murine intracranial xenograft model of GBM used to assess YAP’s migratory and invasive capacity in vivo*. Right:* Number of cells invading past the midline into the contralateral hemisphere. i.s. = injection site, c.c. = corpus callosum, st = striatum, ctx = cortex, v = ventricle, m.l. = midline, l.e. of brain = lateral edge of brain. Quantifications are per mouse and are pooled from 5 mice per group. Student’s t-test; * = P<0.05, between shYAP and shCtrl cells at a corresponding distance from midline. **Q,** Analysis of focal adhesion (FA) in shCtrl (N=25 cells) or shYAP (N=16 cells) JHGBM651 cells. Results are representative of 3 independent experiments. Student’s t-test; * = P<0.05, between shCtrl and shYAP. **R,** Analysis of focal adhesion (FA) in shCtrl (N=12 cells) or shYAP (N=12 cells) GBMA172 cells. Results are representative of 3 independent experiments. Scale = 5μm. Student’s t-test; * = P<0.05, between shCtrl and shYAP. **S-U,** Representative immunoblots of YAP, total and phospho-Thr^508^ LIMK protein levels in shCtrl, shYAP JHGBM612, NHA, and MCF10A cells of 3 independent experiments. 2 independent shYAP constructs were used in JHGBM612 and NHA cells. **A-B, G-H, J, M, Q-R,** Data shown is mean ± SEM.

**Figure S2.**
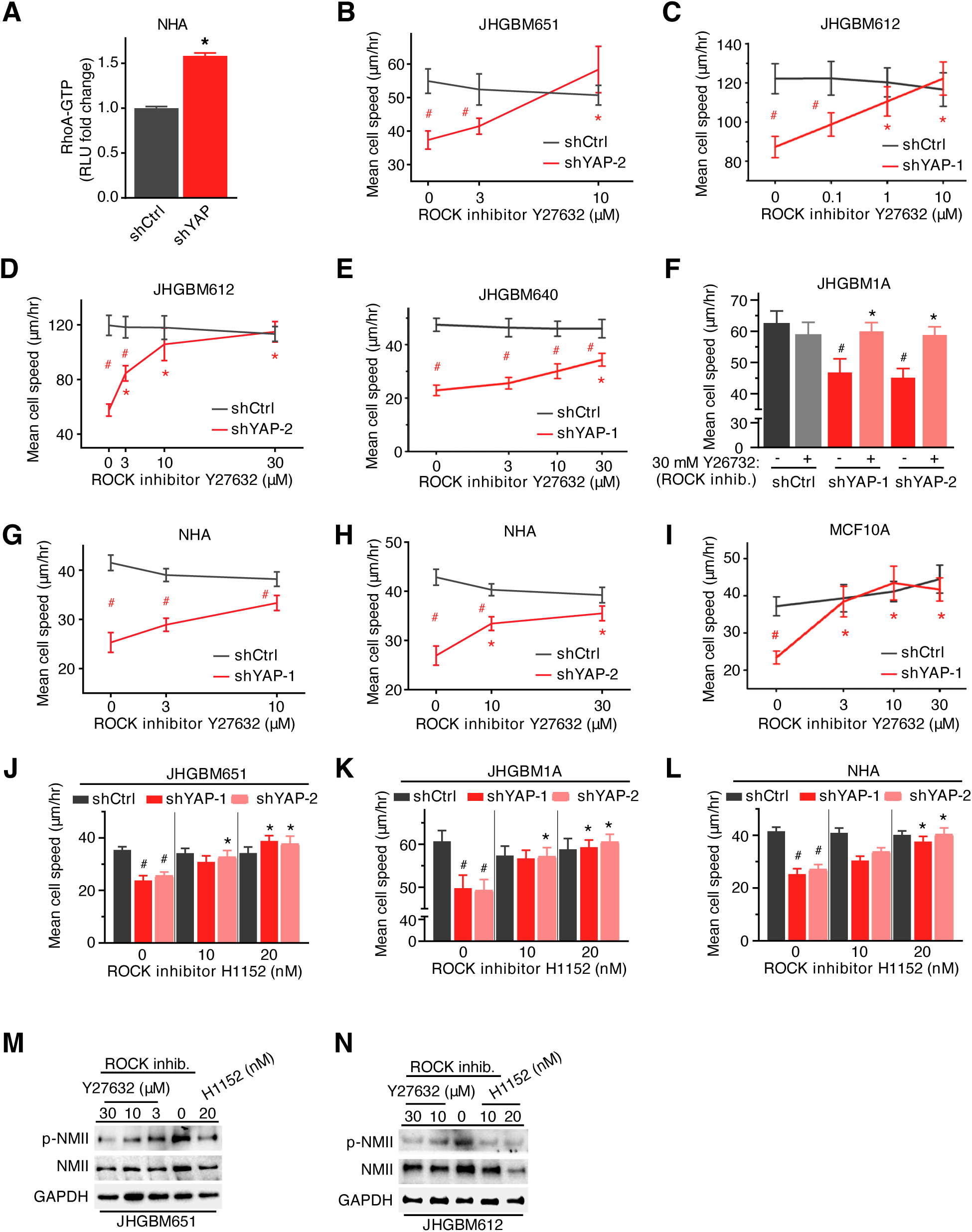
YAP regulates migration by inhibiting RhoA-ROCK signaling. **A,** G-LISA analysis of RhoA-GTP levels of shCtrl or shYAP NHA cells. Student’s t-test; * = P<0.05, between shCtrl and shYAP. Results are pooled from 3 independent experiments. **B-L,** Mean migration speed of shCtrl, shYAP-1, or -2 JHGBM651/612/640/1A, NHA, and MCF10A cells treated with increasing doses of ROCK inhibitors Y27632 or H1152. Mean migration speed is calculated from 30 cells. Results are representative of 3 independent experiments. Wilcoxon Rank-Sum test; Red and black * = P<0.05, comparing to no drug treatment in shYAP-1 or -2 cells; Red # = P<0.05, comparing to shCtrl cells at corresponding drug concentration; Black # = P<0.05, comparing to shCtrl cells with no drug treatment. **M-N,** Representative immunoblots of NMII and phospho-Ser^1943^ NMII expression in JHGBM651 or JHGBM612 cells treated with vehicle or ROCK inhibitors Y27632 or H1152 of 3 independent experiments. **A-L,** Data shown is mean ± SEM.

**Figure S3.**
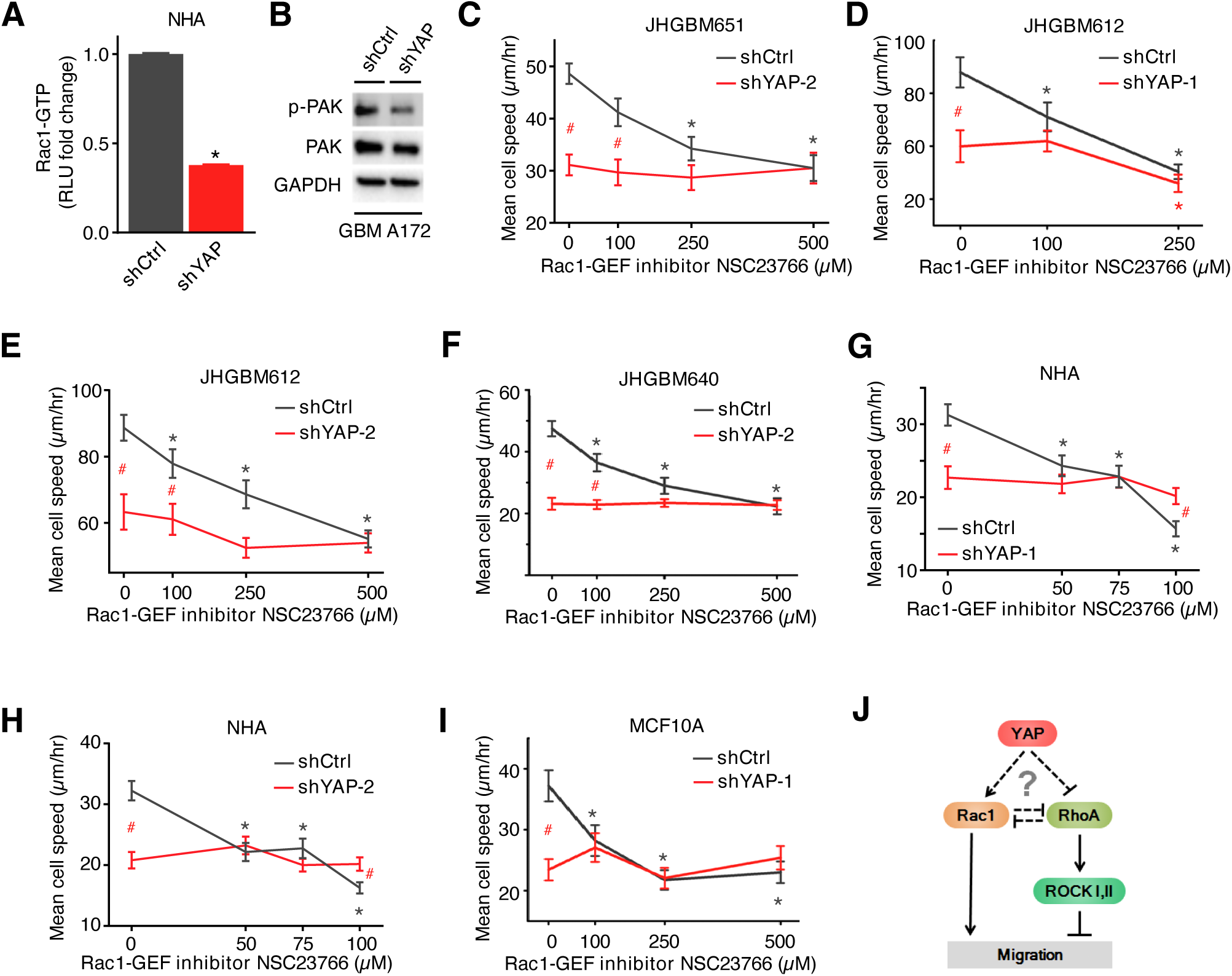
YAP regulates migration by activating Rac1 signaling. **A,** G-LISA analysis of Rac1-GTP levels of shCtrl or shYAP NHA cells. Results are pooled from 3 independent experiments. Student’s t-test; * = P<0.05, between shCtrl and shYAP. **B,** Representative immunoblot of YAP, total and phospho-Thr^423/402^ PAK in shCtrl and shYAP GBMA172 cells of 3 independent experiments. **C-I,** Mean migration speed of shCtrl, shYAP-1, or -2 JHGBM651/612/640, NHA, and MCF10A cells treated with the Rac1 inhibitor NSC23766. Mean migration speed is calculated from 30 cells. Results are representative of 3 independent experiments. Wilcoxon Rank-Sum test; Black * = P<0.05, comparing to no drug treatment in shCtrl cells; Red # = P<0.05, between shYAP-1 or -2 and shCtrl cells at corresponding drug concentrations. **J,** Schematic representation of YAP modulating Rho-GTPases. **A, C-I,** Data shown is mean ± SEM.

**Figure S4.**
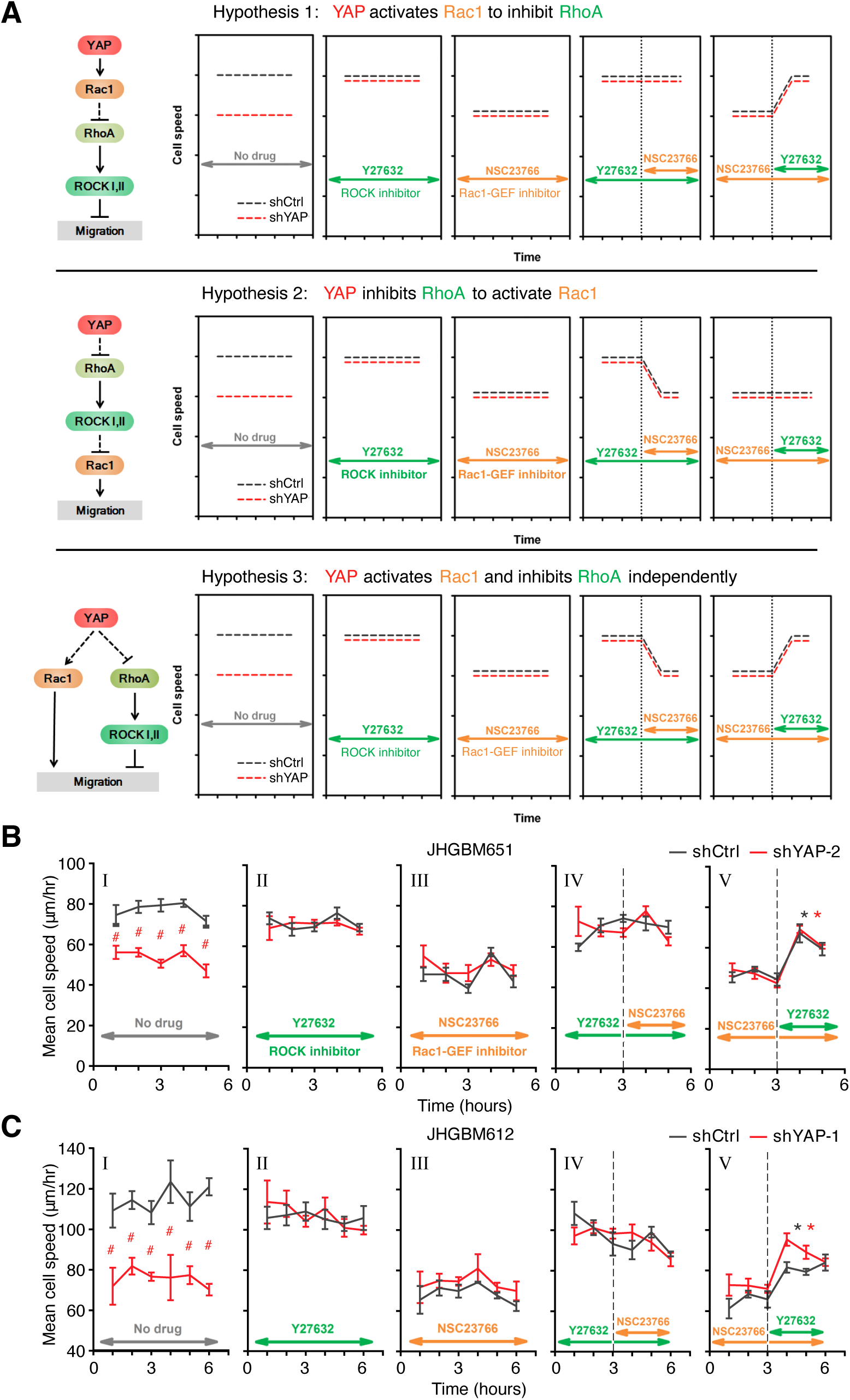
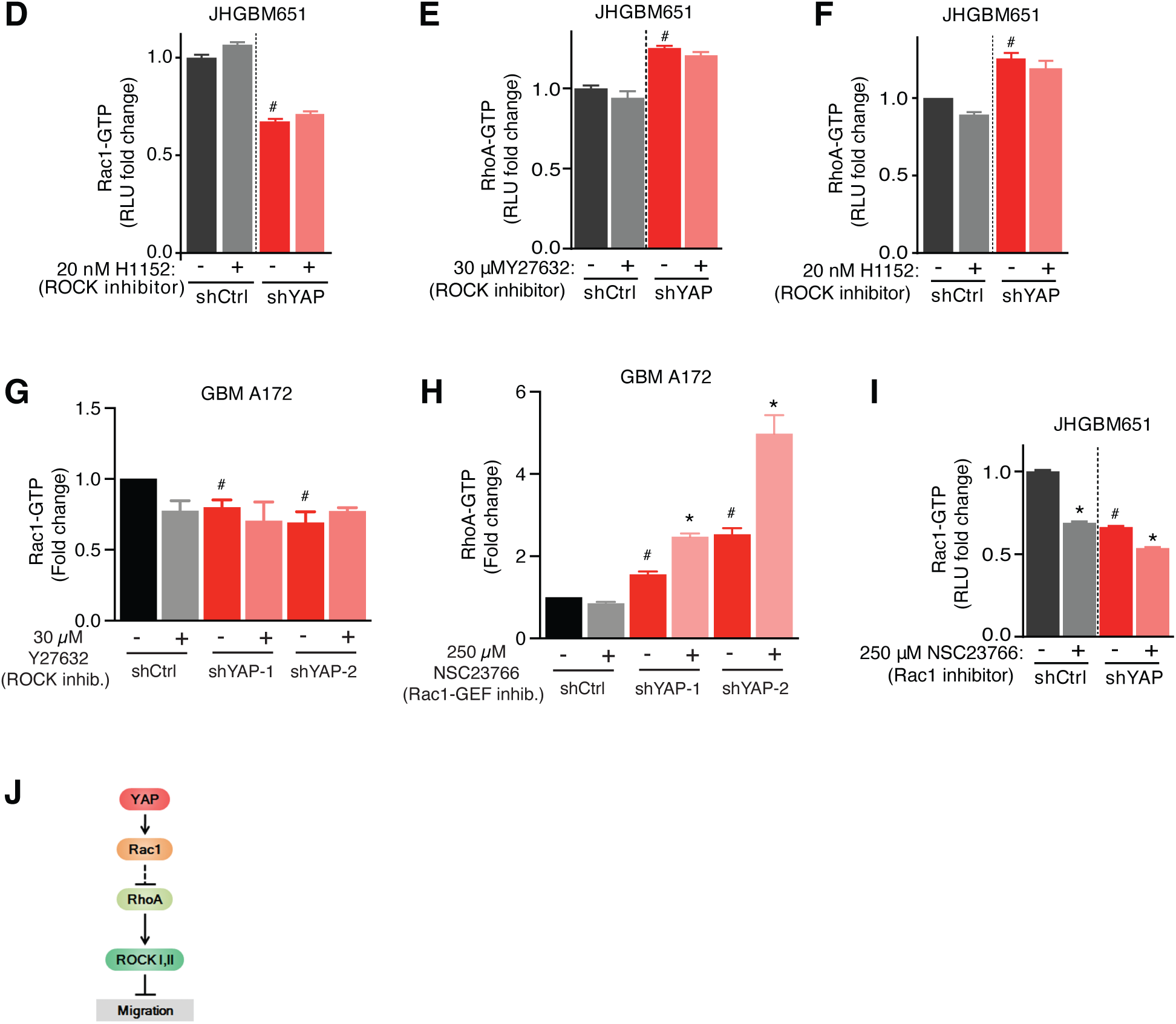
YAP regulates migration by modulating a Rho-GTPase switch. **A,** Experimental paradigm to delineate the hierarchy of molecular events of YAP’s modulation of a Rho-GTPase switch. **B-C,** Mean migration speed of shCtrl, shYAP-1, or -2 JHGBM651 and JHGBM612 cells treated with no drug/vehicle, 10μM Y27632, or 250μM NSC23766 for 3 or 6 hours as indicated. Mean migration speed is calculated from 30 cells. Results are representative of 3 independent experiments. Wilcoxon Rank-Sum test; Black * = P<0.05, between mean speed from hour1 to hour2 and that from hour4 to hour5 in shCtrl cells; Red *= P<0.05, between mean speed from hour1 to hour2 and that from hour4 to hour5 in shYAP-1 or -2 cells; Red # = P<0.05, between mean speed in shYAP-1 or -2 cells and that in shCtrl cells at corresponding time points. **D-F, I,** G-LISA analysis of Rac1-GTP or RhoA-GTP levels of shCtrl or shYAP cells treated with vehicle, ROCK inhibitors Y27632 (30μM) or H1152 (20nM), or Rac1-GEF inhibitor NSC23766 (250μM). Results are pooled from 3 independent experiments. Student’s t-test; * = P<0.05, comparing to no drug treatment; # = P<0.05, comparing to shCtrl cells with no drug treatment. **G-H,** Rac1-GTP and RhoA-GTP pull-down assay of shCtrl or shYAP cells treated with vehicle, ROCK inhibitor Y27632 (30μM), or Rac1-GEF inhibitor NSC23766 (250uM). Results are pooled from 3 independent experiments. Student’s t-test; * = P<0.05, comparing to no drug treatment; # = P<0.05, ## = P<0.01, comparing to shCtrl cells with no drug treatment. **J,** Schematic representation of the hierarchy of YAP’s modulation of a Rho-GTPase switch. **B-I,** Data shown is mean ± SEM.

**Figure S5.**
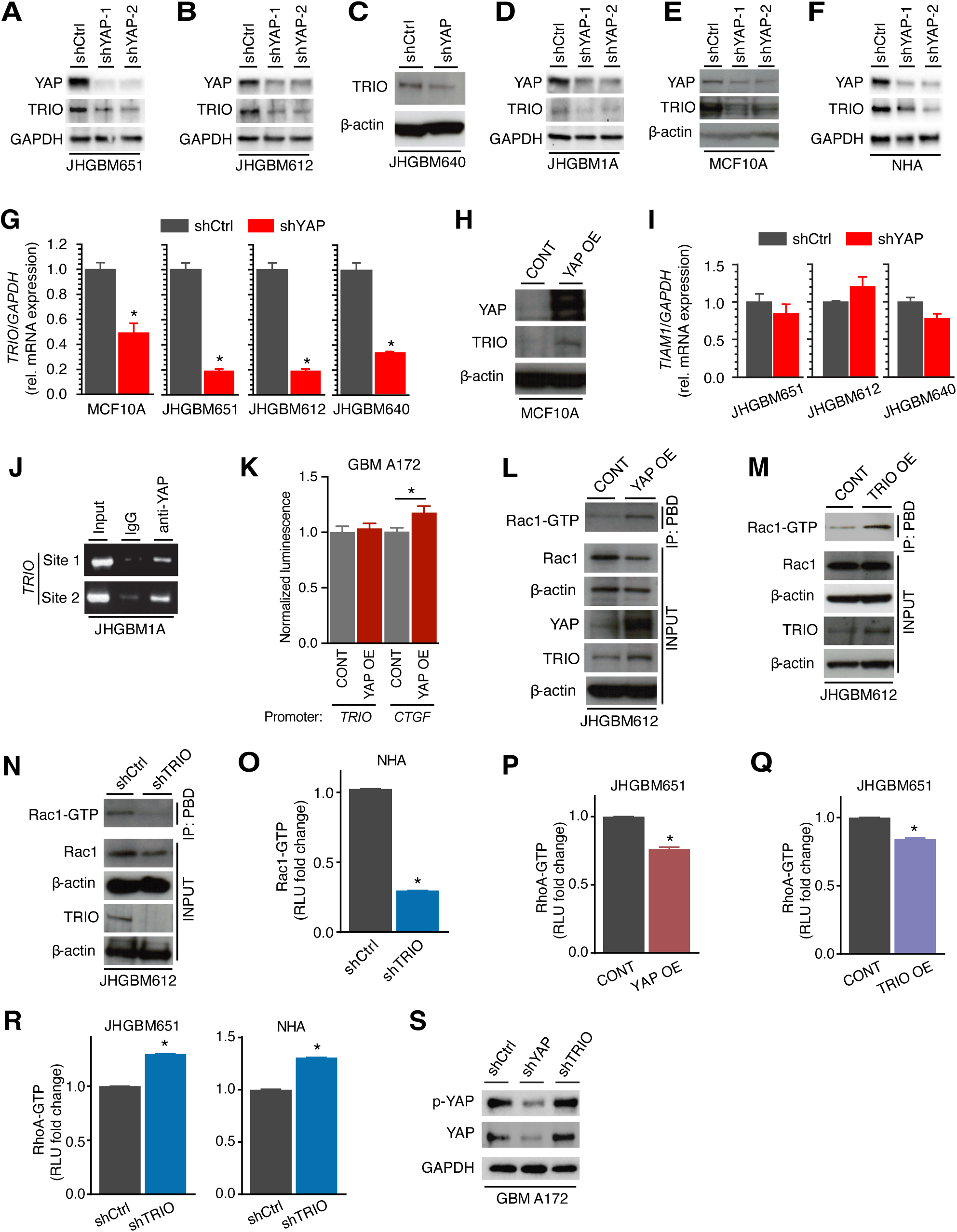
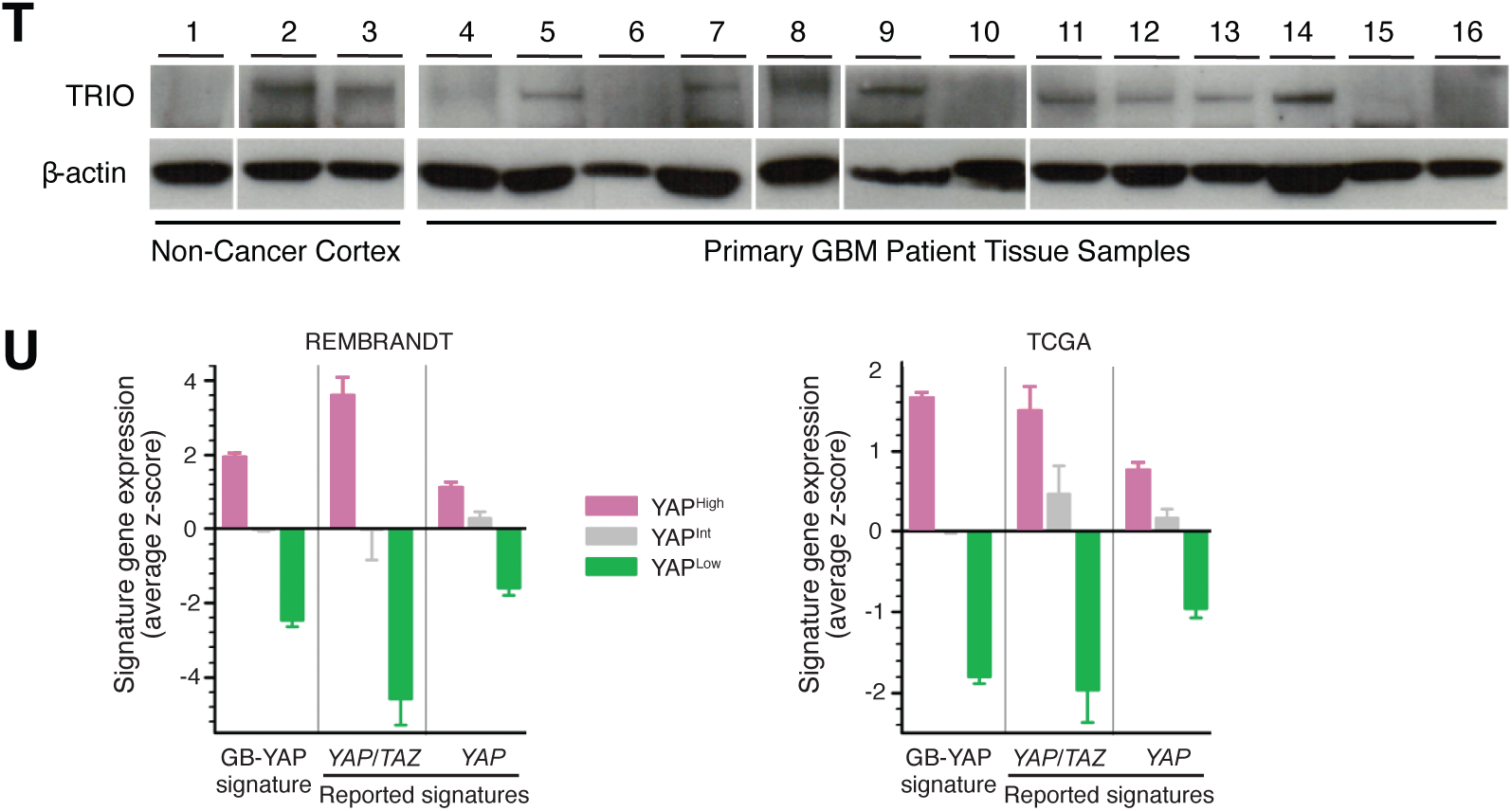
YAP transactivates TRIO signaling. **A-F,** Representative immunoblots of YAP and TRIO protein levels in shCtrl, shYAP JHGBM651/612/640/1A, MCF10A, and NHA cells of 3 independent experiments. 2 independent shYAP constructs are used in JHGBM651/612/1A, MCF10A and NHA cells. **G,** Relative *TRIO* mRNA expression in shCtrl or shYAP JHGBM651/612/640 and MCF10A cells. Results are pooled from 3 independent experiments. Student’s t-test; * = P<0.05, between shCtrl and shYAP. **H,** Representative immunoblot of YAP and TRIO protein levels in the empty backbone vector transduced (CONT) or YAP OE MCF10A cells of 3 independent experiments. **I,** Relative *TIAM1* mRNA expression in shCtrl or shYAP JHGBM651/612/640 cells. Results are pooled from 3 independent experiments. **J,** ChIP-PCR querying across two enhancer elements of *TRIO* in JHGBM1A cells. Results are representative of 3 independent experiments. **K,** GFP-luciferase reporter assay with the promoter sequence of *TRIO* and *CTGF* (positive control) in the empty backbone vector transduced (CONT) and YAP OE GBMA172 cells. Results are pooled from 3 independent experiments. Student’s t-test; * = P<0.05, between CONT and YAP OE. **L-N,** Representative immunoblots of pull-down analysis of Rac1-GTP levels in the empty backbone vector transduced (CONT) and YAP OE JHGBM612 cells, and TRIO OE JHGBM612 cells or shCtrl and shTRIO JHGBM612 cells, respectively, of 3 independent experiments. **O,** G-LISA analysis of Rac1-GTP levels in shCtrl and shTRIO NHA cells. Results are pooled from 3 independent experiments. Student’s t-test; * = P<0.05, between shCtrl and shTRIO. **P-Q,** G-LISA analysis of RhoA-GTP levels in the empty backbone vector transduced (CONT), YAP OE, and TRIO OE JHGBM651 cells. Results are pooled from 3 independent experiments. Student’s t-test; * = P<0.05, between CONT and YAP or TRIO OE. **R,** G-LISA analysis of RhoA-GTP levels in shCtrl and shTRIO JHGBM651 and NHA cells. Results are pooled from 3 independent experiments. Student’s t-test; * = P<0.05, between shCtrl and shTRIO. **S,** Representative immunoblots of total and phospho-Ser^127^ YAP protein levels in shCtrl, shYAP, and shTRIO GBMA172 cells of 3 independent experiments. **T,** Representative immunoblots of TRIO protein level in non-cancer cortex and patient-derived primary glioblastoma tissues. Results are representative of 3 independent experiments using the same set of non-cancer cortex and GBM samples. **U,** Expression of YAP/TAZ and YAP signatures, derived from a variety of other tissues, in YAP high, intermediate, or low patient groups in the REMBRANDT and TCGA GBM datasets. **G, I, K, O-R, U,** Data shown is mean ± SEM.

**Figure S6.**
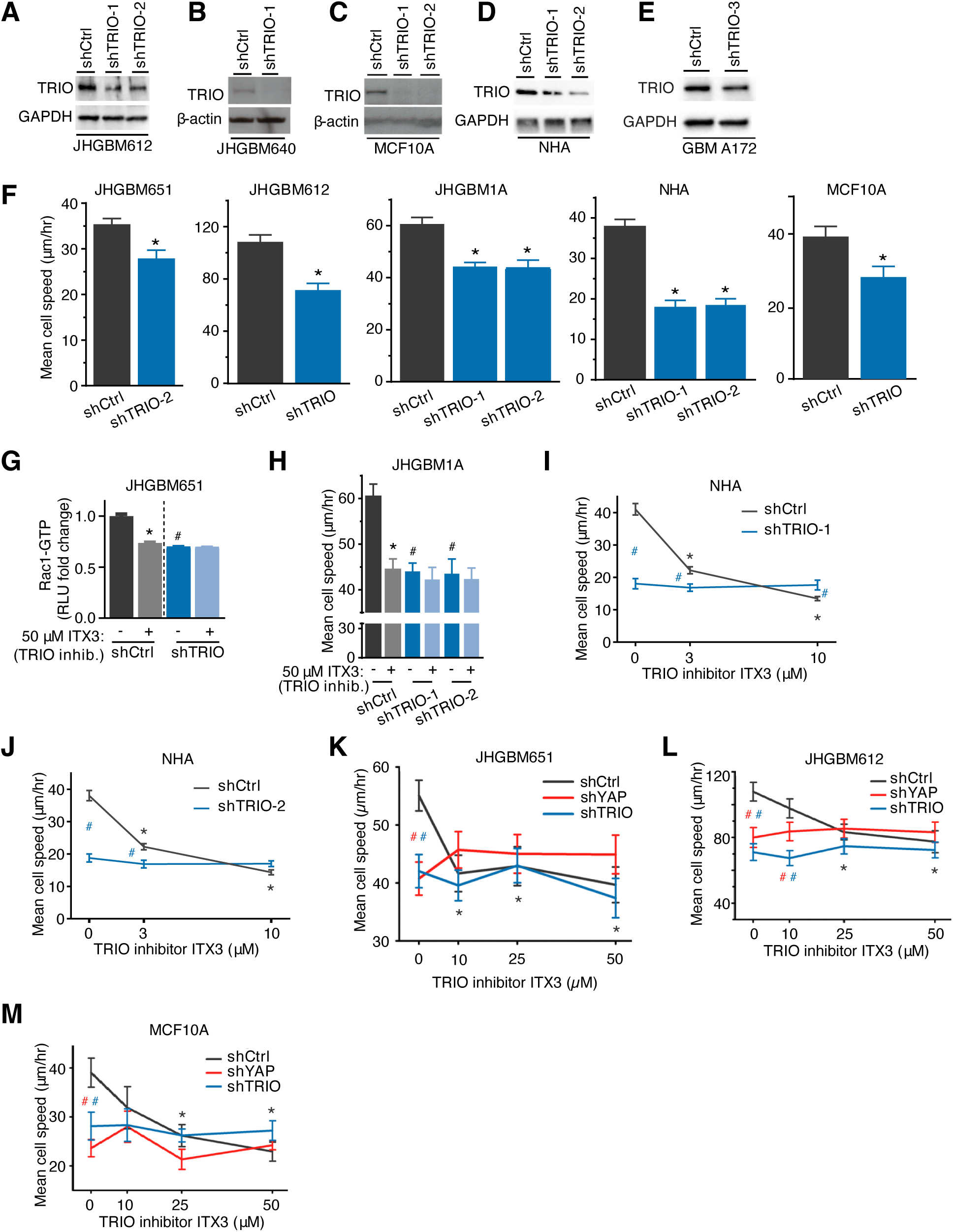
YAP regulates migration through TRIO signaling. **A-E,** Representative immunoblots of TRIO protein level in shCtrl, shTRIO JHGBM612/640, MCF10A, NHA and GBMA172 cells of 3 independent experiments. 2 independent shTRIO constructs are used in JHGBM612, MCF10A, and NHA cells. **F,** Mean migration speed of shCtrl, shTRIO-1, or -2 JHGBM651/612/1A, NHA, and MCF10A cells. Wilcoxon Rank-Sum test; * = P<0.05, comparing to shCtrl cells. **G,** G-LISA analysis of Rac1-GTP levels in shCtrl or shTRIO JHGBM651 cells treated with vehicle or the TRIO inhibitor ITX3 (50μM). Results are pooled from 3 independent experiments. Student’s t-test; * = P<0.05, comparing to no drug treatment; # = P<0.05, comparing to shCtrl with no drug treatment. **H,** Mean migration speed of shCtrl, shTRIO-1, or -2 GBM1A cells treated with vehicle or TRIO inhibitor ITX3 (50μM). Wilcoxon Rank-Sum test; * = P<0.05, comparing to no drug treatment; # = P<0.05, comparing to shCtrl with no drug treatment. **I-M,** Mean migration speed of shCtrl, shTRIO-1, or -2, or shYAP-1, or -2 JHGBM651/612, NHA, and MCF10A cells treated with increasing doses of the TRIO inhibitor ITX3. Wilcoxon Rank-Sum test; Black * = P<0.05, comparing to no drug treatment in shCtrl cells; Red # = P<0.05, between shYAP and shCtrl at corresponding drug concentration; Blue # = P<0.05, between shTRIO-1 or -2 and shCtrl at corresponding drug concentration. **F, H-M,** Mean migration speed is calculated from 30 cells. Results are representative of 3 independent experiments. **F-M,** Data shown is mean ± SEM.

**Figure S7.**
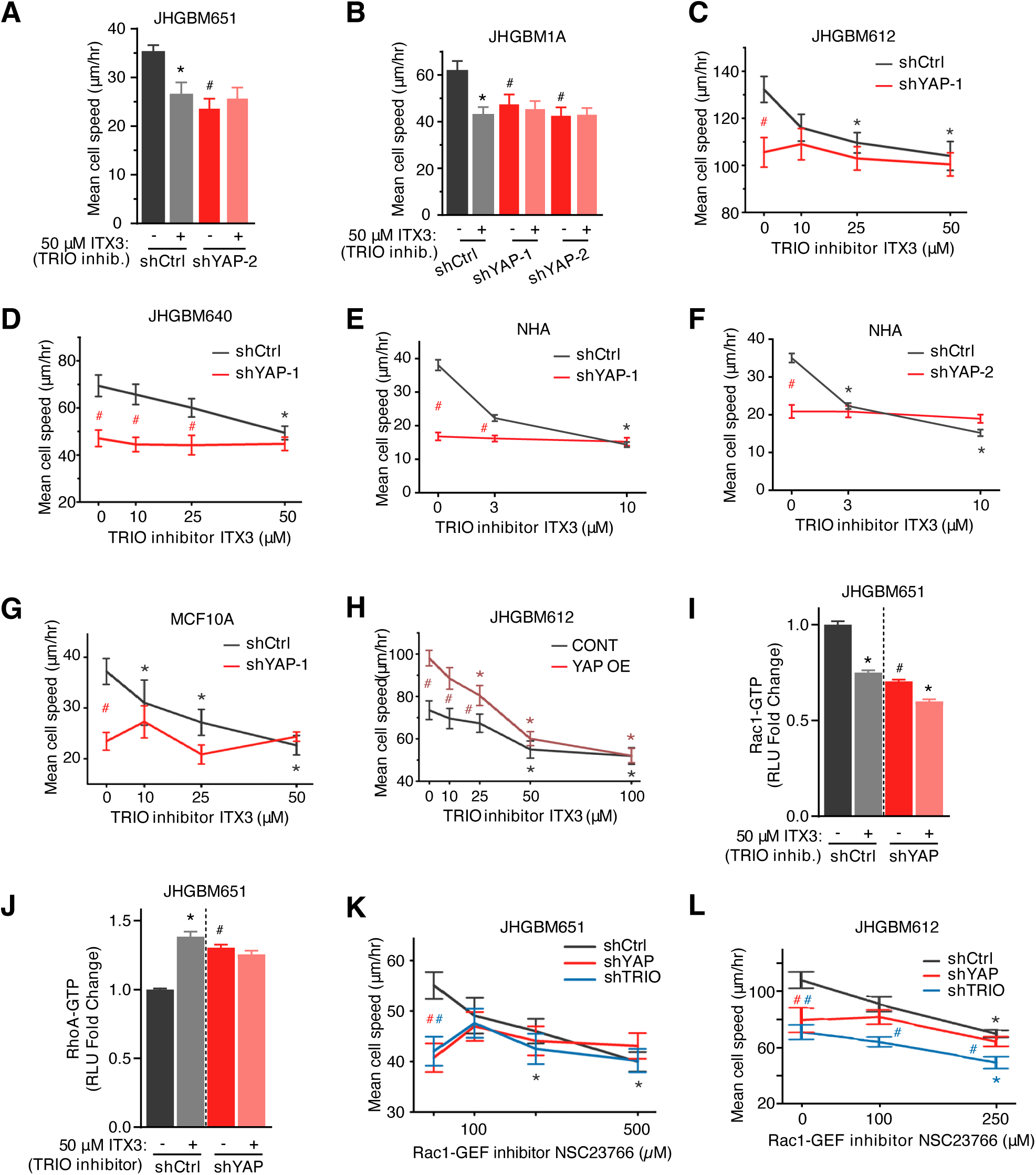
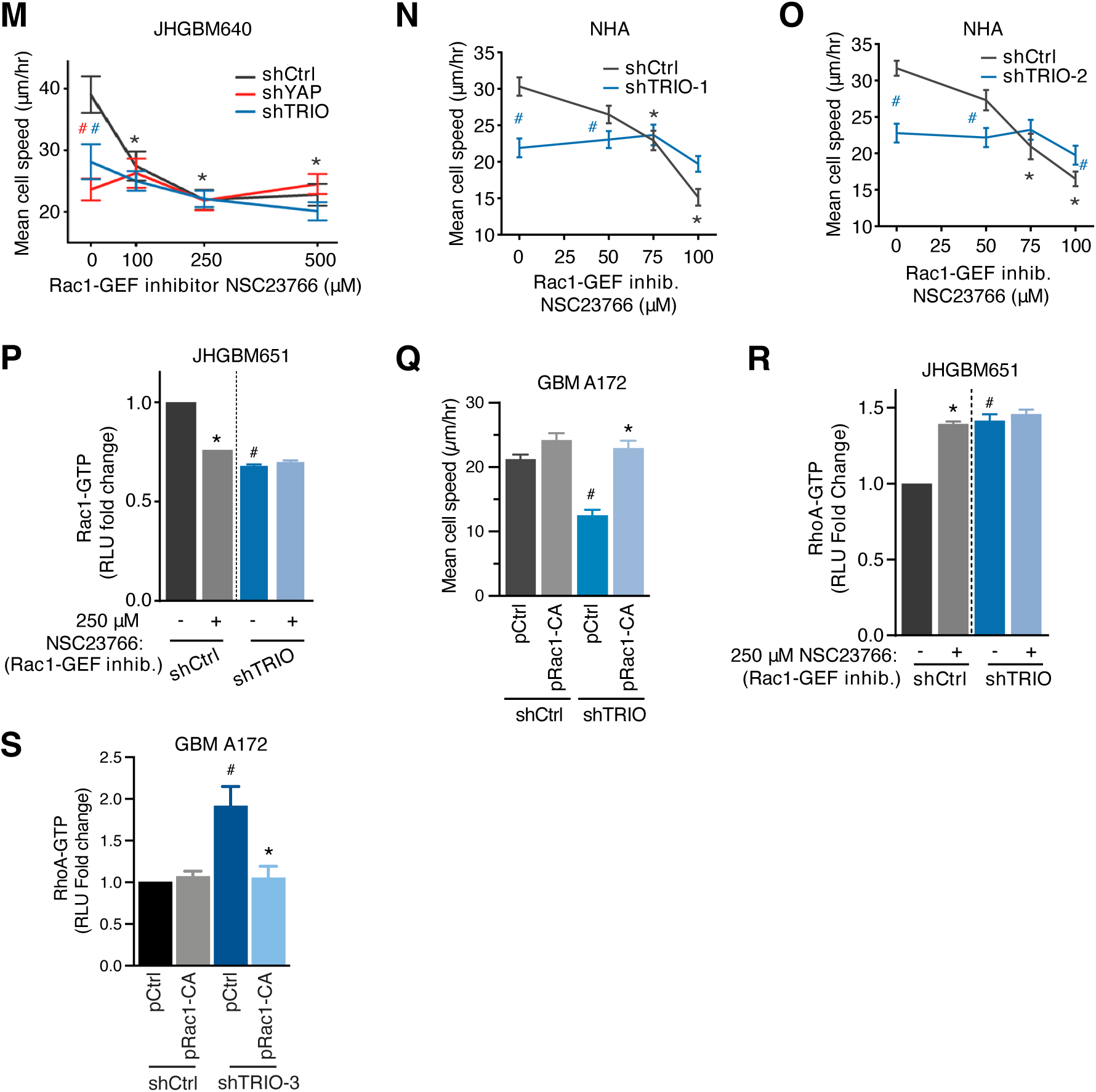
YAP-TRIO signaling modulates Rac1-RhoA to drive migration. **A-G,** Mean migration speed of shCtrl, shYAP-1, or -2 JHGBM651/612/640/1A, NHA, and MCF10A cells treated with increasing doses of the TRIO inhibitor ITX3. **H,** Mean migration speed of the empty backbone vector transduced (CONT) or YAP OE JHGBM612 cells treated with vehicle or increasing doses of the TRIO inhibitor ITX3. **A-H,** Wilcoxon Rank-Sum test; Black * = P<0.05, comparing to no drug treatment in shCtrl or CONT cells; Red * = P<0.05, comparing to no drug treatment in shYAP or YAP OE cells; Red # = P<0.05, between YAP OE and CONT at corresponding drug concentration. **I-J,** G-LISA analysis of Rac1-GTP or RhoA-GTP levels in shCtrl or shYAP JHGBM651 cells treated with vehicle or the TRIO inhibitor ITX3 (50μM). Results are pooled from 3 independent experiments. Student’s t-test; * = P<0.05, comparing to no drug treatment in shCtrl or shYAP cells; # = P<0.05, comparing to shCtrl cells with no drug treatment. **K-O,** Mean migration speed of shCtrl, shYAP, or shTRIO JHGBM651/612/640 and NHA cells treated with increasing doses of the Rac1-GEF inhibitor NSC23766. Wilcoxon Rank-Sum test; Black * = P<0.05, comparing to no drug treatment in shCtrl cells; Red # = P<0.05, between shYAP-1 or -2 and shCtrl cells at corresponding drug concentration; Blue # = P<0.05, between shTRIO-1 or -2 and shCtrl cells at corresponding drug concentration. **P, R,** G-LISA analysis of Rac1-GTP or RhoA-GTP levels in shCtrl or shTRIO JHGBM651 cells treated with vehicle or the Rac1-GEF inhibitor NSC23766 (250μM). Results are pooled from 3 independent experiments. Student’s t-test; * = P<0.05, comparing to no drug treatment in shCtrl cells; # = P<0.05, comparing to shCtrl cells with no drug treatment. **Q,** Mean migration speed of shCtrl and shTRIO GBMA172 cells transfected with the empty backbone vector (pCtrl) or a constitutively active Rac1 vector (pRac1-CA). Wilcoxon Rank-Sum test; * = P<0.05, comparing to pCtrl in shTRIO cells; # = P<0.05, comparing to pCtrl shCtrl cells. **S,** RhoA-GTP pull-down assay of shCtrl or shTRIO GBMA172 cells transfected with the empty backbone vector (pCtrl) or a constitutively active Rac1 vector (pRac1-CA). Student’s t-test; * = P<0.05, comparing to shTRIO cells transfected with pCtrl; # = P<0.05, comparing to shCtrl cells transfected with pCtrl. **A-H, K-O, Q,** Mean migration speed is calculated from 30 cells. Results are representative of 3 independent experiments. All data shown is mean ± SEM.

**Figure S8.**
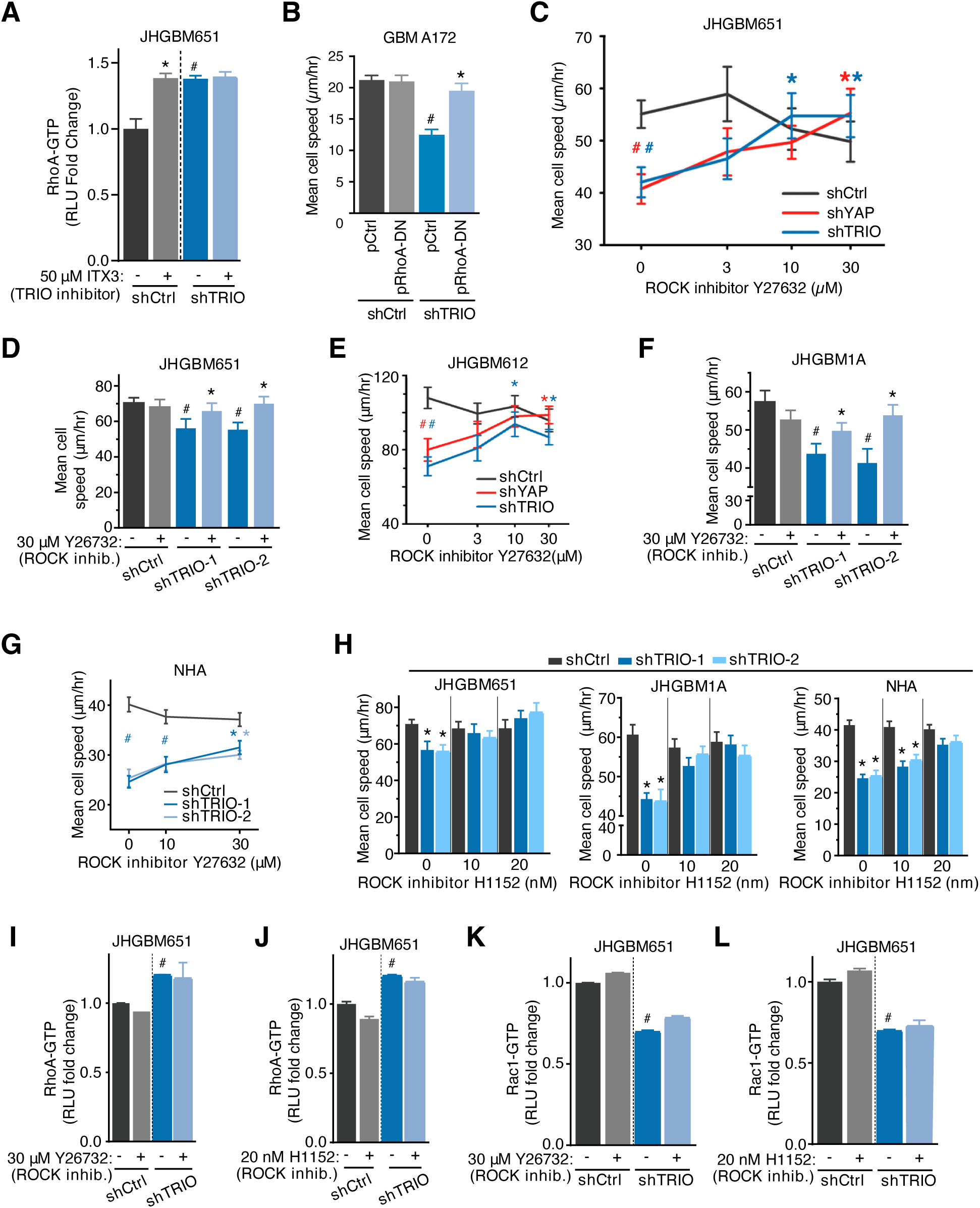
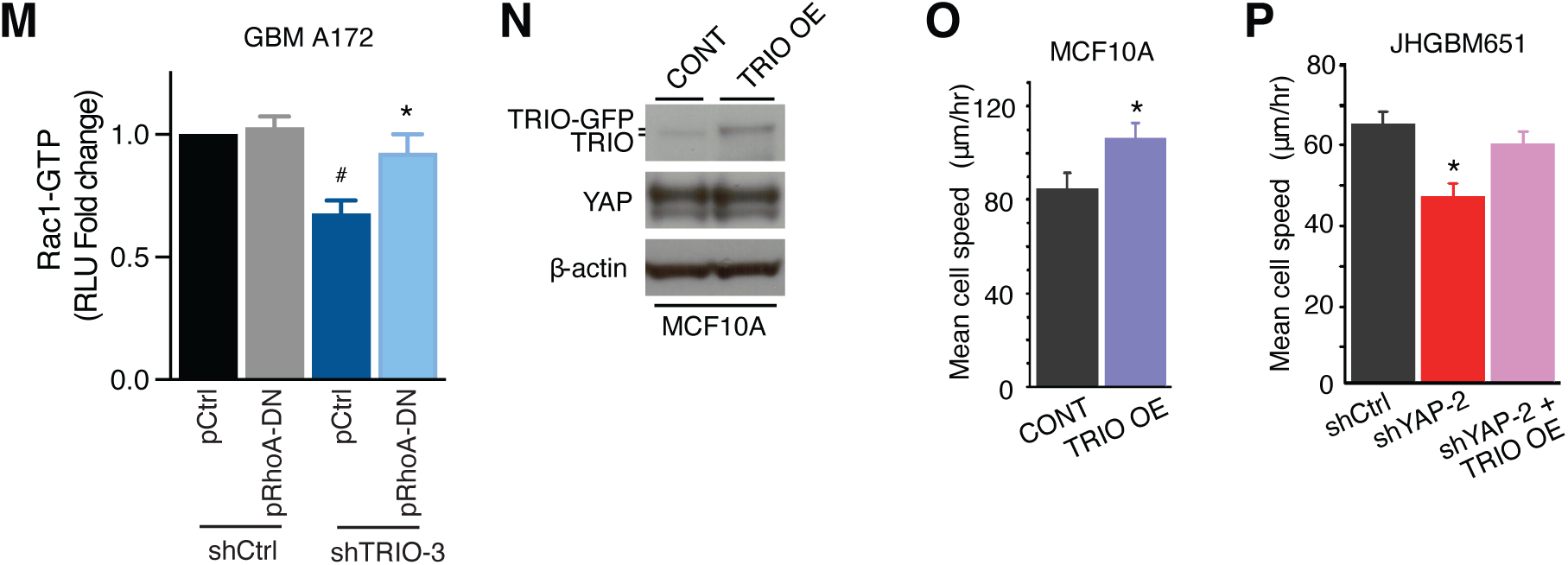
YAP-TRIO signaling regulates migration by modulating Rac1-RhoA crosstalk. **A,** G-LISA analysis of RhoA-GTP levels in shCtrl or shTRIO JHGBM651 cells treated with vehicle or the TRIO inhibitor ITX3 (50μM). Results are pooled from 3 independent experiments. Student’s t-test; * = P<0.05, comparing to no drug treatment in shCtrl cells; # = P<0.05, comparing to shCtrl cells with no drug treatment. **B,** Mean migration speed of shCtrl and shTRIO GBMA172 cells transfected with the empty backbone vector (pCtrl) or a dominant-negative RhoA vector (pRhoA-DN). Wilcoxon Rank-Sum test; * = P<0.05, comparing to pCtrl in shTRIO cells; # = P<0.05, comparing to pCtrl shCtrl cells. **C, E, G-H,** Mean migration speed of shCtrl, shYAP, or shTRIO JHGBM651/612/1A, and NHA cells treated with increasing doses of the ROCK inhibitor Y27632 or H1152. Wilcoxon Rank-Sum test; Black * = P<0.05, comparing to no drug treatment in shCtrl or shTRIO cells; Red * = P<0.05, comparing to no drug treatment in shYAP cells; Blue * = P<0.05, comparing to no drug treatment in shTRIO cells; Red # = P<0.05, between shYAP and shCtrl cells at corresponding drug concentration; Blue # = P<0.05, between shTRIO and shCtrl cells at corresponding drug concentration. **D, F,** Mean migration speed of shCtrl, shTRIO-1 or -2 JHGBM651/1A and NHA cells treated with vehicle or the ROCK inhibitor Y26732 (30μM). Wilcoxon Rank-Sum test; * = P<0.05, comparing to no drug treatment in shTRIO cells; # = P<0.05, comparing to no drug treatment in shCtrl cells. **I-L,** G-LISA analysis of Rac1-GTP or RhoA-GTP levels in shCtrl or shTRIO JHGBM651 cells treated with vehicle or the ROCK inhibitor Y27632 (30μM) or H1152 (20nM). Results are pooled from 3 independent experiments. Student’s t-test; # = P<0.05, comparing to no drug treatment in shCtrl cells. **M,** Rac1-GTP pull-down assay of shCtrl or shTRIO GBMA172 cells transfected with the empty backbone vector (pCtrl) or a dominant-negative vector (pRhoA-DN). Student’s t-test; * = P<0.05, comparing to shTRIO cells transfected with pCtrl; # = P<0.05, comparing to shCtrl cells transfected with pCtrl. **N**, Representative immunoblots of TRIO and YAP protein levels in the empty backbone vector transduced (CONT) of TRIO OE MCF10A cells of 3 independent experiments. **O,** Mean migration speed of the empty backbone vector transduced (CONT) or TRIO OE MCF10A cells. Wilcoxon Rank-Sum test, * = P<0.05, comparing to CONT cells. **P,** Mean migration speed of shCtrl, shYAP-2, and shYAP-2+TRIO OE JHGBM651 cells. Wilcoxon Rank-Sum test, * = P<0.05, comparing to shCtrl cells. **B-H, O-P,** Mean migration speed is calculated from 30 cells. Results are representative of 3 independent experiments. **A-M, O-P,** Data shown is mean ± SEM.

**Figure S9.**
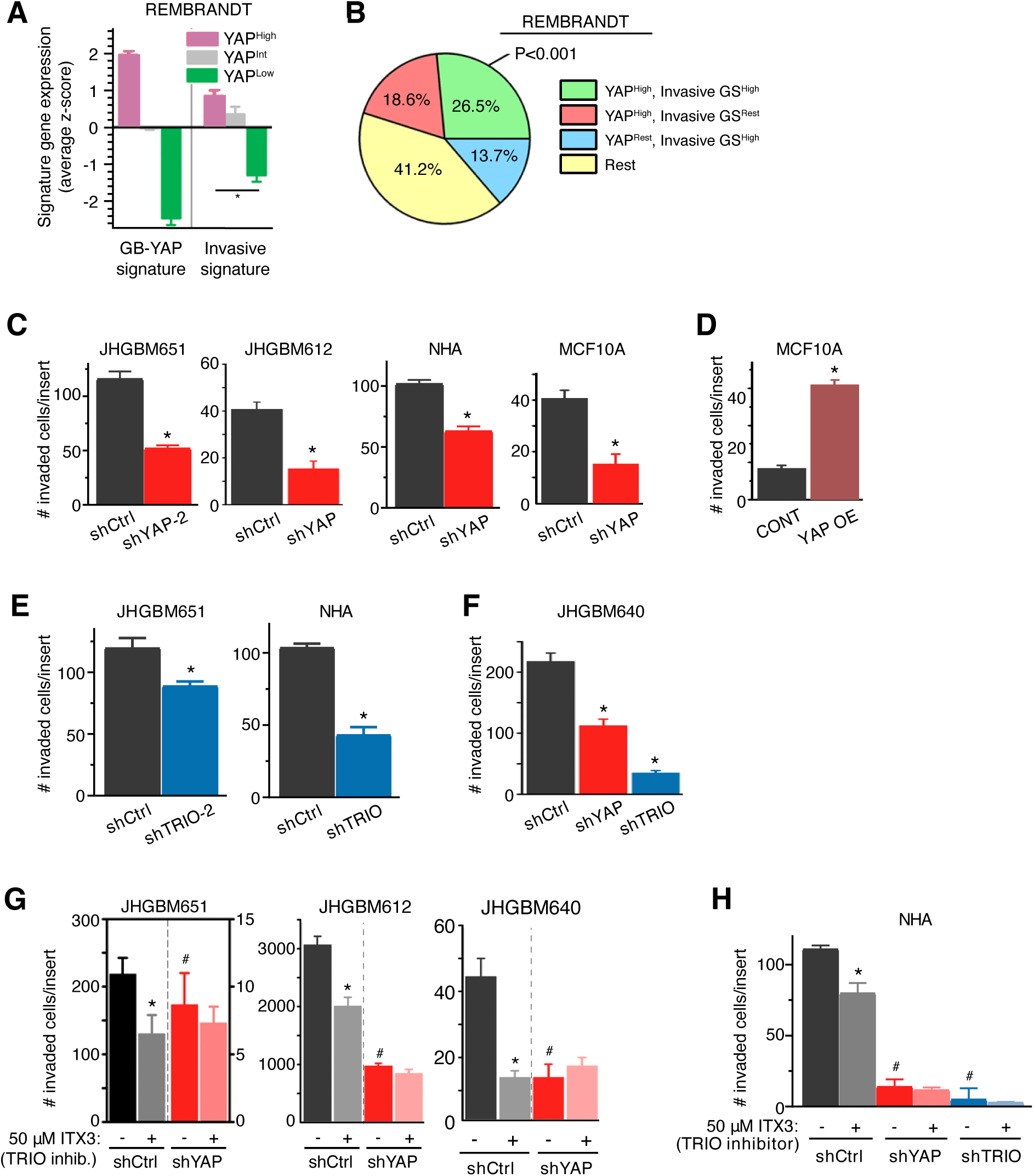
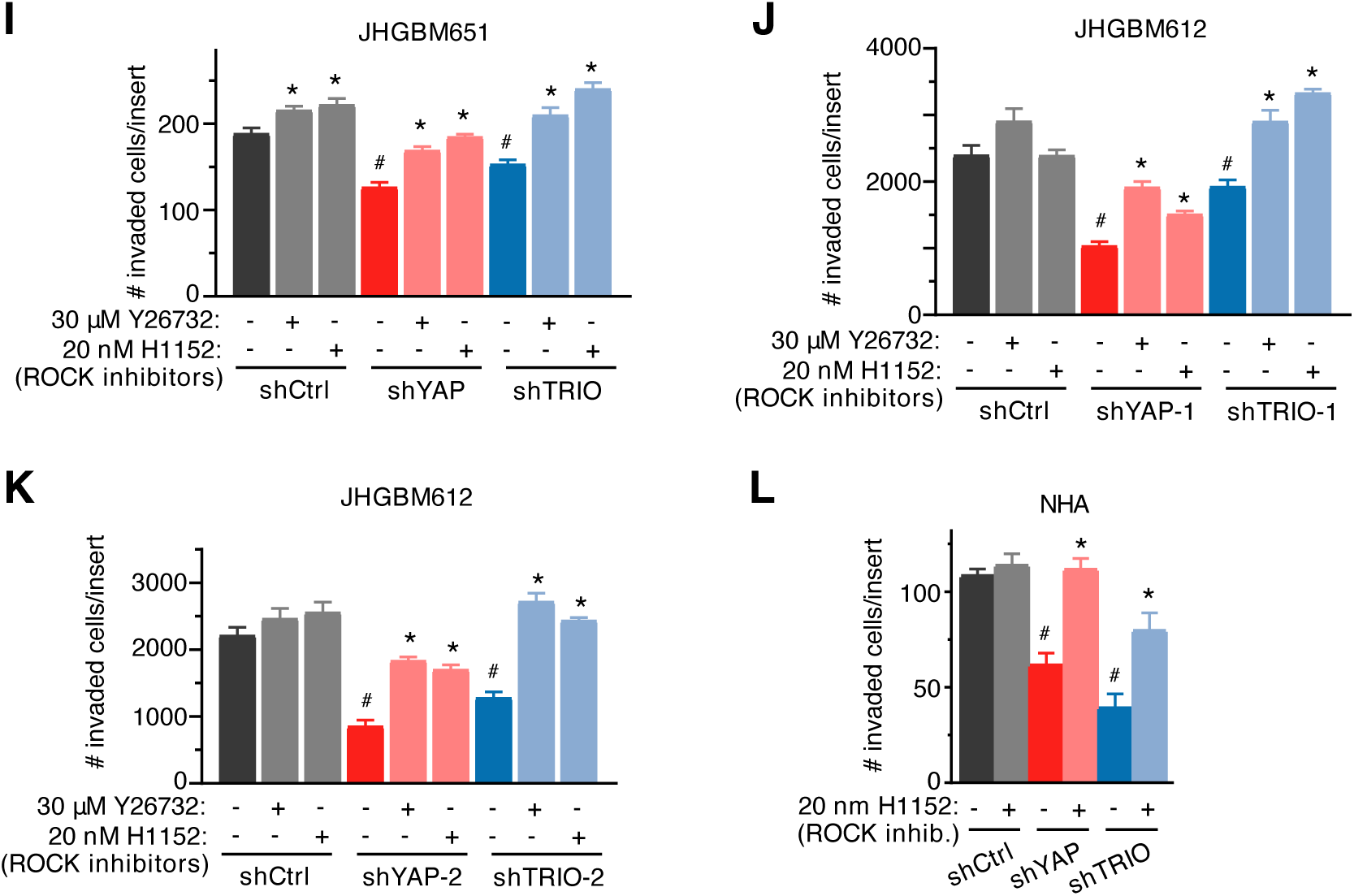
YAP-TRIO signaling promotes cell invasion. **A,** Expression of the Invasive signatures in YAP high, intermediate, or low patient groups in the REMBRANDT GBM dataset. Student’s t-test; * = P<0.05, between YAP^High^ and YAP^Low^. **B,** Percentage of patients with elevated GB-YAP signature, Invasive signature, or both in the REMBRANDT GBM dataset. P = Fisher’s exact test using the sizes of the shown patient groups. **C-D,** Matrigel Boyden invasion assay of shCtrl, shYAP-1, or -2 JHGBM651/612, NHA, MCF10A cells and the empty backbone vector transduced (CONT) or YAP OE MCF10A cells, respectively. Results are pooled from 3 independent experiments. Student’s t-test, * = P<0.05, comparing to shCtrl or CONT cells. **E,** Matrigel Boyden invasion assay of shCtrl, shTRIO, or -2 JHGBM651 and NHA cells. Results are pooled from 3 independent experiments. Student’s t-test; * = P<0.05, comparing to shCtrl cells. **F,** Matrigel Boyden invasion assay of shCtrl, shYAP and shTRIO JHGBM640 cells. Results are pooled from 3 independent experiments. Student’s t-test; * = P<0.05, comparing to shCtrl cells. **G-H,** Matrigel Boyden invasion assay of shCtrl, shYAP, or shTRIO JHGBM651/612/640 and NHA cells treated with the TRIO inhibitor ITX3 (50μM). Results are pooled from 3 independent experiments. **I-L,** Matrigel Boyden invasion assay of shCtrl, shYAP-1/2, or shTRIO1/2 JHGBM651/612 and NHA cells treated with the ROCK inhibitors Y26732 (30μM) or H1152 (20nM). Results are pooled from 3 independent experiments. **G-L,** Student’s t-test; * = P<0.05, comparing to no drug treatment in shCtrl cells; # = P<0.05, comparing to shCtrl cells with no drug treatment. **A, C-L,** Data shown is mean ± SEM.

**Figure S10.**
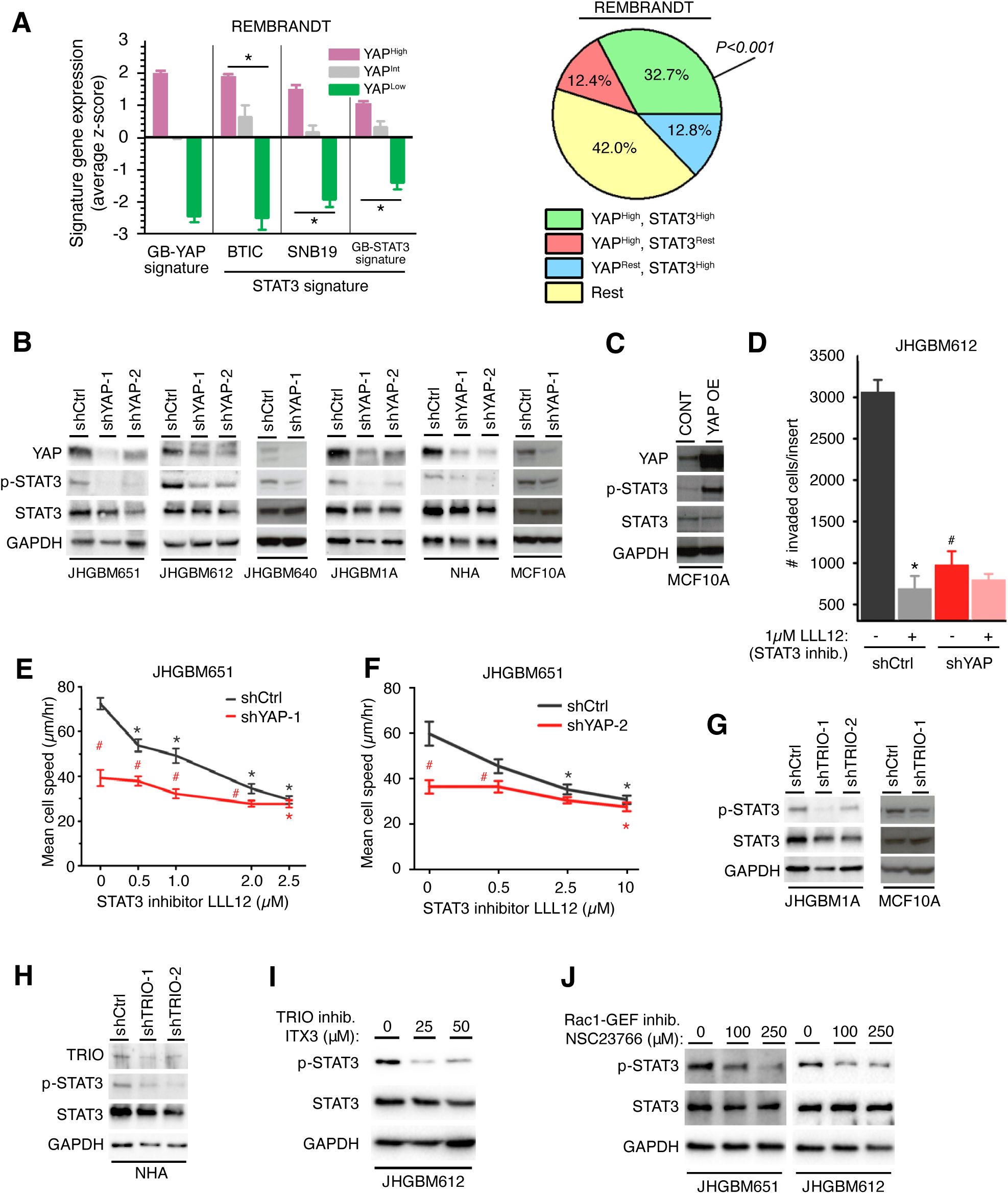
STAT3 is potentially involved in YAP-mediated cell invasion. **A,** Left: Expression of different GB-YAP and STAT3 signatures in YAP high, intermediate, or low patient groups in the REMBRANDT GBM dataset. Student’s t-test; * = P<0.05, between YAP^High^ and YAP^Low^. Right: Percentage of patients with elevated GB-YAP or GB-STAT3 signature in the REMBRANT GBM dataset. P = Fisher’s exact test. **B-C,** Representative immunoblots of YAP, total and phospho-Tyr^705^ STAT3 protein levels in shCtrl, shYAP-1, or -2 JHGBM651/612/640/1A, NHA, and MCF10A cells and the empty backbone vector transduced (CONT) or YAP OE MCF10A cells of 3 independent experiments. **D,** Matrigel Boyden invasion assay of shCtrl and shYAP JHGBM612 cells treated with vehicle or the STAT3 inhibitor LLL12 (1μM). Results are pooled from 3 independent experiments. Student’s t-test; * = P<0.05, comparing to no drug treatment in shCtrl cells; # = P<0.05, comparing to shCtrl cells with no drug treatment. **E-F,** Mean migration speed of shCtrl, shYAP-1, or -2 JHGBM651 cells treated with increasing doses of the STAT3 inhibitor LLL12. Mean migration speed is calculated from 30 cells. Results are representative of 3 independent experiments. Wilcoxon Rank-Sum test; Black * = P<0.05, comparing to no drug treatment in shCtrl cells; Red * = P<0.05, comparing to no drug treatment in shYAP cells; Red # = P<0.05, between shYAP and shCtrl cells at corresponding drug concentration. **G-H,** Representative immunoblots of TRIO, total and phospho-Tyr^705^ STAT3 protein levels in shCtrl, shTRIO-1, or -2 JHGBM1A, NHA, and MCF10A cells of 3 independent experiments. **I,** Representative immunoblots of total and phospho-Tyr^705^ STAT3 protein levels in JHGBM612 cells treated with increasing doses of the TRIO inhibitor ITX3 or Rac1-GEF inhibitor NSC23766 of 3 independent experiments. **J,** Representative immunoblots of total and phospho-Tyr^705^ STAT3 protein levels in JHGBM651/612 cells treated with increasing doses of the Rac1-GEF inhibitor NSC23766 of 3 independent experiments. **A, D-F,** Data shown is mean ± SEM.

**Figure S11.**
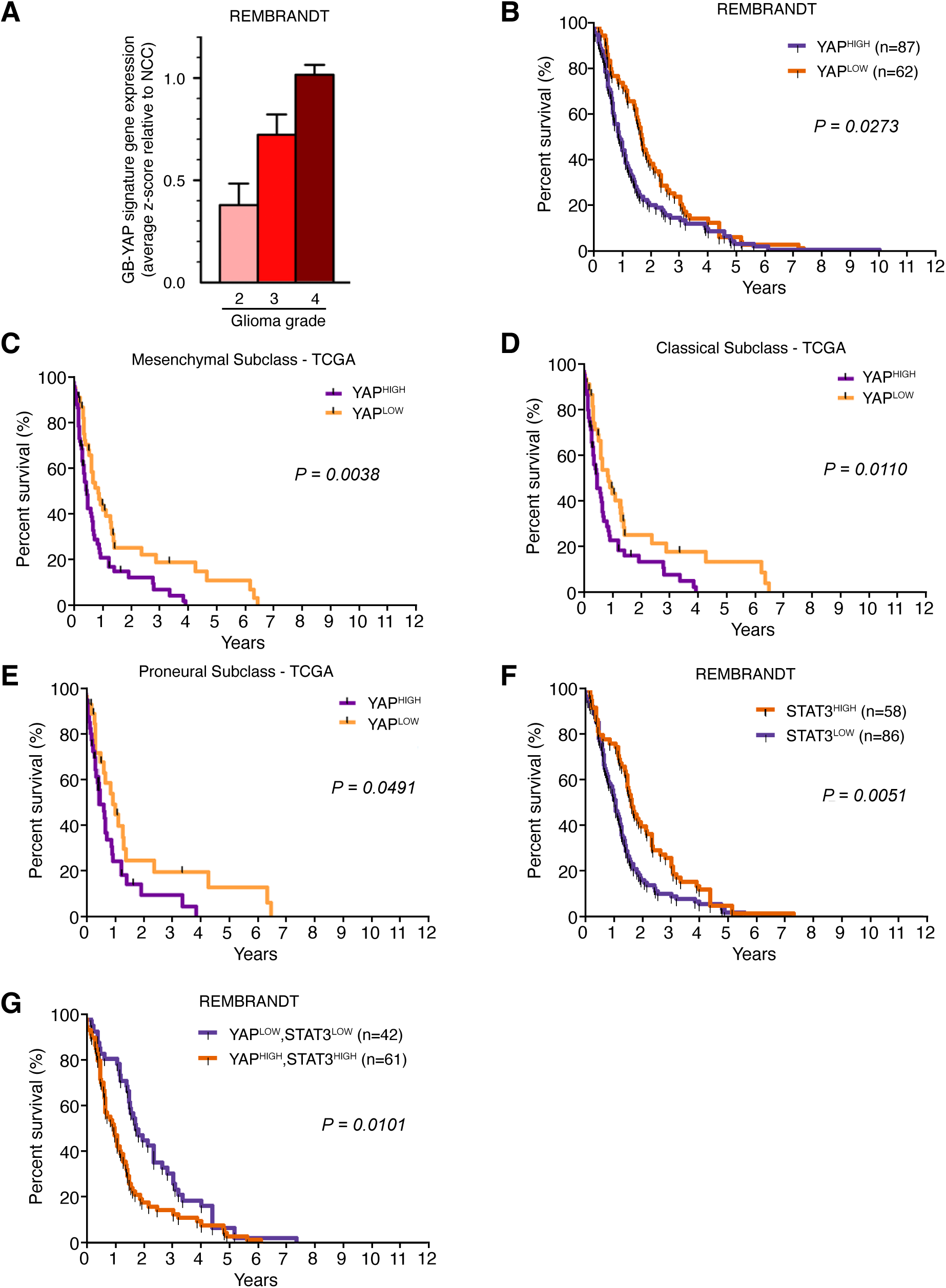
Elevated YAP signaling predicts poor clinical outcomes in all GBM subtypes. **A,** Expression of GB-YAP signature across different glioma grades in REMBRANDT dataset. Data shown is mean ± SEM. **B,** Kaplan-Meier graph of cumulative overall survival in patient groups defined by GB-YAP signature expression in REMBRANDT dataset. P = Log-Rank test. **C-E,** Kaplan-Meier graphs of cumulative overall survival in patient groups defined by GB-YAP signature expression in each indicated GBM subclass in the TCGA dataset. P=Log-Rank test. **F,** Kaplan-Meier graph of cumulative overall survival in patient groups defined by GB-STAT3 signature expression in REMBRANDT dataset. P=Log-Rank test. **G,** Kaplan-Meier graph of cumulative overall survival in patient groups defined by both the GB-YAP and GB-STAT3 signature expression in the REMBRANDT dataset. P = Log-Rank test.

**Figure S12.**
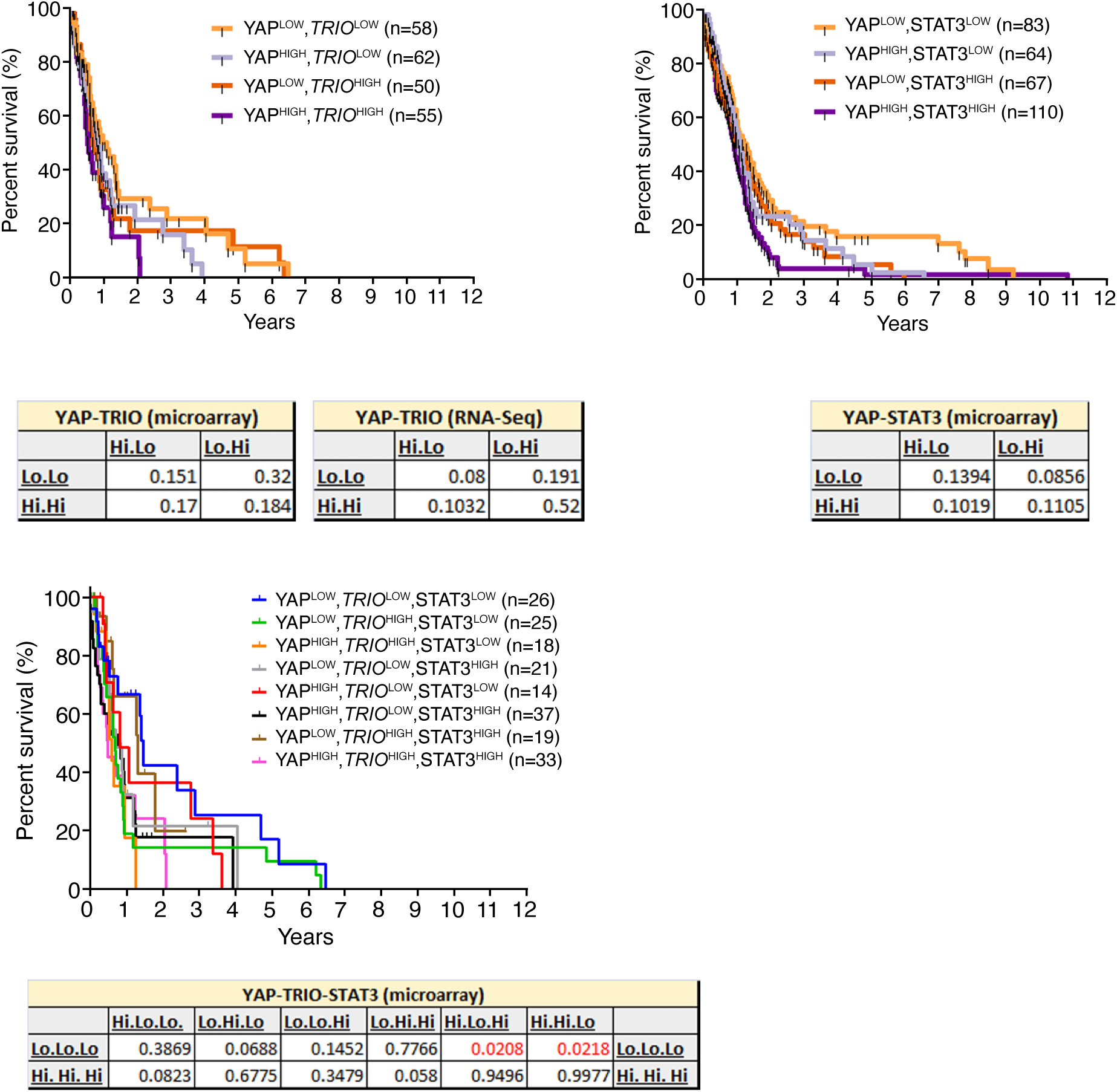
Each component of the YAP-TRIO-STAT3 network is indispensable to promoting poor clinical outcomes in GBM patients. Kaplan-Meir graphs of all combinations of mixed expression groups of YAP, *TRIO*, STAT3 in TCGA microarray dataset. Tables include p-value of comparisons between different combination groups. YAP-TRIO survival combinations were also evaluated in RNA-seq expression data from TCGA (table shown). P = Log-Rank test.

**Movie S1: Migration of shCtrl and shYAP GBM cells.** A representative time-lapse video of shCtrl (left) and shYAP (right) JHGBM651 cells. Images captured at 20-minute intervals for 6 hours. Each colored line represents the migration pattern of each cell. Scale bar = 100 μm. Movie is from a representative of 3 independent experiments.

**Movie S2. Migration of shCtrl and shYAP GBM cells with vehicle or ROCK inhibitor treatment.** A representative time-lapse video of shCtrl (left) and shYAP (right) JHGBM651 cells with vehicle (top) or treatment with the ROCK inhibitor Y27632 (bottom). Images are captured at 20-minute intervals for 6 hours. Each colored line represents the migration pattern of each cell. Scale bar = 100 μm. Movie is from a representative of 3 independent experiments.

**Movie S3. Migration of shCtrl and shYAP GBM cells with vehicle or Rac1-GEF inhibitor treatment.** A representative time-lapse video of shCtrl (left) and shYAP (right) JHGBM651 cells with vehicle (top) or treatment with the Rac1-GEF inhibitor NSC23766 (bottom). Images are captured at 20-minute intervals for 6 hours. Each colored line represents the migration pattern of each cell. Scale bar = 100 μm. Movie is from a representative of 3 independent experiments.

**Movie S4. Migration of shCtrl and shYAP GBM cells with vehicle or TRIO inhibitor treatment.** A representative time-lapse video of shCtrl (left) and shYAP (right) JHGBM651 cells with vehicle (top) or treatment with TRIO inhibitor ITX3 (bottom). Images are captured at 20-minute intervals for 6 hours. Each colored line represents the migration pattern of each cell. Scale bar = 100 μm. Movie is from a representative of 3 independent experiments.

**Movie S5. Migration of shCtrl and shTRIO GBM cells.** A representative time-lapse video of shCtrl (left) and shTRIO (right) JHGBM651 cells. Images captured at 20-minute intervals for 6 hours. Each colored line represents the migration pattern of each cell. Scale bar = 100 μm. Movie is from a representative of 3 independent experiments.

**Movie S6. Migration of shCtrl and shTRIO GBM cells with vehicle or ROCK inhibitor treatment.** A representative time-lapse video of shCtrl (left) and shTRIO (right) JHGBM651 cells with vehicle (top) or treatment with ROCK inhibitor Y27632 (bottom). Images are captured at 20-minute intervals for 6 hours. Each colored line represents the migration pattern of each cell. Scale bar = 100 μm. Movie is from a representative of 3 independent experiments.

**Movie S7. Migration of shCtrl and shYAP GBM cells with vehicle or STAT3 inhibitor treatment.** A representative time-lapse video of shCtrl (left) and shYAP (right) JHGBM651 cells with vehicle (top) or treatment with STAT3 inhibitor LLL12 (bottom). Images are captured at 20-minute intervals for 6 hours. Each colored line represents the migration pattern of each cell. Scale bar = 100 μm. Movie is from a representative of 3 independent experiments.

**Table S1:** Patient samples. Annotation of intraoperatively obtained human GBM tissues and cells with corresponding clinical characteristics of donor patient. This table contains GBM tissue and cell ID numbers with their corresponding de-identified patient IDs, age, gender, and final pathology diagnosis.

| Sample # | Identifier | Sex | Age | Pathology |
| --- | --- | --- | --- | --- |
| 1 | JHNC693 | F | 57 | Normal neocortex (meningioma; WHO grade I) |
| 2 | JHNC574 | F | 53 | Normal neocortex (mesial temporal sclerosis; hippocampus) |
| 3 | JHNC557 | M | 29 | Normal neocortex (mesial temporal sclerosis; hippocampus) |
| 4 | JHNC533 | M | 49 | Normal neocortex (mesial temporal sclerosis; hippocampus) |
| 5 | JHNC503 | M | 28 | Normal neocortex (mesial temporal sclerosis; hippocampus) |
| 6 | JHNC482 | F | 31 | Normal neocortex (mesial temporal sclerosis; hippocampus) |
| 7 | JHNC399 | F | 48 | Normal neocortex (mesial temporal sclerosis; hippocampus) |
| 8 | JHGB698 | F | 76 | Glioblastoma |
| 9 | JHGB697 | F | 60 | Glioblastoma (recurrent) |
| 10 | JHGB695 | F | 63 | Glioblastoma |
| 11 | JHGB692 | M | 56 | Glioblastoma |
| 12 | JHGB661 | M | 41 | Glioblastoma (primitive neuroectodermal + oligodendroglial features) |
| 13 | JHGB658 | F | 58 | Glioblastoma |
| 14 | JHGB653 | M | 74 | Glioblastoma |
| 15 | JHGB652 | M | 34 | Glioblastoma |
| 16 | JHGB651 | M | 49 | Glioblastoma |
| 17 | JHGB620 | F | 73 | Glioblastoma |
| 18 | JHGB615 | F | 76 | Glioblastoma |
| 19 | JHGB609 | M | 42 | Glioblastoma |
| 20 | JHGB608 | M | 69 | Glioblastoma |
| 21 | JHGB579 | M | 73 | Glioblastoma |
| 22 | JHGB577 | F | 57 | Glioblastoma |
| 23 | JHGB576 | M | 80 | Glioblastoma |
| 24 | JHGB567 | M | 48 | Glioblastoma |
| 25 | JHGB549 | F | 83 | Glioblastoma |
| 26 | JHGB547 | M | 64 | Glioblastoma (recurrent) |
| 27 | JHGB544 | M | 70 | Glioblastoma |
| 28 | JHGB529 | M | 86 | Glioblastoma |
| 29 | JHGB640 | F | 49 | Glioblastoma |
| 30 | JHGB612 | F | 56 | Glioblastoma |
| 31 | JHGB531 | F | 77 | Glioblastoma |
| 32 | JHGB496 | F | 53 | Glioblastoma (recurrent, secondary gliosarcoma) |

**Table S2.** shRNA plasmids. TRC (The RNAi Consortium) clone identifier numbers from Sigma-Aldrich for all shRNA constructs used in this study.

| Target Gene | Company | TRC Number | TRC Version |
| --- | --- | --- | --- |
| Human *YAP* clone 1 | Sigma-Aldrich | TRCN0000107265 | TRC 1 |
| Human *YAP* clone 2 | Sigma-Aldrich | TRCN0000107267 | TRC 1 |
| Mouse *Yap*/ Human *YAP* clone 3 | Sigma-Aldrich | TRCN0000107266 | TRC 1 |
| Human *YAP* clone 4 | Sigma-Aldrich | TRCN0000107268 | TRC 1 |
| Human *YAP* clone 5 | Sigma-Aldrich | TRCN0000107269 | TRC 1 |
| Human *TRIO* clone 1 | Sigma-Aldrich | TRCN0000195292 | TRC 1 |
| Human *TRIO* clone 2 | Sigma-Aldrich | TRCN0000196250 | TRC 1 |
| Human *TRIO* clone 3 | Sigma-Aldrich | TRCN0000196751 | TRC 1 |

**Table S3.**
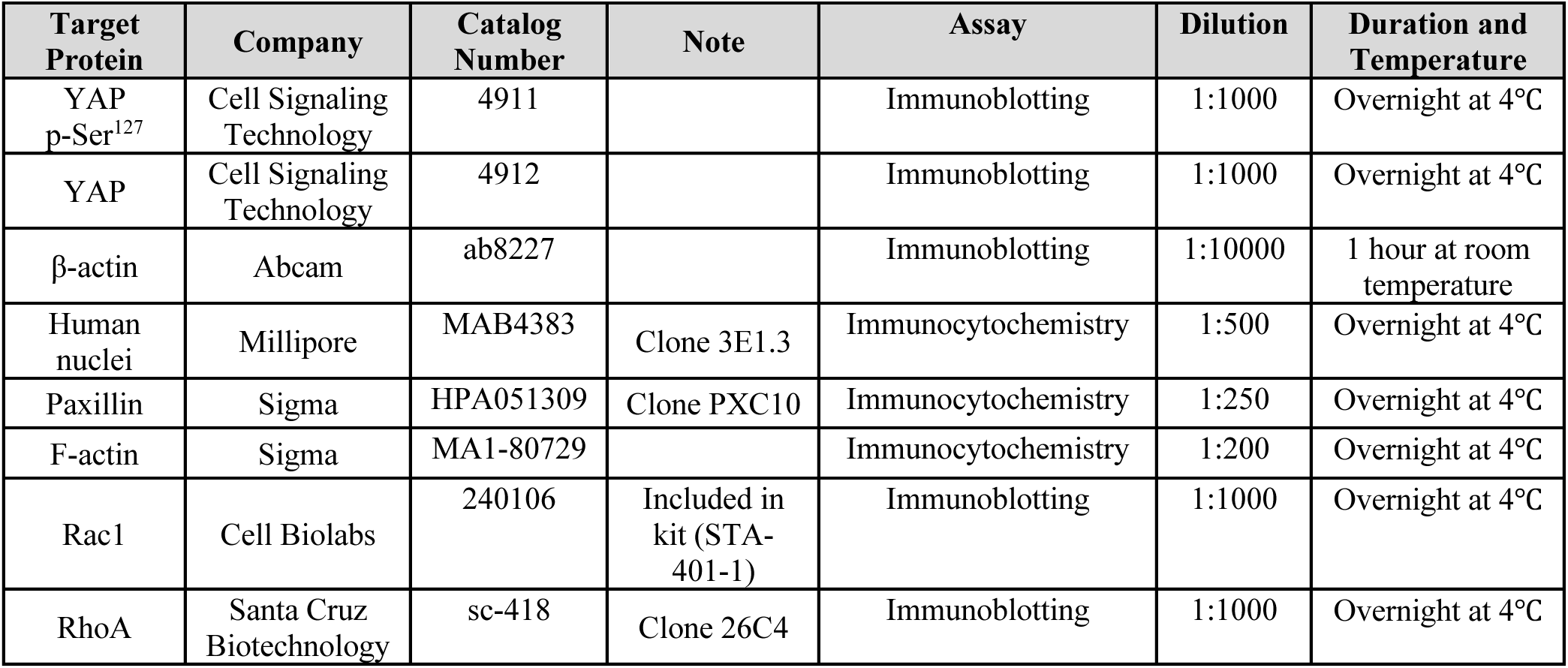
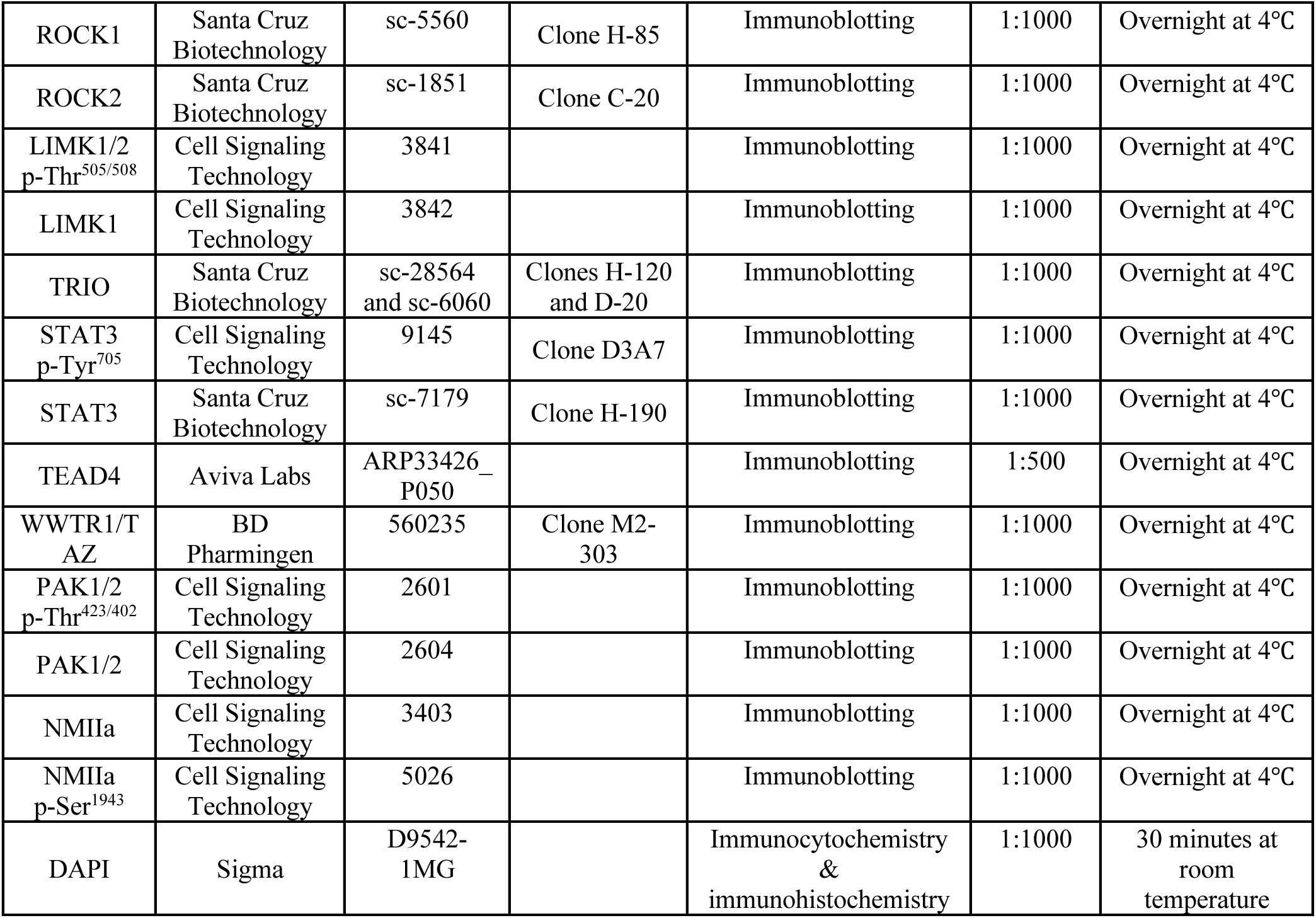
Antibodies. The information of supplier, catalog number, dilution ratio, incubation duration and temperature for all antibodies used in this study.

**Table S4.**
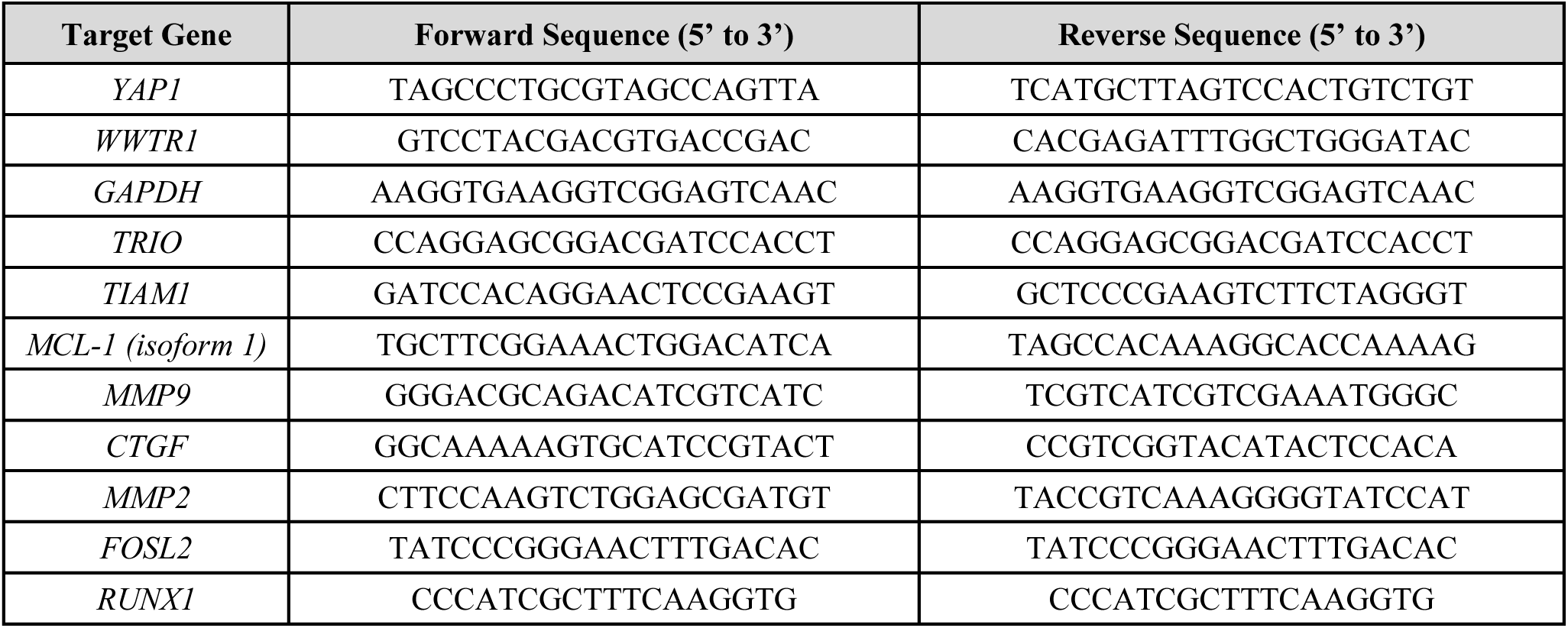
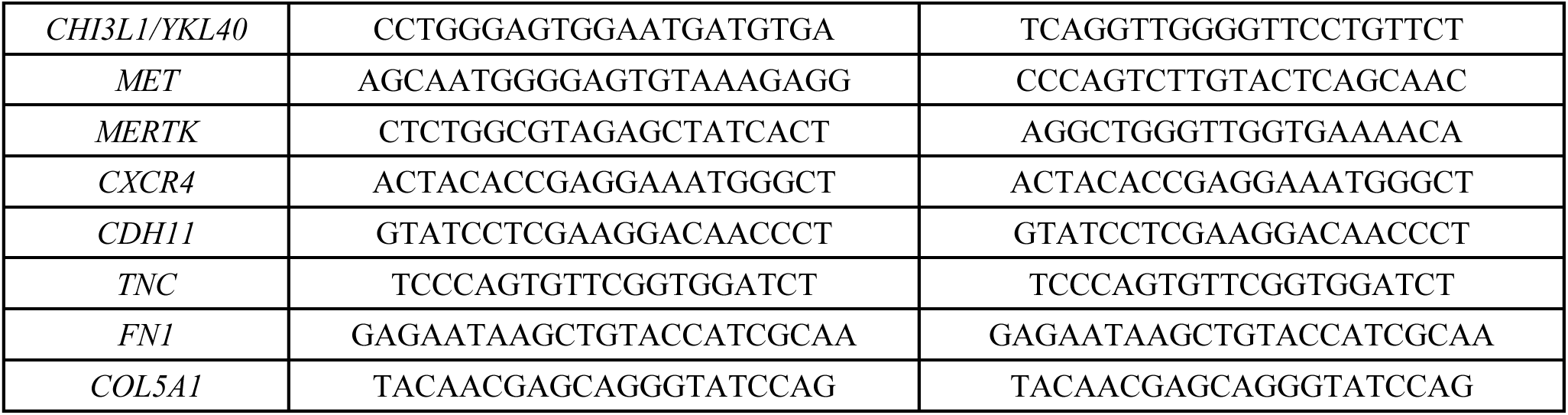
Primer pairs. Sequences for all primer pairs used in the quantitative real-time PCR experiments.

**Data File S1:** Microarray data of shCtrl or shYAP JHGBM651 and JHGBM612 cells.

**Data File S2:** Binding motif sequence of TEAD and TEAD4 binding motifs on *TRIO* enhancer sites.

## REFERENCES AND NOTES

1. E. Sahai, C. J. Marshall, RHO-GTPases and cancer. Nat Rev Cancer 2, 133–142 (2002).

2. F. Calvo, N. Ege, A. Grande-Garcia, S. Hooper, R. P. Jenkins, S. I. Chaudhry, K. Harrington, P. Williamson, E. Moeendarbary, G. Charras, E. Sahai, Mechanotransduction and YAP-dependent matrix remodelling is required for the generation and maintenance of cancer-associated fibroblasts. Nat Cell Biol 15, 637–646 (2013).

3. D. Chen, Y. Sun, Y. Wei, P. Zhang, A. H. Rezaeian, J. Teruya-Feldstein, S. Gupta, H. Liang, H.-K. Lin, M.-C. Hung, L. Ma, LIFR is a breast cancer metastasis suppressor upstream of the Hippo-YAP pathway and a prognostic marker. Nat Med 18, 1511–1517 (2012).

4. J. Dong, G. Feldmann, J. Huang, S. Wu, N. Zhang, S. A. Comerford, M. F. Gayyed, R. A. Anders, A. Maitra, D. Pan, Elucidation of a universal size-control mechanism in Drosophila and mammals. Cell 130, 1120–1133 (2007).

5. S. Dupont, L. Morsut, M. Aragona, E. Enzo, S. Giulitti, M. Cordenonsi, F. Zanconato, J. Le Digabel, M. Forcato, S. Bicciato, N. Elvassore, S. Piccolo, Role of YAP/TAZ in mechanotransduction. Nature 474, 179– 183 (2011).

6. J. M. Lamar, P. Stern, H. Liu, J. W. Schindler, Z.-G. Jiang, R. O. Hynes, The Hippo pathway target, YAP, promotes metastasis through its TEAD-interaction domain. Proceedings of the National Academy of Sciences 109, E2441–E2450 (2012).

7. M. Overholtzer, J. Zhang, G. A. Smolen, B. Muir, W. Li, D. C. Sgroi, C.-X. Deng, J. S. Brugge, D. A. Haber, Transforming properties of YAP, a candidate oncogene on the chromosome 11q22 amplicon. Proc Natl Acad Sci U S A 103, 12405–12410 (2006).

8. S. R. Shah, J. M. David, N. D. Tippens, A. Mohyeldin, J. C. Martinez-Gutierrez, S. Ganaha, P. Schiapparelli, D. H. Hamilton, C. Palena, A. Levchenko, A. Quiñones-Hinojosa, Brachyury-YAP Regulatory Axis Drives Stemness and Growth in Cancer. Cell Rep 21, 495–507 (2017).

9. S. Wei, J. Wang, O. Oyinlade, D. Ma, S. Wang, L. Kratz, B. Lal, Q. Xu, S. Liu, S. R. Shah, H. Zhang, Y. Li, A. Quiñones-Hinojosa, H. Zhu, Z.-Y. Huang, L. Cheng, J. Qian, S. Xia, Heterozygous IDH1R132H/WT created by “single base editing” inhibits human astroglial cell growth by downregulating YAP. Oncogene 37, 5160–5174 (2018).

10. S. A. Mosaddad, Y. Salari, S. Amookhteh, R. S. Soufdoost, Response to Mechanical Cues by Interplay of YAP/TAZ Transcription Factors and Key Mechanical Checkpoints of the Cell:A Comprehensive Review. Cell Physiol Biochem 55, 33–60 (2021).

11. S. Piccolo, T. Panciera, P. Contessotto, M. Cordenonsi, YAP/TAZ as master regulators in cancer: modulation, function and therapeutic approaches. Nat Cancer, 1–18 (2022).

12. O. Kilic, A. Yoon, S. R. Shah, H. M. Yong, A. Ruiz-Valls, H. Chang, R. A. Panettieri, S. B. Liggett, A. Quiñones-Hinojosa, S. S. An, A. Levchenko, A microphysiological model of the bronchial airways reveals the interplay of mechanical and biochemical signals in bronchospasm. Nat Biomed Eng 3, 532–544 (2019).

13. B. A. Orr, H. Bai, Y. Odia, D. Jain, R. A. Anders, C. G. Eberhart, Yes-associated protein 1 is widely expressed in human brain tumors and promotes glioblastoma growth. J Neuropathol Exp Neurol 70, 568– 577 (2011).

14. D. Pan, The hippo signaling pathway in development and cancer. Dev Cell 19, 491–505 (2010).

15. Y. Xu, I. Stamenkovic, Q. Yu, CD44 attenuates activation of the Hippo signaling pathway and is a prime therapeutic target for glioblastoma. Cancer Res 70, 2455–2464 (2010).

16. S. R. Shah, A. Quinones-Hinojosa, S. Xia, Advances in Brain Cancer: Creating Monoallelic Single Point Mutation in IDH1 by Single Base Editing. J Oncol Res Ther 5, 166 (2019).

17. L. Li, J. Luo, J.-Y. Fang, R. Zhang, J.-B. Ma, Z.-P. Zhu, Expression characteristics of the yes-associated protein in breast cancer: A meta-analysis. Medicine (Baltimore) 101, e30176 (2022).

18. X. Hu, Y. Zhang, H. Yu, Y. Zhao, X. Sun, Q. Li, Y. Wang, The role of YAP1 in survival prediction, immune modulation, and drug response: A pan-cancer perspective. Front Immunol 13, 1012173 (2022).

19. M. Castellan, A. Guarnieri, A. Fujimura, F. Zanconato, G. Battilana, T. Panciera, H. L. Sladitschek, P. Contessotto, A. Citron, A. Grilli, O. Romano, S. Bicciato, M. Fassan, E. Porcù, A. Rosato, M. Cordenonsi, S. Piccolo, Single-cell analyses reveal YAP/TAZ as regulators of stemness and cell plasticity in glioblastoma. Nat Cancer 2, 174–188 (2021).

20. S. Schmidt, A. Debant, Function and regulation of the Rho guanine nucleotide exchange factor Trio. Small GTPases 5, e29769 (2014).

21. E. Bell, L. J. Karnosh, Cerebral hemispherectomy; report of a case 10 years after operation. J Neurosurg 6, 285–293 (1949).

22. K. L. Chaichana, P. Zadnik, J. D. Weingart, A. Olivi, G. L. Gallia, J. Blakeley, M. Lim, H. Brem, A. Quiñones-Hinojosa, Multiple resections for patients with glioblastoma: prolonging survival. J Neurosurg 118, 812–820 (2013).

23. W. E. Dandy, REMOVAL OF RIGHT CEREBRAL HEMISPHERE FOR CERTAIN TUMORS WITH HEMIPLEGIA: PRELIMINARY REPORT. Journal of the American Medical Association 90, 823–825 (1928).

24. Y. Matsukado, C. S. Maccarty, J. W. Kernohan, The growth of glioblastoma multiforme (astrocytomas, grades 3 and 4) in neurosurgical practice. J Neurosurg 18, 636–644 (1961).

25. J. Lee, S. Kotliarova, Y. Kotliarov, A. Li, Q. Su, N. M. Donin, S. Pastorino, B. W. Purow, N. Christopher, W. Zhang, J. K. Park, H. A. Fine, Tumor stem cells derived from glioblastomas cultured in bFGF and EGF more closely mirror the phenotype and genotype of primary tumors than do serum-cultured cell lines. Cancer Cell 9, 391–403 (2006).

26. S. M. Pollard, K. Yoshikawa, I. D. Clarke, D. Danovi, S. Stricker, R. Russell, J. Bayani, R. Head, M. Lee, M. Bernstein, J. A. Squire, A. Smith, P. Dirks, Glioma stem cell lines expanded in adherent culture have tumor-specific phenotypes and are suitable for chemical and genetic screens. Cell Stem Cell 4, 568–580 (2009).

27. A. J. Ridley, A. Hall, The small GTP-binding protein rho regulates the assembly of focal adhesions and actin stress fibers in response to growth factors. Cell 70, 389–399 (1992).

28. M. Parri, P. Chiarugi, Rac and Rho GTPases in cancer cell motility control. Cell Communication and Signaling 8, 23 (2010).

29. G. A. Wildenberg, M. R. Dohn, R. H. Carnahan, M. A. Davis, N. A. Lobdell, J. Settleman, A. B. Reynolds, p120-catenin and p190RhoGAP regulate cell-cell adhesion by coordinating antagonism between Rac and Rho. Cell 127, 1027–1039 (2006).

30. L. K. Nguyen, B. N. Kholodenko, A. von Kriegsheim, Rac1 and RhoA: Networks, loops and bistability. Small GTPases 9, 316–321 (2018).

31. J. L. Bos, H. Rehmann, A. Wittinghofer, GEFs and GAPs: Critical Elements in the Control of Small G Proteins. Cell 129, 865–877 (2007).

32. Y. Gao, J. B. Dickerson, F. Guo, J. Zheng, Y. Zheng, Rational design and characterization of a Rac GTPase-specific small molecule inhibitor. Proc Natl Acad Sci U S A 101, 7618–7623 (2004).

33. F. Zanconato, M. Forcato, G. Battilana, L. Azzolin, E. Quaranta, B. Bodega, A. Rosato, S. Bicciato, M. Cordenonsi, S. Piccolo, Genome-wide association between YAP/TAZ/TEAD and AP-1 at enhancers drives oncogenic growth. Nat Cell Biol 17, 1218–1227 (2015).

34. N. D. Heintzman, R. K. Stuart, G. Hon, Y. Fu, C. W. Ching, R. D. Hawkins, L. O. Barrera, S. Van Calcar, C. Qu, K. A. Ching, W. Wang, Z. Weng, R. D. Green, G. E. Crawford, B. Ren, Distinct and predictive chromatin signatures of transcriptional promoters and enhancers in the human genome. Nat Genet 39, 311– 318 (2007).

35. A. Sethi, M. Gu, E. Gumusgoz, L. Chan, K.-K. Yan, J. Rozowsky, I. Barozzi, V. Afzal, J. A. Akiyama, I. Plajzer-Frick, C. Yan, C. S. Novak, M. Kato, T. H. Garvin, Q. Pham, A. Harrington, B. J. Mannion, E. A. Lee, Y. Fukuda-Yuzawa, A. Visel, D. E. Dickel, K. Y. Yip, R. Sutton, L. A. Pennacchio, M. Gerstein, Supervised enhancer prediction with epigenetic pattern recognition and targeted validation. Nat Methods 17, 807–814 (2020).

36. F. Jin, Y. Li, J. R. Dixon, S. Selvaraj, Z. Ye, A. Y. Lee, C.-A. Yen, A. D. Schmitt, C. A. Espinoza, B. Ren, A high-resolution map of the three-dimensional chromatin interactome in human cells. Nature 503, 290–294 (2013).

37. I. Hazan, J. Monin, B. A. M. Bouwman, N. Crosetto, R. I. Aqeilan, Activation of Oncogenic Super-Enhancers Is Coupled with DNA Repair by RAD51. Cell Reports 29, 560–572.e4 (2019).

38. J. A. Castro-Mondragon, R. Riudavets-Puig, I. Rauluseviciute, R. Berhanu Lemma, L. Turchi, R. Blanc-Mathieu, J. Lucas, P. Boddie, A. Khan, N. Manosalva Pérez, O. Fornes, T. Y. Leung, A. Aguirre, F. Hammal, D. Schmelter, D. Baranasic, B. Ballester, A. Sandelin, B. Lenhard, K. Vandepoele, W. W. Wasserman, F. Parcy, A. Mathelier, JASPAR 2022: the 9th release of the open-access database of transcription factor binding profiles. Nucleic Acids Research 50, D165–D173 (2022).

39. J.-W. Jang, M.-K. Kim, Y.-S. Lee, J.-W. Lee, D.-M. Kim, S.-H. Song, J.-Y. Lee, B.-Y. Choi, B. Min, X.-Z. Chi, S.-C. Bae, RAC-LATS1/2 signaling regulates YAP activity by switching between the YAP-binding partners TEAD4 and RUNX3. Oncogene 36, 999–1011 (2017).

40. M. Cordenonsi, F. Zanconato, L. Azzolin, M. Forcato, A. Rosato, C. Frasson, M. Inui, M. Montagner, A. R. Parenti, A. Poletti, M. G. Daidone, S. Dupont, G. Basso, S. Bicciato, S. Piccolo, The Hippo Transducer TAZ Confers Cancer Stem Cell-Related Traits on Breast Cancer Cells. Cell 147, 759–772 (2011).

41. N. Bouquier, E. Vignal, S. Charrasse, M. Weill, S. Schmidt, J.-P. Léonetti, A. Blangy, P. Fort, A cell active chemical GEF inhibitor selectively targets the Trio/RhoG/Rac1 signaling pathway. Chem Biol 16, 657–666 (2009).

42. G. Salloum, L. Jaafar, M. El-Sibai, Rho A and Rac1: Antagonists moving forward. Tissue and Cell 65, 101364 (2020).

43. K. M. Byrne, N. Monsefi, J. C. Dawson, A. Degasperi, J.-C. Bukowski-Wills, N. Volinsky, M. Dobrzyński, M. R. Birtwistle, M. A. Tsyganov, A. Kiyatkin, K. Kida, A. J. Finch, N. O. Carragher, W. Kolch, L. K. Nguyen, A. von Kriegsheim, B. N. Kholodenko, Bistability in the Rac1, PAK, and RhoA Signaling Network Drives Actin Cytoskeleton Dynamics and Cell Motility Switches. Cell Syst 2, 38–48 (2016).

44. R. Liu, X. Wang, G. Y. Chen, P. Dalerba, A. Gurney, T. Hoey, G. Sherlock, J. Lewicki, K. Shedden, M. F. Clarke, The Prognostic Role of a Gene Signature from Tumorigenic Breast-Cancer Cells. New England Journal of Medicine 356, 217–226 (2007).

45. H. Yu, D. Pardoll, R. Jove, STATs in cancer inflammation and immunity: a leading role for STAT3. Nat Rev Cancer 9, 798–809 (2009).

46. J. Corry, H. R. Mott, D. Owen, Activation of STAT transcription factors by the Rho-family GTPases. Biochemical Society Transactions 48, 2213–2227 (2020).

47. A. R. Simon, H. G. Vikis, S. Stewart, B. L. Fanburg, B. H. Cochran, K.-L. Guan, Regulation of STAT3 by Direct Binding to the Rac1 GTPase. Science 290, 144–147 (2000).

48. K. Zhou, J. Rao, Z. Zhou, X. Yao, F. Wu, J. Yang, L. Yang, X. Zhang, Y. Cui, X.-W. Bian, Y. Shi, Y. Ping, RAC1-GTP promotes epithelial-mesenchymal transition and invasion of colorectal cancer by activation of STAT3. Laboratory Investigation 98, 989–998 (2018).

49. T. R. Faruqi, D. Gomez, X. R. Bustelo, D. Bar-Sagi, N. C. Reich, Rac1 mediates STAT3 activation by autocrine IL-6. Proc Natl Acad Sci U S A 98, 9014–9019 (2001).

50. E. L. Morgan, A. Macdonald, Autocrine STAT3 activation in HPV positive cervical cancer through a virus-driven Rac1—NFκB—IL-6 signalling axis. PLOS Pathogens 15, e1007835 (2019).

51. H. Bagci, M. Laurin, J. Huber, W. J. Muller, J.-F. Côté, Impaired cell death and mammary gland involution in the absence of Dock1 and Rac1 signaling. Cell Death Dis 5, e1375–e1375 (2014).

52. H. Kim, J. K. Sonn, Rac1 promotes chondrogenesis by regulating STAT3 signaling pathway. Cell Biology International 40, 976–983 (2016).

53. L. Zhao, X. Du, K. Huang, T. Zhang, Z. Teng, W. Niu, C. Wang, G. Xia, Rac1 modulates the formation of primordial follicles by facilitating STAT3-directed Jagged1, GDF9 and BMP15 transcription in mice. Sci Rep 6, 23972 (2016).

54. G. Liu, Y. Xia, H. Wang, X. Jin, S. Chen, W. Chen, C. Zhang, Y. He, Downregulation of CYRI-B promotes migration, invasion and EMT by activating the Rac1-STAT3 pathway in gastric cancer. Experimental Cell Research 423, 113453 (2023).

55. M. S. Carro, W. K. Lim, M. J. Alvarez, R. J. Bollo, X. Zhao, E. Y. Snyder, E. P. Sulman, S. L. Anne, F. Doetsch, H. Colman, A. Lasorella, K. Aldape, A. Califano, A. Iavarone, The transcriptional network for mesenchymal transformation of brain tumours. Nature 463, 318–325 (2010).

56. Q. Wang, B. Hu, X. Hu, H. Kim, M. Squatrito, L. Scarpace, A. C. deCarvalho, S. Lyu, P. Li, Y. Li, F. Barthel, H. J. Cho, Y.-H. Lin, N. Satani, E. Martinez-Ledesma, S. Zheng, E. Chang, C.-E. G. Sauvé, A. Olar, Z. D. Lan, G. Finocchiaro, J. J. Phillips, M. S. Berger, K. R. Gabrusiewicz, G. Wang, E. Eskilsson, J. Hu, T. Mikkelsen, R. A. DePinho, F. Muller, A. B. Heimberger, E. P. Sulman, D.-H. Nam, R. G. W. Verhaak, Tumor Evolution of Glioma-Intrinsic Gene Expression Subtypes Associates with Immunological Changes in the Microenvironment. Cancer Cell 32, 42–56.e6 (2017).

57. H. S. Phillips, S. Kharbanda, R. Chen, W. F. Forrest, R. H. Soriano, T. D. Wu, A. Misra, J. M. Nigro, H. Colman, L. Soroceanu, P. M. Williams, Z. Modrusan, B. G. Feuerstein, K. Aldape, Molecular subclasses of high-grade glioma predict prognosis, delineate a pattern of disease progression, and resemble stages in neurogenesis. Cancer Cell 9, 157–173 (2006).

58. T. M. Malta, C. F. de Souza, T. S. Sabedot, T. C. Silva, M. S. Mosella, S. N. Kalkanis, J. Snyder, A. V. B. Castro, H. Noushmehr, Glioma CpG island methylator phenotype (G-CIMP): biological and clinical implications. Neuro Oncol 20, 608–620 (2018).

59. B. Salhia, N. L. Tran, A. Chan, A. Wolf, M. Nakada, F. Rutka, M. Ennis, W. S. McDonough, M. E. Berens, M. Symons, J. T. Rutka, The guanine nucleotide exchange factors trio, Ect2, and Vav3 mediate the invasive behavior of glioblastoma. Am J Pathol 173, 1828–1838 (2008).

60. M. Sakabe, J. Fan, Y. Odaka, N. Liu, A. Hassan, X. Duan, P. Stump, L. Byerly, M. Donaldson, J. Hao, M. Fruttiger, Q. R. Lu, Y. Zheng, R. A. Lang, M. Xin, YAP/TAZ-CDC42 signaling regulates vascular tip cell migration. Proceedings of the National Academy of Sciences 114, 10918–10923 (2017).

61. J. P. Vaqué, R. T. Dorsam, X. Feng, R. Iglesias-Bartolome, D. J. Forsthoefel, Q. Chen, A. Debant, M. A. Seeger, B. R. Ksander, H. Teramoto, J. S. Gutkind, A genome-wide RNAi screen reveals a Trio-regulated Rho GTPase circuitry transducing mitogenic signals initiated by G protein-coupled receptors. Mol Cell 49, 94–108 (2013).

62. J. Cai, X. Song, W. Wang, T. Watnick, Y. Pei, F. Qian, D. Pan, A RhoA-YAP-c-Myc signaling axis promotes the development of polycystic kidney disease. Genes Dev 32, 781–793 (2018).

63. X. Feng, M. S. Degese, R. Iglesias-Bartolome, J. P. Vaque, A. A. Molinolo, M. Rodrigues, M. R. Zaidi, B. R. Ksander, G. Merlino, A. Sodhi, Q. Chen, J. S. Gutkind, Hippo-independent activation of YAP by the GNAQ uveal melanoma oncogene through a trio-regulated rho GTPase signaling circuitry. Cancer Cell 25, 831–845 (2014).

64. R. Gargini, M. Escoll, E. García, R. García-Escudero, F. Wandosell, I. M. Antón, WIP Drives Tumor Progression through YAP/TAZ-Dependent Autonomous Cell Growth. Cell Rep 17, 1962–1977 (2016).

65. H. Sabra, M. Brunner, V. Mandati, B. Wehrle-Haller, D. Lallemand, A.-S. Ribba, G. Chevalier, P. Guardiola, M. R. Block, D. Bouvard, β1 integrin-dependent Rac/group I PAK signaling mediates YAP activation of Yes-associated protein 1 (YAP1) via NF2/merlin. J Biol Chem 292, 19179–19197 (2017).

66. F.-X. Yu, J. Luo, J.-S. Mo, G. Liu, Y. C. Kim, Z. Meng, L. Zhao, G. Peyman, H. Ouyang, W. Jiang, J. Zhao, X. Chen, L. Zhang, C.-Y. Wang, B. C. Bastian, K. Zhang, K.-L. Guan, Mutant Gq/11 promote uveal melanoma tumorigenesis by activating YAP. Cancer Cell 25, 822–830 (2014).

67. P. Tocci, R. Cianfrocca, V. Di Castro, L. Rosanò, A. Sacconi, S. Donzelli, S. Bonfiglio, G. Bucci, E. Vizza, G. Ferrandina, G. Scambia, G. Tonon, G. Blandino, A. Bagnato, β-arrestin1/YAP/mutant p53 complexes orchestrate the endothelin A receptor signaling in high-grade serous ovarian cancer. Nat Commun 10, 3196 (2019).

68. Y. Qiao, J. Chen, Y. B. Lim, M. L. Finch-Edmondson, V. P. Seshachalam, L. Qin, T. Jiang, B. C. Low, H. Singh, C. T. Lim, M. Sudol, YAP Regulates Actin Dynamics through ARHGAP29 and Promotes Metastasis. Cell Reports 19, 1495–1502 (2017).

69. J. Shen, Q. Huang, W. Jia, S. Feng, L. Liu, X. Li, D. Tao, D. Xie, YAP1 induces invadopodia formation by transcriptionally activating TIAM1 through enhancer in breast cancer. Oncogene 41, 3830–3845 (2022).

70. J. W. Haskins, D. X. Nguyen, D. F. Stern, Neuregulin 1-activated ERBB4 interacts with YAP to induce Hippo pathway target genes and promote cell migration. Sci Signal 7, ra116 (2014).

71. J. He, Q. Bao, Y. Zhang, M. Liu, H. Lv, Y. Liu, L. Yao, B. Li, C. Zhang, S. He, G. Zhai, Y. Zhu, X. Liu, K. Zhang, X.-J. Wang, M.-H. Zou, Y. Zhu, D. Ai, Yes-Associated Protein Promotes Angiogenesis via Signal Transducer and Activator of Transcription 3 in Endothelial Cells. Circ Res 122, 591–605 (2018).

72. J. Park, D. Eisenbarth, W. Choi, H. Kim, C. Choi, D. Lee, D.-S. Lim, YAP and AP-1 Cooperate to Initiate Pancreatic Cancer Development from Ductal Cells in Mice. Cancer Research 80, 4768–4779 (2020).

73. Y. Liu, C. Xu, J. Li, Y. Zhang, X. Wang, Y. Wang, J. Qin, Z. Zheng, Y. Xia, YAP promotes AP-1 expression in tubular epithelial cells in the kidney. American Journal of Physiology-Renal Physiology 324, F581–F589 (2023).

74. L. He, H. Pratt, M. Gao, F. Wei, Z. Weng, K. Struhl, YAP and TAZ are transcriptional co-activators of AP-1 proteins and STAT3 during breast cellular transformation. eLife 10, e67312 (2021).

75. J. Park, D.-H. Kim, S. R. Shah, H.-N. Kim, Kshitiz, P. Kim, A. Quiñones-Hinojosa, A. Levchenko, Switch-like enhancement of epithelial-mesenchymal transition by YAP through feedback regulation of WT1 and Rho-family GTPases. Nat Commun 10, 2797 (2019).

76. D. Hanahan, R. A. Weinberg, Hallmarks of Cancer: The Next Generation. Cell 144, 646–674 (2011).

77. B. S. Wong, S. R. Shah, C. L. Yankaskas, V. K. Bajpai, P.-H. Wu, D. Chin, B. Ifemembi, K. ReFaey, P. Schiapparelli, X. Zheng, S. S. Martin, C.-M. Fan, A. Quiñones-Hinojosa, K. Konstantopoulos, A microfluidic cell-migration assay for the prediction of progression-free survival and recurrence time of patients with glioblastoma. Nat Biomed Eng 5, 26–40 (2021).

78. J. Kim, J. G. Shamul, S. R. Shah, A. Shin, B. J. Lee, A. Quinones-Hinojosa, J. J. Green, Verteporfin-Loaded Poly(ethylene glycol)-Poly(beta-amino ester)-Poly(ethylene glycol) Triblock Micelles for Cancer Therapy. Biomacromolecules 19, 3361–3370 (2018).

79. J. G. Shamul, S. R. Shah, J. Kim, P. Schiapparelli, C. A. Vazquez-Ramos, B. J. Lee, K. K. Patel, A. Shin, A. Quinones-Hinojosa, J. J. Green, Verteporfin-Loaded Anisotropic Poly(Beta-Amino Ester)-Based Micelles Demonstrate Brain Cancer-Selective Cytotoxicity and Enhanced Pharmacokinetics. Int J Nanomedicine 14, 10047–10060 (2019).

80. S. R. Shah, J. Kim, P. Schiapparelli, C. A. Vazquez-Ramos, J. C. Martinez-Gutierrez, A. Ruiz-Valls, K. Inman, J. G. Shamul, J. J. Green, A. Quinones-Hinojosa, Verteporfin-Loaded Polymeric Microparticles for Intratumoral Treatment of Brain Cancer. Mol Pharm 16, 1433–1443 (2019).

81. X. Wang, B. Wu, Z. Zhong, Downregulation of YAP inhibits proliferation, invasion and increases cisplatin sensitivity in human hepatocellular carcinoma cells. Oncol Lett 16, 585–593 (2018).

82. Z. Zhou, J.-S. Zhu, C.-P. Gao, L.-P. Li, C. Zhou, H. Wang, X.-G. Liu, siRNA targeting YAP gene inhibits gastric carcinoma growth and tumor metastasis in SCID mice. Oncol Lett 11, 2806–2814 (2016).

83. J. Wang, L. Yuan, X. Xu, Z. Zhang, Y. Ma, L. Hong, J. Ma, Rho-GEF Trio regulates osteosarcoma progression and osteogenic differentiation through Rac1 and RhoA. Cell Death Dis 12, 1148 (2021).

84. O. Gonzalez-Perez, H. Guerrero-Cazares, A. Quiñones-Hinojosa, Targeting of deep brain structures with microinjections for delivery of drugs, viral vectors, or cell transplants. J Vis Exp, 2082 (2010).

85. X. Xie, P. Rigor, P. Baldi, MotifMap: a human genome-wide map of candidate regulatory motif sites. Bioinformatics 25, 167–174 (2009).

86. M. Dai, P. Wang, A. D. Boyd, G. Kostov, B. Athey, E. G. Jones, W. E. Bunney, R. M. Myers, T. P. Speed, H. Akil, S. J. Watson, F. Meng, Evolving gene/transcript definitions significantly alter the interpretation of GeneChip data. Nucleic Acids Res 33, e175 (2005).

87. R. A. Irizarry, C. Wang, Y. Zhou, T. P. Speed, Gene set enrichment analysis made simple. Stat Methods Med Res 18, 565–575 (2009).

